# Major stages of vertebrate adaptive radiation are assembled from a disparate spatiotemporal landscape

**DOI:** 10.1101/2020.03.12.988774

**Authors:** Emilie J. Richards, Joseph A. McGirr, Jeremy R. Wang, Michelle E. St. John, Jelmer W. Poelstra, Maria J. Solano, Delaney C. O’Connell, Bruce J. Turner, Christopher H. Martin

## Abstract

To investigate the origins and stages of vertebrate adaptive radiation, we reconstructed the spatial and temporal histories of genetic variants underlying major phenotypic axes of diversification from the genomes of 202 Caribbean pupfishes. Ancient standing variation from disparate spatial sources was reassembled into new combinations which are under strong selection for adaptation to novel trophic niches on only a single island throughout the Caribbean. This occurred in three stages: first, standing variation associated with feeding behavior swept, then standing variation regulating craniofacial development and pigmentation, and finally de novo variation for craniofacial development. Our results provide clear support for two longstanding hypotheses about adaptive radiation and demonstrate how ancient alleles maintained for millennia in distinct environmental refugia can be assembled into new adaptive combinations.

**One Sentence Summary:** Ancient origins of adaptive radiation

## Main Text

Adaptive radiations are fundamental to understanding the biodiversity of life. These bursts of phenotypic and ecological diversification may occur in response to ecological opportunity provided by unoccupied niche space (*1, 2*). However, the origins and major features of this process are still controversial. For example, ecological opportunity does not explain why only some lineages radiate (*3–6*). One hypothesis is that introgression from disparate source populations might be necessary to trigger diversification (*7, 8*). Despite substantial evidence of adaptive introgression during radiation (*9–12*), no previous studies have compared adaptive introgression between closely related radiating and non-radiating lineages to distinguish introgression as a trigger. A parallel debate centers on whether adaptive diversification proceeds in three distinct temporal stages: shifts in habitat preference first, followed by trophic morphology, and finally sexual communication (*13*). This ‘behavior-first’ view is also commonly proposed for initiating adaptive trait evolution (*14–16*). However, existing evidence for the temporal stages hypothesis comes from ancestral state reconstructions of rapidly diversifying traits which are unreliable without fossil data (*17–19*).

Here we provide strong support for these two major hypotheses of adaptive radiation using multiple lines of genomic, transcriptomic, and phenotypic evidence in a nascent adaptive radiation of Caribbean pupfishes. This sympatric radiation contains a widespread generalist algae and detritus-eating species (*Cyprinodon variegatus*) and two trophic specialists endemic to 10 ky old hypersaline lakes on San Salvador Island (SSI), Bahamas: a molluscivore *C. brontotheroides* with a unique nasal protrusion and a scale-eater *C. desquamator* with striking two-fold longer oral jaws. This clade exhibits hallmarks of adaptive radiation. First trait diversification rates are up to 1,400 times faster than non-radiating generalist populations on neighboring Bahamian islands in nearly identical hypersaline lake environments. Second, craniofacial diversity within the radiation is comparable to all other Cyprinodontidae species combined (*5, 20*).

To investigate the spatiotemporal history of adaptive variants unique to trophic specialists on SSI we first constructed a high quality *de novo* hybrid assembly for *C. brontotheroides* (1.16 Gb size; scaffold N50 = 32 Mb; L50 = 15; 86.4% complete Actinopterygii BUSCOs) and resequenced 202 genomes (7.9x median coverage) from across the range of *Cyprinodon* and the two closest outgroups *Megupsilon* and *Cualac* (Fig. 1A;Table S1; Data S1). Population structure across the Caribbean was largely explained by geographic distance (Fig. 1) and the SSI radiation did not contain higher overall genetic diversity than the rest of the Caribbean (Fig. S1). All Caribbean populations experienced similar declines in effective population size following the last glacial maximum 15 kya when an order of magnitude more Caribbean coastal habitat was above sea level (Fig. 1D).

**Fig. 1.**
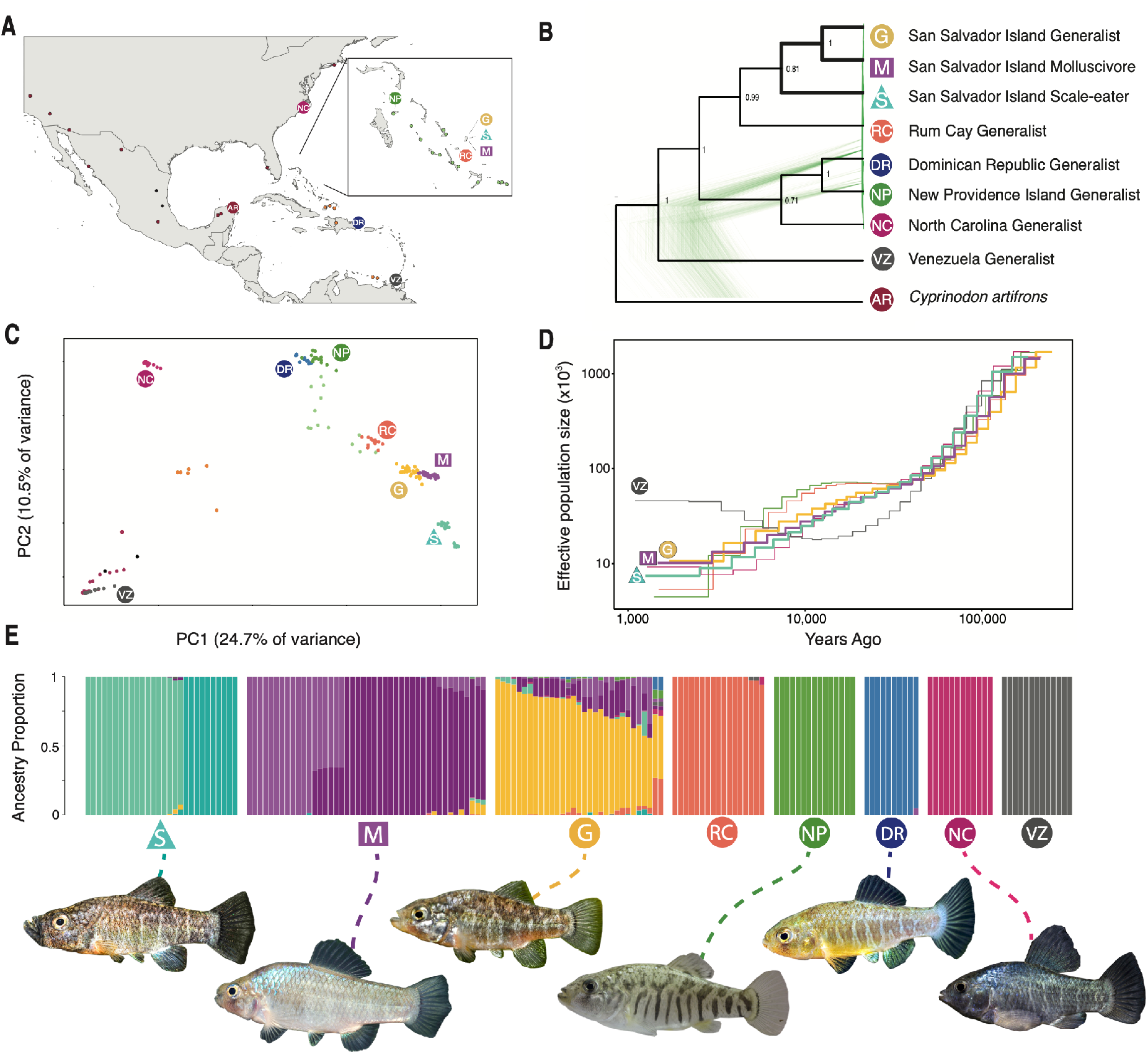
Genetic diversity of pupfishes across the Caribbean. A) Sample locations of *Cyprinodon* pupfishes (*n* = 202) with eight focal populations (*n* ≥ 10 per population) marked with symbols and single individuals from other locations in the Bahamas (light green), Caribbean (orange), continental North and South America (maroon), and *Megupsilon* and *Cualac* outgroups to *Cyprinodon* (black). B) Maximum clade credibility phylogeny estimated with SNAPP from 10K SNPs for focal populations and the outgroup *Cyprinodon artifrons overlaying 500 gene trees randomly sampled from the posterior distribution and visualized with Densitree*. C) Principal component analysis of *Cyprinodon* pupfishes. D) Changes in effective population size over time for focal populations in the Caribbean inferred using MSMC. E) Ancestry proportions across individuals in San Salvador and 5 other Caribbean populations. Proportions estimated from a LD-pruned dataset in ADMIXTURE with k=11.

We scanned 5.5 million variants across 202 Caribbean-wide pupfish genomes to identify 3,464 scale-eater and 1,491 molluscivore candidate adaptive variants, respectively, showing evidence of both strong genetic differentiation between trophic specialists (*F_st_* ≥ 0.95) and a hard selective sweep (Fig. 2A; Fig S2; Data S2 and S3). 28% of these candidate adaptive variants were found in cis-regulatory regions, 12% in intronic regions, and 2% in coding regions, resulting in a total of 176 candidate genes for adaptive radiation (Table S2–S3). These genes were enriched (FDR < 0.05) for developmental processes associated with the major axes of phenotypic diversification in this radiation, including feeding behavior, muscle tissue, and craniofacial development (Fig. 2C; Table S4). 45% of these genes were also differentially expressed between trophic specialists (FDR < 0.05; Data S4-S5) in whole embryos at 2 and/or 8 days post fertilization (dpf) (*21*).

**Fig. 2.**
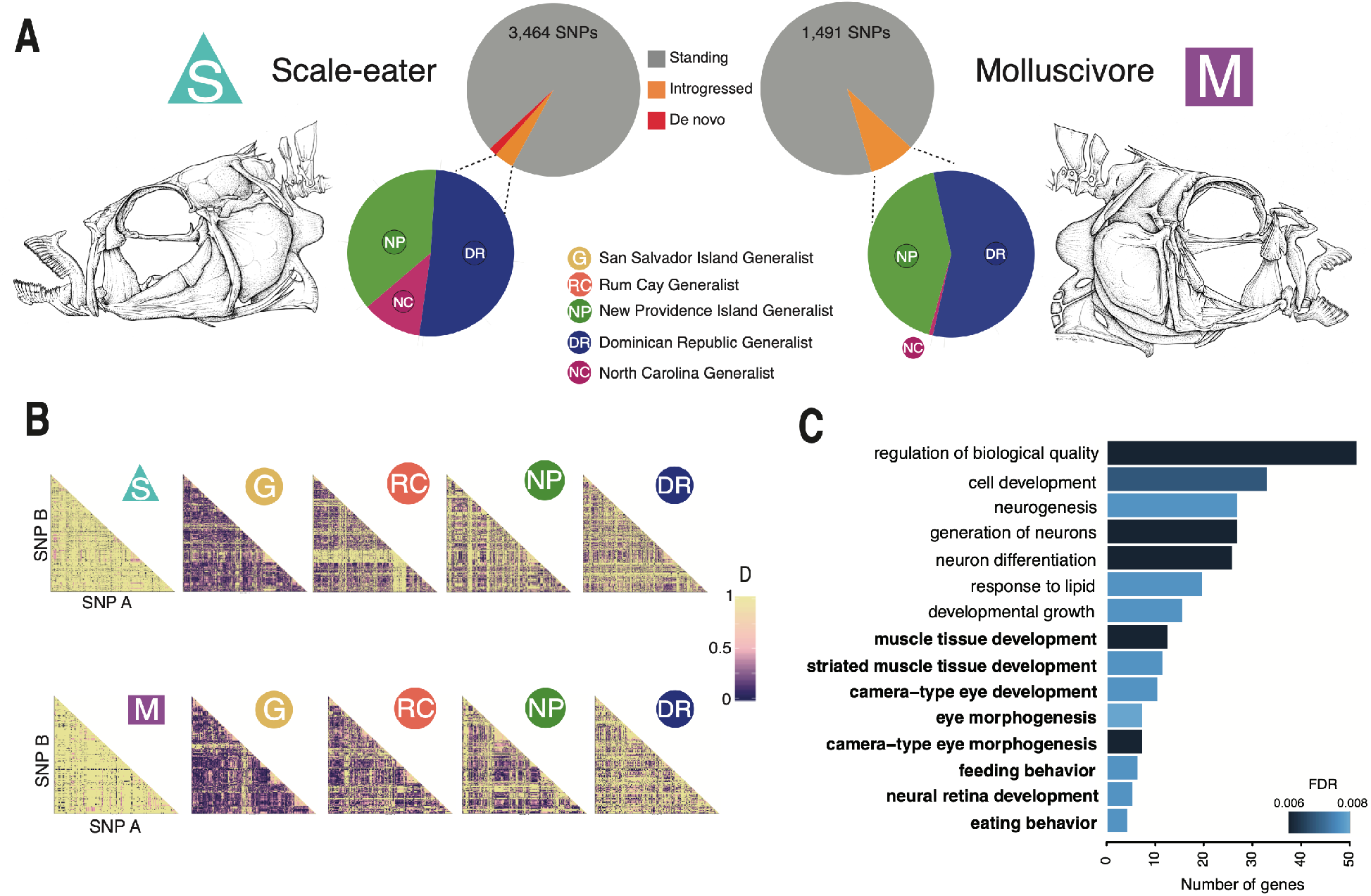
All candidate adaptive genetic variants underlying trophic specialist pupfishes. A) Origins of all candidate adaptive variants for trophic specialists (F_st_ > 0.95; hard selective sweep CLR≥ 4.47.; see supplementary methods for more details) divided into three categories: SSI only (de novo: red), introgression from a specific source population (introgressed: orange), or observed in multiple Caribbean populations as standing genetic variation (standing: grey). Introgressed variation is further broken down by focal source population (Tables S9–S12). B) Heatmaps of linkage disequilibrium among all pairwise combinations of candidate adaptive variants for scale-eaters (S; top row) and molluscivores (M; bottom row) on SSI in comparison to linkage patterns among these SNPSs in generalists on SSI (G) and three other focal generalist populations across the Caribbean (Venezuela removed from comparison due to its unique bottleneck history among the populations). Note the breakdown in linkage disequilibrium among candidate adaptive variants outside of trophic specialist populations. C) Top 15 GO categories in which scale-eater candidate adaptive variants were significantly enriched, with relevant terms corresponding to major axes of divergence in this radiation highlighted in bold (FDR < 0.01; full list of terms with FDR < 0.05 in Table S4).

Even though both trophic specialists are endemic to SSI, we also found nearly all their candidate adaptive variants to occur as standing genetic variation across the Caribbean (molluscivore: 100%; scale-eater: 98%; Fig. 2A). Furthermore, nearly half these variants were ancient and also found in *Cualac* or *Megupsilon* outgroups to *Cyprinodon* (41% and 55% of scale-eater and molluscivore adaptive variants, respectively), which diverged over 5 Mya (*22*). However, most adaptive variants did not show any evidence of selection in five other focal high-coverage Caribbean generalist populations (only 2% and 6% of scale-eater and molluscivore variants, respectively; Fig. S3) and strong linkage disequilibrium among adaptive variants in SSI trophic specialists was not observed in these focal populations (Fig. 2B). Thus, novel trophic specialists within a microendemic adaptive radiation were almost entirely assembled from ancient standing genetic variation through strong selection for new adaptive combinations of alleles.

Multiple lines of evidence suggest that more hybridization and adaptive introgression took place in SSI populations than other Caribbean island populations, consistent with the hypothesis that hybridization triggered adaptive radiation. First, the strongest signal of introgression across the Caribbean was into the root node of the SSI radiation (Fig. 3B). Second, trophic specialists on San Salvador experienced at least twice as much adaptive introgression as other generalist populations across the Caribbean (*P* < 0.006; Fig. 3C). Third, based on differences between the distribution of tract lengths for neutral and adaptively introgressed regions, we infer that adaptive introgression only resulted from older hybridization events with generalist source populations, despite evidence of recent and continuous introgression to the present (Fig 3D-E).

**Fig. 3.**
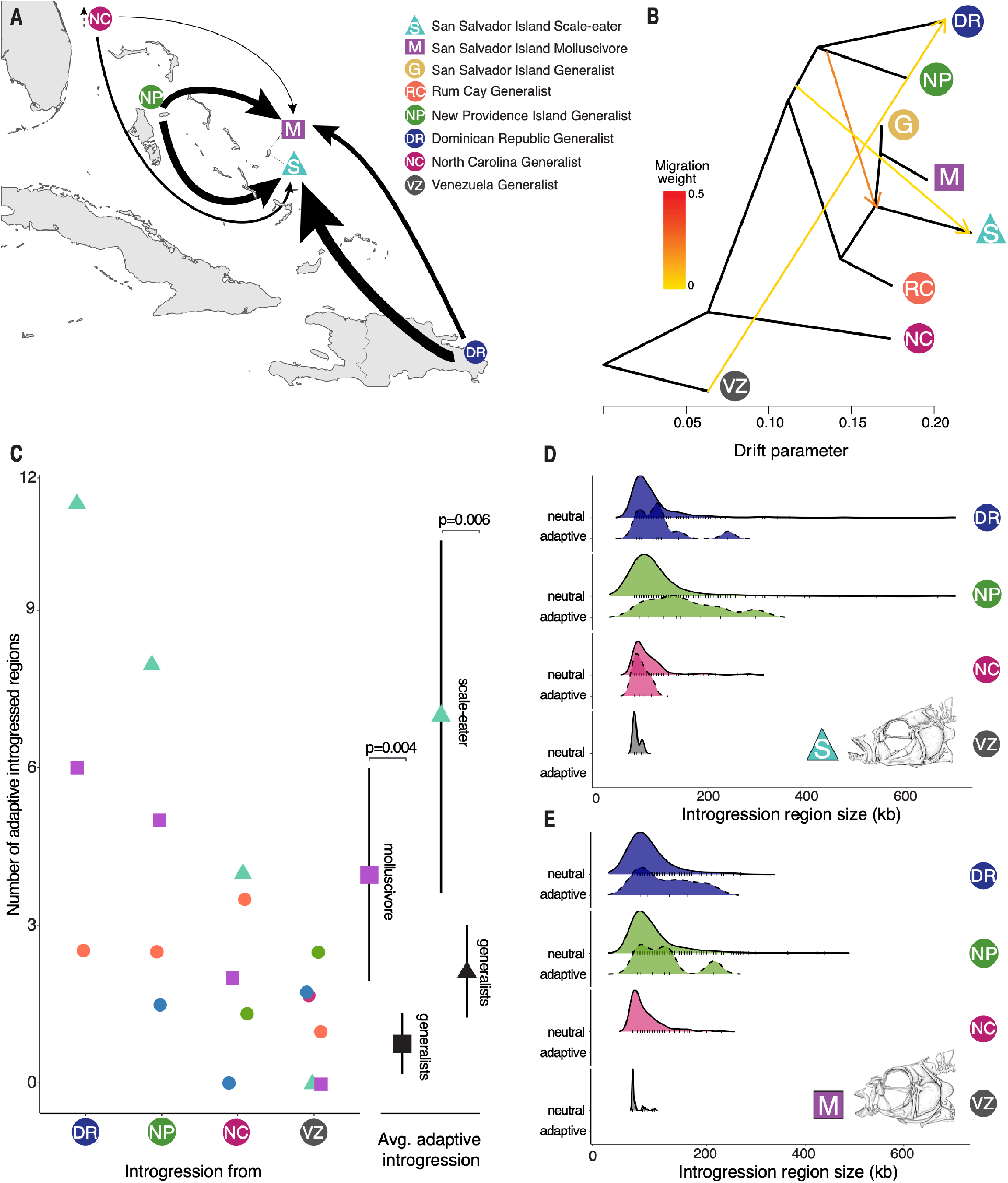
A history of hybridization across the Caribbean. A) A summary map of adaptive introgression into SSI trophic specialists from focal generalist populations across the Caribbean, with thickness of arrows proportional to the number of outlier regions of the *f_d_* statistic. B) Genome-wide population graph inferred from *Treemix* with the 3 strongest signals of admixture. Note that the strongest signal is into the root node of the SSI radiation. C) The total number of candidate adaptive introgression regions from outgroup generalist populations is often larger in specialists (triangle: scale-eater and square: molluscivore) than in other outgroup generalist populations (circle, color matches population legend in panel A). The bootstrapped mean and confidence interval for the number of adaptive introgression events into the molluscivore and scale-eater populations compared to other Caribbean generalist populations are shown to the right of the panel (triangle: scale-eater introgression regions and square: molluscivore introgression regions). D-E) Density plots of the size distribution of adaptive introgression regions (dashed line) and neutral introgression regions (solid line) across the genomes of D) scale-eaters and E) molluscivores, divided into the four focal generalist source populations. Tick marks on the bottom of density curves indicate the number of introgression regions observed.

We also observed distinct temporal stages of adaptive radiation based on divergence time estimates of candidate loci and timing of selective sweeps. SSI pupfish diversified along the three major axes of vertebrate adaptive radiation (*13*), including their trophic environment (mediated through foraging behavior: scale-eating or snail-eating), trophic morphology, and sexual communication signals. Based on both GO annotations of genes near candidate adaptive variants and genome-wide association mapping (GWAS) in Caribbean pupfishes for these trait axes, we found that the divergence times of loci associated with these three major stages of adaptive radiation exhibited a significant temporal signal overall (ANOVA, *P* = 0.03; Fig. 4A), driven by a ‘behavior-first’ pattern (permutation test; *P*=0.0041). Independent adaptive variants in the *cis*-regulatory regions of two genes associated with feeding behavior (*prlh*, *cfap20;* (*23, 24*)) were the oldest estimated sweeps out of all adaptive loci associated with these axes and swept from standing genetic variation ten times older than the radiation itself (Fig. 4A,C; 95% HPD sweep ages: 6,747-8,490 and 6,594-9,210 years, respectively). *Cfap20* and *prlh* were also both differentially expressed between trophic specialists at early developmental stages, consistent with a *cis*-regulatory function of these variants (Data S4-S5;(*21*)).

**Fig. 4.**
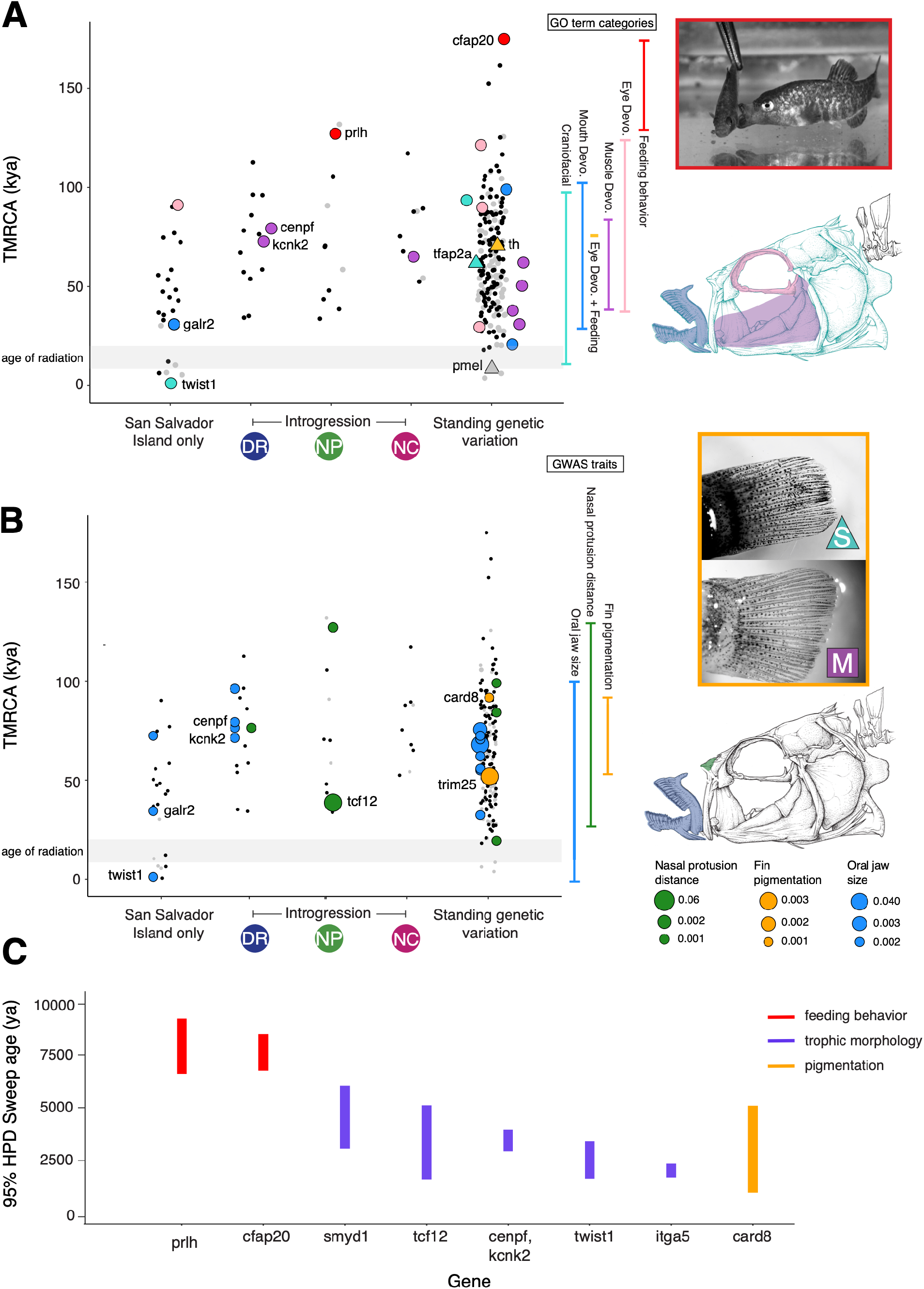
The spatiotemporal landscape of adaptive radiation in pupfish. Time to most recent common ancestor (TMRCA) of candidate adaptive variants based on *Dxy* for 50-kb region containing candidate adaptive variants. Variants are separated by spatial distribution: SSI only (de novo), introgression, and standing genetic variation. Gray lines highlight the approximate age of the radiation: from the peak of the last glacial maximum (approximately 20 kya; (*32*)) to the youngest age estimate for filling of hypersaline lakes on SSI (∼3 kya: (*33*)). A) Variants are colored by their adaptive relevance to this system. Black: adaptive variants annotated for non-focal GO terms or unannotated; Gray: nearly fixed neutral variants between specialists with no signal of hard selective sweep; and triangles represent variants associated with pigmentation. All variants annotated for the GO categories of feeding behavior (red), muscle (purple), eye, mouth development (blue), and craniofacial development (mouth and eye and/or muscle(teal)) are highlighted. Genes highlighted in the text are labeled by their associated variant. B) Variants are colored by significant association (99_th_ percentile of PIP) in GWAS with a larger lower oral jaw size, darker caudal fin pigmentation, and smaller nasal protrusion distance. Dot sizes scale with PIP score. C) 95% HPD interval of the posterior distribution for selective sweep ages for focal gene regions. Regions are colored by GO and/or GWAS annotations relevant to the three major stages of adaptive radiation: habitat preference (feeding behavior), trophic morphology (craniofacial and muscle), and sexual communication (pigmentation).

Consistent with the second stage of trophic morphological divergence, adaptive variants in the regulatory regions of genes associated with eye development (*gnat2, tbc1d20, zhx2*), muscle development (*fhod3, smyd1, kcnk2, fhl2, pdim5*), and craniofacial morphology (*bcor, itga5, tfap2a*, *med1*) swept from both standing genetic variation and adaptive introgression from three different focal populations across the Caribbean (Fig. 4). We also observed a final stage of novel refinement in which two *de novo* variants associated with craniofacial morphology arose on SSI and swept in scale-eating populations (Fig 4). One of these variants is a non-synonymous substitution in the second exon of *twist1*, a transcription factor known to influence cleft palate and oral jaw size (*25*), which was significantly associated with SSI pupfish oral jaw size in a genome-wide association scan (99^th^ PIP percentile GWAS; Fig 4B), is highly conserved across ray-finned fishes (GERP score: 2.19; Fig S4), and is one of the most recent sweeps of any adaptive variant detected in our analyses (95% HPD: 1,636-3,413 ya). The second variant is in the regulatory region upstream of the gene *galr2*, which produces a transmembrane receptor for galanin, a peptide known to facilitate bone formation (*26*). This gene was significantly associated with SSI pupfish oral jaw size (99^th^ PIP percentile GWAS; Fig 4B; Data S6) and lies within a significant QTL that accounts for 15% of the variation in oral jaw size in an F2 intercross between SSI specialist species (Table S5; (*27*)). Molluscivores displayed similar temporal stages with craniofacial variation sweeping recently (Fig. S5), but contained no *de novo* variants. 85% of candidate adaptive regions under selection in molluscivores were also under selection in scale-eaters but contained different fixed variants (Fig. S3), consistent with previously observed patterns of parallel metabolic pathway gene expression in trophic specialists but divergent genotypes (*21*).

There was no evidence for a temporally distinct third stage of diversification in sexual signals, despite the striking divergence in male reproductive pigmentation between SSI trophic specialists. Instead, pigmentation diverged throughout the process of adaptive radiation. Adaptive variants in two genes known to affect pigmentation (*tfap2a*, *th*; Fig 4A) and two additional candidate variants associated with pupfish caudal fin pigmentation (99th PIP percentile GWAS; Fig. 4B; Data S7) indicate that diversification in male reproductive coloration spans a range of divergence times, with some variants more ancient than the radiation itself (Fig 4B). For example, two nearly fixed adaptive variants in the regulatory region of *card8*, a homolog of *nlrp1* which is associated with vitiligo and pigmentation loss in humans [(*28*); 99^th^ PIP percentile GWAS; differentially expressed at 2 and 8 dpf; Fig 4B; Data S4-S5,S7], swept at the same time as many variants associated with craniofacial morphology during the second stage of adaptive radiation. Divergence estimates of loci associated with pigmentation also span a large range: from variants in the regulatory region of *card8*, which is ten times older than the radiation, to a single variant fixed in scale-eaters and in the neighboring Rum Cay generalist population found in the regulatory region of *pmel* (known to affect melanosome development (*29*); differentially expressed between specialists at 8 dpf; Data S5), which is as young as the radiation itself (Fig. 4A). This broad range of divergence times and sweep ages may indicate that the distinctive light/dark reproductive coloration associated with trophic specialists diverged throughout the process of adaptive radiation using existing standing genetic variation, rather than only during a final stage.

Intriguingly, along with distinct temporal patterns of adaptive divergence, we also find distinct spatial patterns to the sources of adaptive introgression. Introgression from different regions of the Caribbean brought in adaptive variation for different major axes of phenotypic diversification within the radiation. Adaptive variants associated with feeding behavior (*prlh;* Fig 4A) and nasal protrusion (*tcf12*; 99^th^ PIP percentile GWAS;Fig 4B;Data S8) originated in the northwestern Bahamas (New Providence Island, Exumas, and Cat Island) whereas adaptive introgression of variants associated with muscle development arrived from the southeastern Caribbean in the Dominican Republic (*cenpf, eya2;* Fig 4A). This suggests that the extant SSI radiation of trophic specialists was reassembled from pools of ancient genetic variation contained in at least two distinct environmental refugia in other regions of the Caribbean, perhaps due to previous ephemeral adaptive radiations within these regions, thus connecting micro- and macroevolutionary-scale processes (*30, 31*).

In conclusion, the SSI adaptive radiation was triggered by the formation of a hybrid swarm of largely ancient alleles maintained within different pools of standing variation in Caribbean and mainland outgroups. Distinct temporal stages of adaptation observed in this nascent radiation are consistent with a behavior-first stage of vertebrate radiation. Much of the adaptive variation contributing to major phenotypic axes of diversification in this radiation originated over longer timescales preceding the divergence of trophic specialist species on SSI, and across large spatial scales. Our results show that adaptive radiations can occupy expansive evolutionary spaces: spanning the existing radiation itself and the multitude of both past and present ephemeral pools of genetic variation that contributed to rapid diversification. Research into the broader spatiotemporal landscape of vertebrate radiations, including the hominin radiation (*31*), can provide clear answers regarding longstanding hypotheses about their origins and contributions to global patterns of biodiversity.

## Acknowledgments

We thank Rebecca Tarvin for helpful comments on the manuscript; the Gerace Research Centre and Troy Day for logistical support; the governments of the Bahamas, Dominican Republic, the National Park Service and U.S. Fish and Wildlife Service for permission to collect and export samples; the Vincent J. Coates Genomics Sequencing Center and Functional Genomics Laboratory at UC Berkeley for performing whole-genome library preparation and sequencing (supported by NIH S10 OD018174 Instrumentation Grant), and the University of North Carolina at Chapel Hill for computational resources.

## Author contributions

Conceptualization – EJR,CHM; Data Curation: EJR,JRW,MJS, DCOC; Formal Analyses: EJR,JAM,MSJ; Resources: CHM, BJT; Visualization: EJR,CHM; Writing – original draft: EJR; Writing – review & editing: EJR, CHM, JAM, MSJ, JRW, JWP, MJS.

## Data and materials availability

Genomic and RNA sequence data are archived at the National Center for Biotechnology Information BioProject Database (Accession: XXX; PRJNA394148, PRJNA391309; and PRJNA305422); and additional data and scripts are on the Dryad Digital Repository (XXX) and Github (https://github.com/joemcgirr).

## Supplementary Materials

Materials and Methods

Figures S1-S13

Tables S1-S14

External Databases S1-S9

References (*34-86*)

## Materials and Methods

### Sampling

Pupfishes were collected from across the complete Atlantic and Caribbean range of *Cyprinodon* from Massachusetts to Venezuela. For the three species of the SSI radiation, individual pupfish were collected from 15 isolated hypersaline lakes on San Salvador Island (Table S1;Data S1) and one estuary (Pigeon Creek) using hand and seine nets between 2011 and 2018. 36 *Cyprinodon variegatus*, 47 *C. brontotheroides*, and 39 *C. desquamator* were sampled across these lakes, including six lakes in which one or two specialist species occur in sympatry with the generalists (Crescent Pond, Storr’s Lake, Little Lake, Oyster Pond, Osprey Lake, Moon Rock Pond). The sampling of outgroup high-coverage focal populations of generalist pupfish included 17 individuals from *C. laciniatus* from Lake Cunningham, New Providence Island, Bahamas; 18 *C. variegatus* from Lake George, Rum Cay, Bahamas; 12 *C. higuey* from Laguna Bavaro, Dominican Republic; 14 *C. variegatus* from Fort Fisher estuary, North Carolina, United States; and 14 *C. dearborni* from Isla Margarita, Venezuela. 37 individuals were also collected from other islands and localities spanning the range of *Cyprinodon* across the Caribbean and Atlantic coasts, including captive-bred individuals from the extinct species *Megupsilon aporus* and threatened species *Cualac tessellatus,* the most closely related outgroup genera to *Cyprinodon* (Fig. 1A; Table S1;Data S1;(*1*)).

Fishes were euthanized in an overdose of buffered MS-222 (Finquel, Inc.) following approved protocols from the University of California, Davis Institutional Animal Care and Use Committee (#17455), the University of North Carolina at Chapel Hill Animal Care and Use Committee (#18-061.0), and the University of California, Berkeley Animal Care and Use Committee (AUP-2015-01-7053) and preserved in 95-100% ethanol.

### Genomic library preparation

DNA was extracted from muscle tissue using DNeasy Blood and Tissue kits (Qiagen, Inc.) and quantified on a Qubit 3.0 fluorometer (Thermofisher Scientific, Inc.). Genomic libraries were prepared using the automated Apollo 324 system (WaterGen Biosystems, Inc.) at the Vincent J. Coates Genomic Sequencing Center (QB3). Samples were fragmented using Covaris sonication, barcoded with Illumina indices, and quality checked using a Fragment Analyzer (Advanced Analytical Technologies, Inc.). Nine to ten samples were pooled per lane for 150PE sequencing on four lanes of an Illumina Hiseq4000 and an additional 96 individuals were sequenced on one 150PE lane of Illumina Novaseq with S4 chemistry. This included 42 individuals from a previous genomic study (*2*).

### De novo genome assembly and annotation

We constructed a hybrid de novo assembly for an inbred lab-raised individual of *C. brontotheroides* using three different sequencing technologies. Oxford Nanopore sequencing was performed at UNC’s High Throughput Sequencing Facility, a 10X Genomics synthetic long-read library was prepared and sequenced by Hudson Alpha, and Chicago and HiC libraries were prepared and sequenced by Dovetail Genomics. Genomic DNA was extracted from an inbred F4 male *C. brontotheroides* individual, an offspring from three generations of full-sib mating in the lab, starting with an F0 generation collected from Crescent Pond, SSI. 10X sequencing was performed on this individual according to 10X Genomics’ recommended protocol, sequenced on a HiSeq4000, resulting in 460 million 2×150 bp reads. DNA was extracted from this same molluscivore individual for Nanopore sequencing using modified a phenol:chloroform extraction protocol (*3*). Two libraries were sequenced on R9.4 flow cells on GridION – one using the Rapid Sequencing Kit (RAD004) and one Ligation Kit (LSK109), producing 4.9 Gbp of sequences with a read length N50 of 4.7 Kbp.

10X Genomics sequences were first assembled using Supernova (v2.0.0, 10X Genomics) to produce a preliminary “pseudohap” assembly. Nanopore reads were corrected using FMLRC (*4*). The Supernova assembly was scaffolded with corrected nanopore reads using LINKS (*5*) using the recommended iterative approach (34 rounds). The nanopore-scaffolded assembly was further scaffolded using HiC and Chicago sequences; we predicted Hi-C contacts using Juicer (v1.6.2; (*6*)), followed by scaffolding with 3D-DNA (v180922) (*7*). We performed a final polishing with four rounds of Racon (v1.3.1; (*8*)) using corrected nanopore reads. The final assembly consists of 1.16 Gbp in 15,698 scaffolds with an N50 of 32,013,756 bp.

To further validate our assembly, we ran BUSCO (v3.0.1) (*9*) to identify known single-copy conserved genes. We found 86.4% of BUSCOs in the Actinopterygii class assembled completely, and 83.4% in a single copy. We annotated this assembly using the Maker pipeline (v3.01.02)(*10*), providing alternate ESTs and protein evidence for ab-initio gene prediction from *C. variegatus* (*11*), which is expected to have very similar genic structure and codon usage. Predicted genes were assigned putative function by aligning (BLASTp) to the UniProt database.

### Population genotyping

Raw reads were mapped from 222 individuals to a de-novo assembly of *Cyprinodon brontotheroides* reference genome (v 1.0; total sequence length = 1,162,855,435 bp; number of scaffold = 15,698, scaffold N50 = 32 Mbp) with bwa-mem (v 0.7.12;(*12*)). Duplicate reads were identified using MarkDuplicates and BAM indices were created using BuildBamIndex in the Picard software package (http://picard.sourceforge.net(v.2.0.1)). We followed the best practices guide recommended in the Genome Analysis Toolkit (v 3.5;(*13*)) to call and refine our single nucleotide polymorphism (SNP) variant dataset using the program HaplotypeCaller. We filtered SNPs based on the recommended hard filter criteria (i.e. QD < 2.0; FS < 60; MQRankSum < - 12.5; ReadPosRankSum < −8;(*13, 14*)) because we lacked high-quality known variants for these non-model species. Variants for San Salvador Island individuals were additionally filtered to remove SNPs with a minor allele frequency below 0.05, genotype quality below 20, or containing more than 20% missing data across all individuals at the site using vcftools (v.0.1.15;(*15*)). This set of 9.3 million variants was then further filtered for variants that had minor allele frequencies above 0.05 and less than 50% missing data across all Caribbean outgroup individuals with population level sampling. Variants in poorly mapped regions were then removed using a mask file generated from the program SNPable (http://bit.ly/snpable; k-mer length =50, and ‘stringency’=0.5). Our final dataset after filtering contained 5.5 million variants.

### Population genetic analyses

This dataset was first pruned for SNPs in strong linkage disequilibrium using the LD pruning function (--indep-pairwise 50 5 0.5) in plink (v1.9)(*16*), leaving 2.6 million variants. To visualize population structure in our dataset, we ran a principle component analysis using the eigenvectors outputted by plink’s pca function (--pca). The first two principal components were plotted in R (R Core Team 2018 v3.5.0). To visualize admixture among the species we estimated the proportion of shared ancestry among individuals in our dataset using ADMIXTURE (v.1.3.0)(*17*). The number of populations (K) was decided upon using ADMIXTURE’s cross-validation method (--cv) across 1-20 population values of K. A K of 11 populations was then chosen using the broken-stick method. Ancestry proportions estimated by ADMIXTURE were plotted in R. Four individuals that appeared to exhibit recent hybrid ancestry between *C. variegatus* and *C. brontotheroides* and two individuals that appeared to exhibit recent hybrid ancestry between *C. variegatus* and *C. desquamator* were removed from downstream analyses. We also excluded 15 individuals that appeared as strong outliers in the PCA and ADMIXTURE analyses (3 *C. variegatus* from San Salvador Island, 1 *C. brontotheroides*, 3 *C. laciniatus*, 2 *C. higuey*, 3 *C. variegatus* from North Carolina, and 3 *C. dearborni* from Venezuela), resulting in 32 *Cyprinodon variegatus*, 44 *C. brontotheroides*, and 26 *C. desquamator* individuals from San Salvador Island, 16 individuals from *C. laciniatus* from Lake Cunningham, New Providence Island in the Bahamas, 17 *C. variegatus* from Lake George, Rum Cay, 10 *C. higuey* from Lake Bavaro, Dominican Republic, 12 *C. variegatus* from Fort Fisher estuary North Carolina, and 11 *C. dearborni* from Isla Margarita, Venezuela (Fig 1E). None of the 37 single individuals from other locations were removed. The final dataset used in downstream analyses included 202 individuals.

Within-population nucleotide diversity (*π*) was calculated in 50-kb windows across the genome for each of eight focal populations with more than 10 individuals sampled. To ensure an equal comparison among populations, we downsampled individuals from each population to the number of individuals in the focal population with the lowest sampling (n=10). We did this by randomly selecting 10 individuals for each population before calculating *π* in sliding windows. We repeated this 100 times and averaged *π* across the replicates (Fig. S1). Due to the large sample size of windows for each population (n=30,762), slight differences in mean genome-wide within-population genetic diversity resulted in statistically significant differences in genome-wide diversity among populations (ANOVA, *P*> 2.2×10^−16^). However, the effect sizes of the difference in these means was small in all comparisons except San Salvador Island generalists and North Carolina and Venezuela generalist populations (Cohen’s d=0.87 and 1.38 respectively). The much lower within-population genetic diversity in Venezuela than other generalist populations could be explained by the fact that this population seems to have undergone a recent bottleneck in the past that was not observed in the other populations sampled (Fig. 1C and S1).

Genome-wide *F_st_* for pairwise San Salvador Island species comparisons was calculated for each variant site and in 50-kb windows with at least 100 variant sites across the genome using the python script popGenWindows.py available from https://github.com/simonhmartin/genomics_general (*18*). Allowing for some admixture among these recent species, we considered divergent SNPs to be those that were nearly fixed between specialist species (*F_st_* ≥ 0.95; Fig. S2; Table S2–S3; Data S2-S3). All divergent variants that were also in a region of a hard selective sweep (see section below) were also in the 90^th^ percentile of *D_xy_* across all genomic windows, as calculated for the same 50-kb windows using the python script popGenWindows.py.

A window size of 50-kb for sliding window tests was based on the extent of linkage disequilibrium (LD) along a scaffold. We calculated LD decay from pairwise calculations of LD between all SNPs within 100-kb of each other along the largest scaffold in the genome using PLINK’s LD function (--*r^2^*). Linkage disequilibrium decayed to background rates after 50-kb at a threshold of *r^2^* ≥0.1 (Fig. S6).

### Mutation rate estimation

The spontaneous mutation rate for Caribbean pupfishes was estimated from high coverage sequencing (15-69x) of parents and offspring from two independent pedigreed crosses of San Salvador Island species: one cross between a second generation inbred lab-reared generalist and third-generation inbred lab-reared molluscivore individual from Little Lake and another between a second-generation lab-reared generalist and second-generation lab-reared scale-eater from Little Lake. Using the same pipeline for alignment to the reference genome and variant calling with GATK as above, we obtained 9 million variants across 7 individuals from these two crosses after using GATK’s recommend hard filter criteria (i.e. QD < 2.0; FS < 60; MQRankSum < - 12.5; ReadPosRankSum < −8). We independently called variants for these same individuals again using samtools mpileup (v1.9) with the command line arguments *bcftools mpileup -Ou | bcftools call -m -Ob -f GQ,GP*. For both sets of variants (GATK and samtools), poorly mapped regions were then removed using a mask file generated from the program SNPable (http://bit.ly/snpable; k-mer length =50, and ‘stringency’=0.5). We further excluded sequences in which indels were called in any sample, as well as 3 bp of sequence around the indel.

After variant calling, we searched for new mutations in the offspring: sites where an offspring is heterozygous for an allele not found in either of the parents. To determine a set of filters for new mutation sites in the offspring, we first looked for variants which were heterozygous in the offspring and alternatively homozygous in the parents (i.e. known heterozygous sites). Eleven measures of variant quality scores for these known heterozygous sites in the offspring were then used to filter sites for new mutations in the offspring (e.g. depth, genotype quality, mapping quality; see Table S6 for full details) following similar pipelines and filters from several previous studies ((*19–21*)). For example, only new mutation sites that had a depth within 2 standard deviations of the mean depth of the known heterozygous sites in the offspring were kept (all thresholds reported in Table S6). Additionally, new mutations in the offspring were determined from sites in which parents were homozygous for the reference allele and the offspring were heterozygous with quality scores within those of the known heterozygote sites (Table S6) and had an allele balance score between 0.3 and 0.7. This set of variants was then filtered for those independently called in both GATK and samtools runs (Table S6) with no known segregating sites in the larger Caribbean population dataset (i.e. not sites where the alternative new allele in the offspring was found in four or more copies in the larger dataset).

Using the GATK function *callable loci*, we then determined the ‘accessible genome’: the total number of base pairs from the genome in which mutations could be confidently called for each cross. This number was estimated using similar variant quality filters as for the new mutation estimate, excluding those filters that were only applicable to the new mutations and heterozygous sites (i.e. filters assessing quality of alternative allele calls). This meant excluding genomic regions where 1) read map depth in any member was not within two standard deviations of the average read map depth (varies by sample; Table S6), 2) regions with mapping quality scores less than 50, and 3) regions with base quality scores less than 30.

Since the new mutations observed could have originated on either chromosome, the point estimate of per site mutation rate is the number of new mutations observed divided by two times the size of the accessible genome. The mutation rates were then averaged across individual offspring for each cross (Table S6) to obtain a mean mutation rate estimate of 1.56 x10^−8^ mutations per site per generation. This is faster than mutation rate estimates for other teleosts (*20, 22, 23*); however, short-lived smaller species with higher metabolism rates like pupfishes are expected to exhibit faster mutation rates (*24*). We estimate generation times in the field to be approximately one year based on laboratory and field (*25*) longevity studies.

### Demographic Inferences

Various demographic histories can shift the distribution of low- and high-frequency derived variants to falsely resemble signatures of hard selective sweeps. In order to account for demography in downstream analyses, we used the MSMC (v. 1.0.1; *24*) to infer historical effective population size (N_e_) changes in seven focal populations. We ran MSMC on unphased GATK-called genotypes separately for an individual in each of seven focal populations (excluding generalist *C. higuey* due to poor sequencing quality of our single high-coverage individual) with higher sequencing coverage (17-28x mean coverage across individuals; Fig 1D;Table S7). As recommended in the MSMC documentation, we masked out sites with less than half or more than double the mean coverage for that individual or with a genotype quality below 20. We also excluded sites with <10 reads as recommended by Nadachowska-Brzyska et al. (*27*). To scale the output of MSMC to real time and effective population sizes, we used a one-year generation time (*24*) and the estimated spontaneous mutation rate of 1.56 x10^−8^ for Caribbean pupfishes (see previous section).

### Introgression

We characterized differential introgression between specialists in the SSI radiation on both a genome-wide and local level. We visualized the directionality of hybridization and introgression on a genome-wide level using *TreeMix* (v 1.13; (*28*)). *TreeMix* estimates a maximum likelihood phylogeny of the focal populations and then fits a user specified number of migration edges to the tree by comparing genetic covariances of allele frequencies among populations. We ran *TreeMix* with *C. dearborni* as the root node, and with 0 through 20 migration edges. The most likely number of migration events was chosen using the broken-stick approach (Fig. S7).

We investigated how signatures of hybridization at the genome-wide level contributed variation potentially important to the divergence between species using the *f_d_* statistic, which is designed to look for signatures of introgression across sliding genomic windows (*18*). The *f_d_* statistic, a modified version of the *D*-statistic, looks at allele frequencies fitting two allelic patterns referred to as ABBA and BABA based on the tree (((P1,P2),P3),O), where O is an outgroup species in which no gene flow is thought to occur with the other populations (*18*). We used 2 individuals of *C. artifrons* from Cancun, Mexico as our distantly related outgroup population for this test, which forms the deepest divergence event with *C. variegatus* within the *Cyprinodon* clade (*1*), and focused on introgression between San Salvador Island specialists and Caribbean populations. Based on the tree (((P1,P2),P3), *C. artifrons*), the *f_d_* statistic was calculated for the combinations of populations in which P2 was either the scale-eater or molluscivore and P3 was one of the Caribbean outgroup populations (Table S8 and S9) in 50-kb sliding windows with a minimum of 100 variant sites with no missing data within a population using the ABBABABA.py script (available on https://github.com/simonhmartin/genomics_general;(18)). To compare these patterns of introgression into the specialist to patterns of introgression into focal generalist populations on other islands, we also calculated *f_d_* statistics across the genomes of generalist populations on islands where we had population-level sampling and sister groups to fit the relationship expected for the test (Table S9B and S9D).

Significance of *f_d_* values in sliding windows across the genome was evaluated using simulations with no migration using ms-move (*29*) using estimates of effective population size changes from our MSMC analyses and a divergence time between the two specialists set to 10,000 years, the age of the hypersaline lakes on SSI (Table S8). Significant regions were determined by calculating the *f_d_* statistic across a genome simulated under a coalescent model with no gene flow and consisting of 150,000 windows, each containing the mean number of variants found across 50-kb windows in the empirical genomes. Empirical windows were considered candidate introgressed regions if the *f_d_* statistic was above the maximum simulated *f_d_* value (Table S8). Consecutive 50-kb *f_d_* outlier windows were merged to estimate the range in the sizes of introgressed regions to approximate the age of introgression events (Fig. 2D).

### Candidate adaptive variants underlying San Salvador Island specialist phenotypes

In the specialists, we also looked for regions that appeared to be under strong divergent selection in the form of a hard selective sweep from the site frequency spectrum calculated with SweeD (v.3.3.4;(*30*)). In this calculation of the composite likelihood ratio (CLR) of a sweep, we incorporated our empirical estimate of the decrease in population size for each focal population estimated from MSMC analyses in 50-kb windows across scaffolds that were at least 100-kb in length (99 scaffolds; 85.6% of the genome). We also calculated CLRs across 100,000 scaffolds consisting of neutrally evolving sequences simulated with ms-move (*29*), controlling for the impact of the inferred population size decreases over time for each population from MSMC runs mentioned above (Fig. 1D; Table S7). The CLR ratios for the simulated datasets were then used to assess outlier CLR ratios from the empirical dataset. Regions with CLR ratios above the 95^th^ percentile value of CLR from the neutral simulated dataset were considered candidate hard selective sweep regions (scale-eater: CLR > 5.28; molluscivore: CLR > 4.47; Table S7). Candidate hard selective sweep regions were also independently inferred for the five focal Caribbean generalist populations (sample size ≥ 10) following the same method outlined above for the specialists (Table S7).

### Spatial distribution of candidate adaptive variants

To determine candidate adaptive variants underlying the divergence observed on San Salvador Island, we looked for regions of the genome that contained signatures of strong genetic divergence and hard selective sweeps. Variants that were nearly fixed between the species on San Salvador (*F_st_* ≥0.95) and located in a candidate selective sweep region (empirical CLR > demographic simulations CLR; Table S7) were considered candidate adaptive variants (Table S2–S3; Data S2-S3). We then surveyed all pupfish individuals sampled from outside these populations for the specialist allele at their respective candidate variant sites (e.g. whether or not the scale-eater allele was found in other populations across the Caribbean). Variants were then separated into two categories of genetic variation: *de novo* (the specialist allele was found only on San Salvador Island) or standing (the specialist allele was also found in at least one generalist population sampled outside of San Salvador Island). Standing genetic variation that was found in candidate introgression regions from the *f_d_* tests for introgression (see Introgression section) was further parsed into the category of introgressed variation and separated by geographic region of introgression (North Carolina (NC), New Providence Island (NP), or Dominican Republic (DR)). Given the amount of candidate adaptive variation that exists as standing genetic variation across the Caribbean (Fig. 2A), we looked for how many of these candidate regions also showed evidence of hard selective sweeps in focal generalist populations outside of San Salvador Island and found that very few of these regions exhibited signatures of a hard selective sweeps outside of San Salvador Island populations (Fig. S3).

### Adaptive introgression

We examined all introgressed regions for evidence of hard selective sweeps on SSI and genetic divergence between trophic specialists, which would provide evidence that secondary gene flow brought in variation potentially important for speciation. We looked for overlap between the *f_d_* introgression outliers, SweeD selective sweep outliers, and whether these overlapping regions contained at least one of the nearly fixed variants (*F_st_* ≥ 0.95) between specialists (*31*) (Table S10–S14).

We were interested in whether San Salvador Island specialist genomes exhibited more introgression in regions undergoing hard selective sweeps on SSI than other generalist populations. In the absence of a clear null expectation for the number of introgressed regions, we calculated the number of these candidate adaptive regions for the specialists that were also outlier *f_d_* regions in other combinations of populations across the Caribbean (Table S9), to compare to the number of adaptive introgression regions observed in the specialists. Since several outgroup generalist populations had multiple values for the number of adaptive introgression regions (due to different combinations of sister lineages (P1) available for testing against: Table S9), only the average number of adaptive introgression regions per generalist population was shown for ease of visualization (Table S9; Fig. 3C). We then performed a Mann-Whitney U test to determine if the mean number of adaptive introgression regions in each specialist was greater than the mean from the rest of the Caribbean (Table S9A v. S9B and Table S9C v. S9D) and calculated 95% confidence intervals around these means using the boot.ci function in the R package boot (v1.3; Fig. 3C). Since neither of the SSI specialists appear to have experienced adaptive introgression from the Venezuela *C. dearborni* population, it was excluded as a potential donor population for the focal generalist populations on other islands as well in these comparative analyses.

### Functional analysis of candidate adaptive variants

#### GO analysis of candidate variant regions

We performed gene ontology (GO) enrichment analyses for genes near candidate adaptive variants using ShinyGo (v.0.51;(*32*)). In the *C. brontotheroides* reference genome annotations (described in *de novo* genome assembly and annotation section), gene symbols largely match human gene symbols. Thus, we searched for enrichment across biological process ontologies curated for human gene functions.

For genes with focal GO terms (e.g. feeding behavior, muscle, mouth, eye and craniofacial development) relevant to stages of diversification in this system (i.e. habitat preference, trophic morphology, and pigmentation; Fig. 2C; Fig. 4; Table S4), we checked other annotation databases and studies for verification of putative function, including Phenoscape Knowledgebase (https://kb.phenoscape.org/#/home), NCBI’s PubMed (https://www.ncbi.nlm.nih.gov/pubmed), and the Gene Ontology database using AMIGO2 (*33*). All genes had consistent annotations across databases, except *galr2*. *Galr2* was annotated for feeding behavior in the Biological Processes database (Ensemble 92), but to our knowledge, the most recent studies indicate that it does not play a role in feeding behavior (*34, 35*). Thus, we removed its annotation as a candidate gene for feeding behavior, but kept it as a candidate for trophic morphology.

#### Differential gene expression

Additionally, we looked for overlap between genes associated with candidate variants and genes differentially expressed between the two specialists in whole embryos at two early developmental stages (2 and 8 days post-fertilization (dpf)) reported in a previous study (*36*). Tables with differentially expressed genes at 2 and 8 dpf from this study are provided in Data S4 and S5.

#### Morphometrics and caudal fin pigmentation

We measured two key morphological traits associated with the major axes of phenotypic diversification in the SSI radiation, lower jaw length and nasal protrusion distance. Ethanol-preserved specimens from SSI were measured from external landmarks on the skull using digital calipers. Measurements were repeated on both lateral sides and averaged for each specimen. Lower jaw length was measured from the quadrate-articular jaw joint to the tip of the most anterior tooth on the dentary (Data S6). Nasal protrusion distance was measured by placing a tangent line from the dorsal surface of the neurocranium to the tip of the premaxilla and measuring the perpendicular distance that the nasal region protrudes from this tangent (Fig. S8A; Data S6). Each specimen was also measured for standard length using digital calipers in order to remove the effects of variation in body size on the craniofacial trait measurements among individuals and species. We log-transformed morphological measurements and regressed them against log-transformed standard length (Fig. S9; Data S6) and used the residuals for association mapping analyses.

The major axis of divergence in reproductive coloration and patterning between trophic specialists on SSI is the overall lightness or darkness of breeding males. Scale-eaters reach a nearly jet black coloration in the wild while guarding a breeding territory whereas molluscivore males remain paler throughout their body and fins. This pair of sympatric specialists exceeds the lightness contrast in male reproductive breeding coloration observed across all other *Cyprinodon* pupfishes. Females of each species show the same general pattern of lightness/darkness. We detected no difference in the total number of melanocytes on the caudal, anal, or pectoral fins among the SSI species (data not shown). Instead, we found that scale-eater individuals were significantly darker on their caudal fins (two-tailed *t*-test, t=5.25, df=45.5, *P*-value= 3.8×10^−6^; Fig. 4B; Data S6), perhaps due to the larger melanocyte areas relative to molluscivores. We found similar patterns for anal and pectoral fins and used only caudal fin lightness values for GWAS (data not shown). A Meiji EMZ-8TR stereomicroscope with standardized external illumination and an OMAX 18 Mp digital microscope camera was used to take lateral photographs of the caudal fin of each individual against the same white reference background in each image (Fig. 4B;Data S6). Adobe Photoshop (Creative Cloud) was used to select a rectangular area from inside the caudal fin, not including the caudal peduncle region or terminal marginal band and measure the mean overall lightness of this region relative to a control region selected from the illuminated white background. Standardized caudal fin pigmentation was then calculated as the proportion of the caudal fin lightness value relative to the control background lightness value for downstream GWAS analyses.

#### Genome-wide association mapping analyses

We employed a Bayesian Sparse Linear Mixed Model (BSLMM) implemented in the GEMMA software package (v. 0.94.1; (*37*)) to identify genomic regions associated with variation in lower oral jaw length, caudal fin pigmentation, and nasal protrusion distance. The BSLMM uses Markov Chain Monte Carlo sampling to estimate the proportion of phenotypic variation explained by all SNPs included in the analysis (proportion of phenotypic variance explained (PVE);Fig. S10A-C), explained by SNPs of large effect (proportion of genetic variance explained by sparse effects (PGE); Fig. S10D-F), and the number of large-effect SNPs needed to explain PGE (nSNPs;Fig. S10G-I). GEMMA also estimates a posterior inclusion probability (PIP) for each SNP. We used PIP (the proportion of iterations in which a SNP is estimated to have a non-zero effect on phenotypic variation) to assess the significance of regions associated with jaw size variation. We performed 10 independent runs of the BSLMM using a step size of 100 million with a burn-in of 50 million steps for three traits (lower oral jaw size (*n* = 78), caudal fin pigmentation (*n* = 61), and nasal protrusion distance (*n* = 65)). We chose to only include SSI individuals in these analyses given extensive Caribbean-wide population structure that could confound significant associations (Fig. 1C). We summed PIP parameter estimates across 20-kb windows to avoid dispersion of the posterior probability density across SNPs in linkage disequilibrium due to physical linkage. Independent runs were consistent in reporting the strongest associations for the same 20-kb windows. We identified regions with strong association with our traits of interests as ones that had a PIP score in the 99^th^ percentile of PIP scores across all regions (Data S7-9). Our PIP estimates for strongly associated windows suggest that jaw length may be controlled predominantly by a few loci of moderate effect (see bimodal PGE distribution, Fig. S10H). This is consistent with a previous QTL mapping study in an F2 intercross between SSI trophic specialists which detected one significant QTL with moderate effects on oral jaw size explaining up to 15% of the variation and three to four additional potential quantitative trait loci (QTL) with similar moderate effects (*38*).

#### QTL analysis for jaw size

We also investigated candidate variants for effects on craniofacial morphology by overlapping scaffolds with a previously published linkage map and QTL analysis of an F2 intercross between specialist species (*38*). We overlapped markers from this study that spanned the 95% Bayesian credible interval for a significant QTL for lower jaw length (LG15; taken from Fig S2 in (*38*)). The fasta sequences for these two markers bookending the QTL region on a single scaffold were then blasted against the *Cyprinodon brontotheriodes* genome using the blastn function in BLAST+ (*39*) and we selected the result with the highest percent identity and lowest e-value (Table S5). We then looked at all the genic regions within the interval between these two markers to investigate overlap between the QTL region and the candidate variants in this current study. The top hits for overlap between the sequences of two markers that spanned the LG15 QTL region and the *Cyprinodon brontotheroides* reference genome showed that this QTL corresponds to a 18 Mb region on scaffold c_bro_v1_0_ scaf8 (Table S5). However, this large region contained only a few candidate adaptive variants associated with the genes *map2k6* (3 variants), *galr2* (2 variants), and *grid2ip* (4 variants).

### Timing of divergence for candidate adaptive variants

If adaptive diversification in this radiation of pupfishes followed the temporal stages hypothesis (habitat preference first, then trophic morphology, and finally sexual communication; (*40*)), we predicted an ordering of divergence times among candidate regions containing genes annotated for traits related to these axes of diversification, with regions containing genes related to habitat preference having older divergence times, followed by regions with genes related to trophic morphology, and finally regions with pigmentation genes having the youngest divergence times. In order to determine if there have been stages of adaptive diversification in this adaptive radiation of pupfishes, we first estimated divergence times between molluscivores and scale-eaters for each candidate adaptive region (i.e. each 50 kb window containing signatures of a hard selective sweep in either specialist and a nearly fixed variant between specialists).

Many methods for estimating divergence times and allele ages rely on the pattern of variation in the haplotype background surrounding the allele of interest. Heuristic approaches, particularly those that use point estimates of number of derived mutations within a chosen distance of the site are accessible, quick ways to approximate divergence times among regions and allele ages without extensive haplotype data (*41, 42*). We estimated sequence divergence in regions surrounding candidate variants using *D_xy_*, an absolute measure of genetic divergence. To get a heuristic estimate of divergence time between specialists at these regions, we used this *D_xy_* count of the number of variants that have accumulated between specialists and the approximation that the observed genetic differences between two lineages should be equal to 2µt: t, the time since their divergence and µ, the mutation rate (*43*). Using the per generation mutation rate estimated above (1.56×10^−8^), we calculated the time since divergence for candidate regions and compared that time to the estimated 10-15 kya age of the radiation (based on estimates of the last period of drying of hypersaline lake basins on San Salvador Island during the last glacial maximum (*44*)).

We plotted the divergence time estimates for candidate adaptive regions by the spatial categories they were assigned to (de novo, introgression, standing genetic variation), and against a background of neutral regions that contained a nearly fixed variant, but no signature of a hard selective sweep (Fig 4, S5 and S11). So that each point in the plot was independent, we pruned variants by randomly selecting one from the group of variants that fell within the same 50-kb window. Some windows had multiple variants with different spatial distributions (e.g. de novo vs. standing genetic variation), so to make sure all points on the figure were independent, we made alternative plots for alternative spatial distributions of variants that occurred within a single 50 kb window (the smaller vs larger spatial distribution; Fig. 4 and Fig. S11). This applied to several variants that were characterized as either introgressed or standing genetic variation in four regions containing genes with relevant adaptive annotations (*galr2, gmp6a, and kcnk2*) from the plot with the smaller spatial distribution (Fig. 4). The alternative, larger spatial distribution (i.e. the regions containing these genes were assigned to the standing genetic variation column instead of the int rogression column) for each of these four annotated 50 kb windows is shown in Figure S11.

To look for stages of diversification along different trait axes, we matched regions to potential phenotypes in two ways: 1) from our GO enrichment analyses that implicated genes in the region as relevant to the major axes of adaptive radiation in this system (e.g. craniofacial development, pigmentation, feeding behavior), and 2) with regions strongly associated with either lower jaw size, nasal protrusion distance, or caudal fin pigmentation in the GWAS for San Salvador Island pupfish species. Regions associated with traits in the GWAS were then polarized for effects on trait values that were in directions relevant to each specialist. For scale-eaters, this meant filtering for regions with significant association with darker caudal fin pigmentation, larger lower oral jaws, and smaller nasal protrusion distances. For molluscivores, this meant filtering for regions with significant associations for lighter caudal fin pigmentation, larger lower oral jaws, and larger nasal protrusion distances. We found 23 regions containing genes with relevant GO terms and 24 regions containing variants significantly associated with traits in the GWAS.

Based on relevant GO terms, we found that regions with genes annotated for feeding behavior have some of the oldest divergence times (Fig 4A and S11) while regions with genes related to craniofacial morphology and pigmentation had younger divergence times (Fig 4A and S11). Similarly, we found younger divergence times among regions with genes annotated for traits related to trophic morphology based on GWAS annotations (Fig 4B).

Considering there are only 23 and 24 observations used to test for stages of adaptive diversification, we also checked that this pattern was statistically robust and not due to chance. First, we used linear models to determine if divergence times among adaptive variant regions could be predicted by the associated adaptive traits. In our first linear model used to test for stages, major GO categories relevant to divergent traits in this system (feeding behavior, eye, mouth, muscle, and craniofacial development) were used as the predictor variable and the rank ordering of divergence times was the response variable. We found that relevant GO categories significantly predicted the rank ordering of divergence times among regions (*df*=5, *F*= 3.03, *P*-value= 0.039). From this, we then tested all pairwise comparisons of different categories of relevant traits to determine which were driving the significant difference in ordering using Tukey’s HSD test. For the stages of trait diversification based on relevant GO terms (feeding behavior, eye, mouth, muscle, and craniofacial development): diversification in feeding behavior was significantly older than mouth development (*P* = 0.022), and moderately older than muscle and craniofacial development (*P* = 0.05 for both). This conservative test supports a ‘behavior-first’ ordering to trait diversification in this system (*45*). Divergence in behavioral traits as a major stage of radiation in pupfish makes sense because both specialists are more aggressive than generalist species (*46*), and adaptive kinematic traits important for scale-eating performance are behaviorally mediated (*47*).

Evidence for the behavior first stage of adaptive diversification is largely driven by the fact that the two oldest divergences times occur in regions with genes annotated for feeding behavior. Therefore, we further investigated the probability that this ‘behavior first’ pattern could have occur by chance given our small sample size. We used a permutation test to calculate the probability of the two regions with the oldest divergence times both being related to feeding behavior can occur randomly. We first ranked the 23 candidate regions based on their corresponding age and randomly reassigned these rankings for each region 10,000 times. We then assessed how many of each of these iterations had the two regions with the oldest divergence times assigned to feeding behavior (as observed in the empirical data). We found that only 41 of the iterations matched the observed value, suggesting that the probability of observing the two regions with the oldest divergence times related to feeding behavior by chance alone is extremely small (*P-value* = 0.0041).

In our second linear model to test for stages of adaptive radiation, significant association with three key adaptive traits measured for our GWAS (lower oral jaw size, caudal fin pigmentation, and nasal protrusion distance) was used as the predictor variable and ordering of divergence times was the response variable. We found that GWAS trait association did not significantly predict the ordering of divergence times among regions (*df*=3, *F*= 1.67, *P*-value = 0.197). However, this is qualitatively similar to our results from the GO analyses considering that we lack a behavioral trait related to feeding behavior in our GWAS analyses.

### Timing of selection for candidate adaptive variants

We also estimated the age of hard selective sweeps using the R package *McSwan* (v1.1.1;; https://github.com/sunyatin/McSwan; (*48*)). *McSwan* detects hard selective sweeps by comparing local site frequency spectra (SFS) simulated under neutral and selective demographic models, which it uses to assign selective sweeps to regions of the genome and predict the age of selection events (*48*). By using information from the SFS, *McSwan* is advantageous for estimating selective sweep ages in non-model organisms because it does not require high quality haplotype data to detect sweeps and predict their ages. However, this flexibility comes at the cost of not jointly estimating the selection coefficient of a particular sweep, so it assumes the strength of selection is equal across all sweeps. With the input of a mutation rate estimate, neutral demographic model (effective population size changes, divergence events), and variant file, *McSwan* generates simulated and observed SFSs and a prior of sweep ages, whose upper bound is determine by the divergence time estimate specified in the demographic model (in our case: 10,000 years). *McSwan* uses these simulated selective and neutral SFSs to scan the input variant file for selective sweep regions and produce a posterior distribution of sweep ages for each sweep region it detects.

To simulate SFSs, we used the mutation rate estimated above (1.56×10^−8^), the same demographic models of changes in effective population sizes used in our SweeD runs for the generalists and scale-eater populations (Table S7), and a divergence time estimate between San Salvador Island generalist and scale-eater of 10,000 years (based on dating of last glacial maximum when the hypersaline lakes were dry on San Salvador Island). We first simulated neutral and selection SFSs that were each comprised of 2,000 simulations (default recommendation) across sequences 50-kb in length. To look for selective sweeps in the specialists, we then generated empirical SFSs from scans across the 500-kb region surrounding each of the 23 sets of candidate variants highlighted in Figure 4. In order to precisely determine the boundaries of hard selective sweep, McSwan iterates its genomic scans over adjacent windows of various lengths and offsets and compares the empirical SFS to the simulated SFS under selection to assign regions as selective sweeps. We set up the iterative scans across these 500-kb regions in sliding windows that ranged from 1000 bp to 200-kb in length and a minimum of 50 variants required per window. Each sliding scan of the 500-kb region was done in 100 overlapping steps (default setting). We then looked for overlap between the regions detected as hard selective sweeps by McSwan to the candidate regions previously detected with SweeD and *F_st_* (Table S2–S3). For the scale-eater population only 9 of the 23 sets of candidate variants detected as hard selective sweeps using SweeD were also detected as hard selective sweeps using *McSwan* and given age estimates. In Tournebize et al. 2019 (*48*), investigations of McSwan’s performance noted poor performance to detect selective sweeps when selection was relatively weak (s ≤0.05) and recent (≤16 kya; Supplemental information Section 2 of (*48*)). Thus, the thirteen additional candidate adaptive regions undetected by McSwan may be under weaker selection or more recent. The candidate variants surrounding the consecutively spaced genes *cenpf* and *kcnk2* were detected as within the same selective sweep in *McSwan* and thus have the same age estimates (Fig. 4C and Fig. S12).

For these 8 regions, we then filtered these distributions of sweep ages for estimates that had a stability value (a parameter that represents the strength of support for a selective sweep model over a neutral model) in the 95^th^ percentile. To get a likely range of selective sweep age estimates for each region, we calculated the 95% high posterior density (HPD) region with the R package HDIntervals (v0.2; https://cran.r-project.org/web/packages/HDInterval/index.html) from their respective posterior distributions. We repeated this process for the 6 sets of candidate regions found in the molluscivore, only three of which were also detected as being under a selective sweep in *McSwan* and given age estimates. The 95% HPD of these age estimates for the scale-eater and molluscivore populations are presented in Figure 4C, S5 and Table S14 and the full posteriors are shown in Figure S12 and S13.

---

**Fig. S1.**
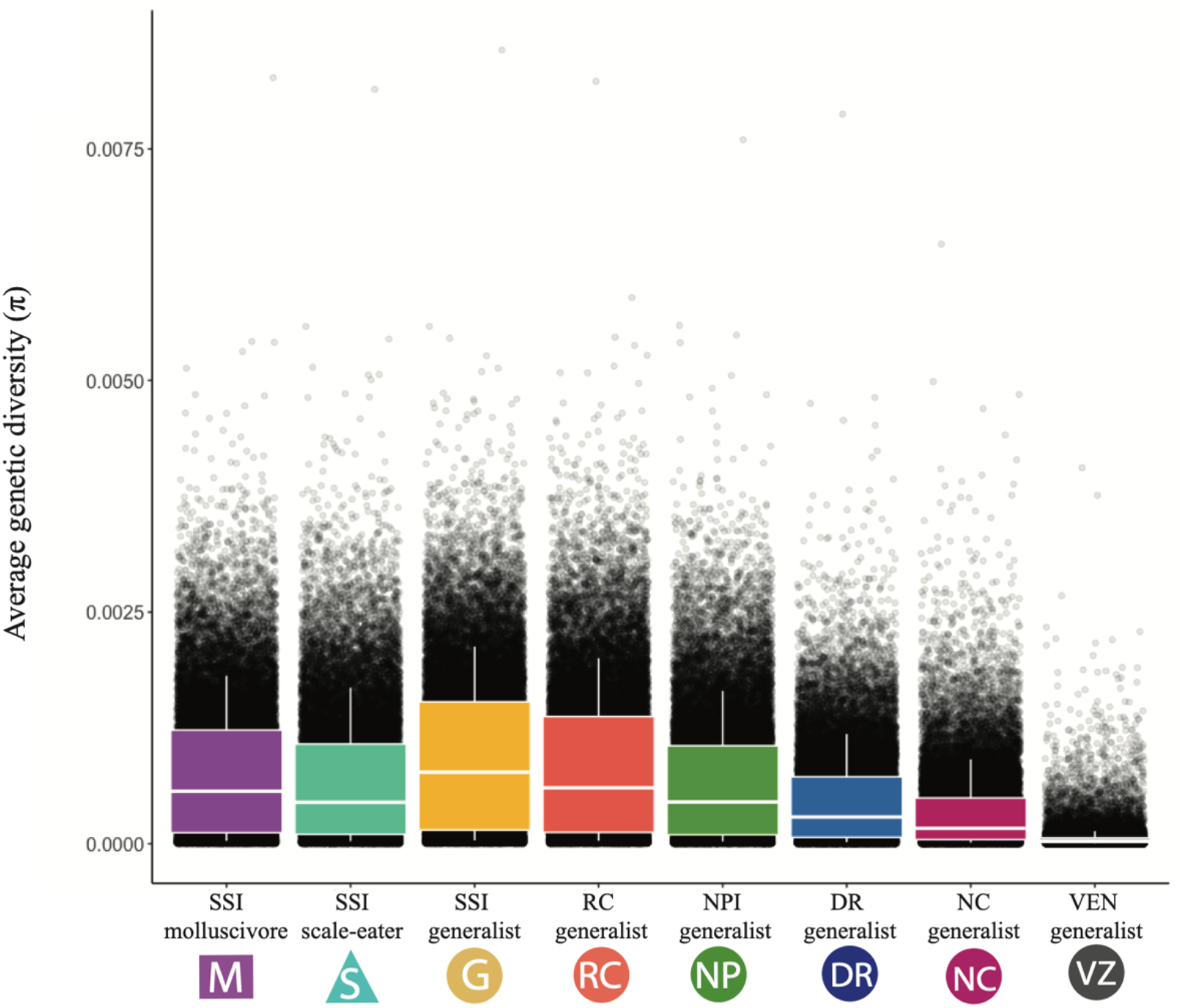
Average genetic diversity is similar across Caribbean pupfish populations. Within population (π) nucleotide diversity in 50-kb sliding windows across the genomes of the San Salvador Island (SSI) species and generalist species on Rum Cay (RC), New Providence Island (NPI), Dominican Republic (DR), North Carolina (NC) and Venezuela (VZ). π values are averaged across 100 random samples of 10 individuals from each population in order to down-sample from populations with larger sample sizes and compare π across populations.

**Fig. S2.**
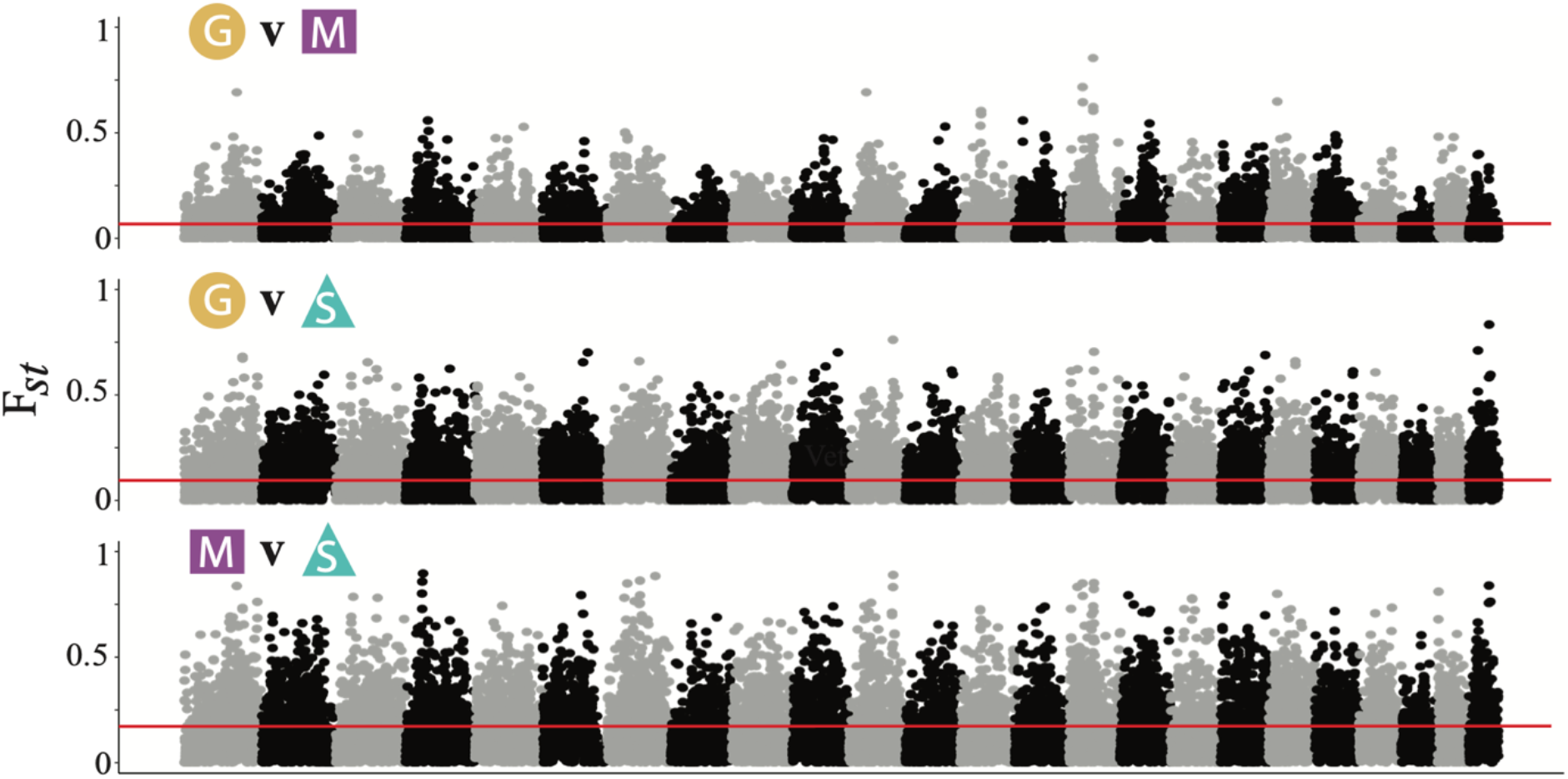
Genetic divergence among SSI species. Manhattan plot of *F_st_* in 50-kb windows across the genome for the three SSI species on the largest 24 scaffolds in the molluscivore (*C. brontotheroides*) genome corresponding to the 24 chromosomes in *Cyprinodon* (*49*). Solid red line represents the average *F_st_* values for each comparison (generalist vs. molluscivore; 0.07; generalist vs. scale-eater: 0.11; molluscivore vs. scale-eater: 0.15).

**Fig. S3.**
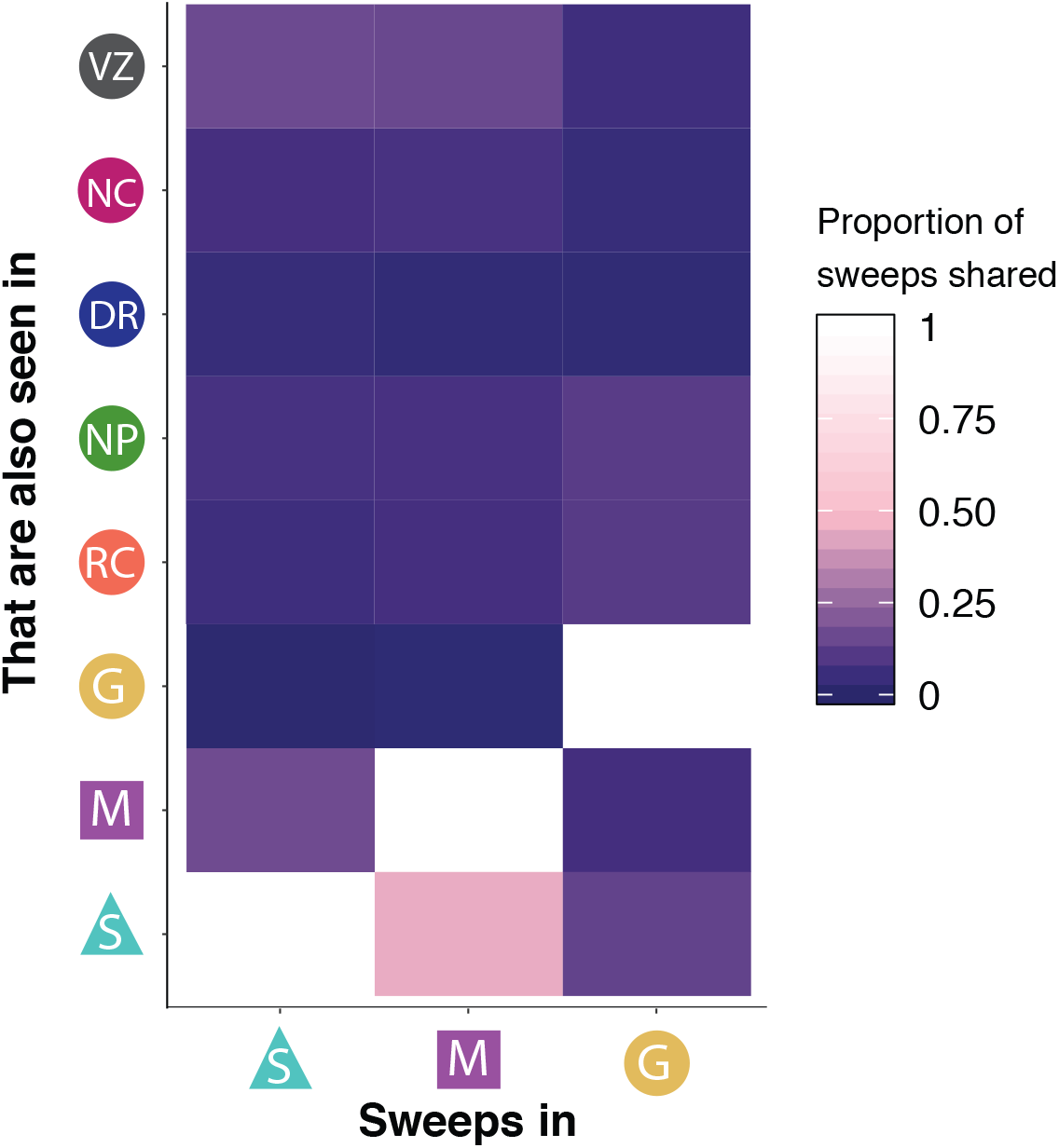
Selective sweeps in SSI population shared with other Caribbean populations. The proportion of hard selective sweeps in the SSI species that are also found sweeping in other Caribbean populations. Note that 42% of selective sweeps in the molluscivore population also showed signs of a sweep in the scale-eater population.

**Fig. S4.**
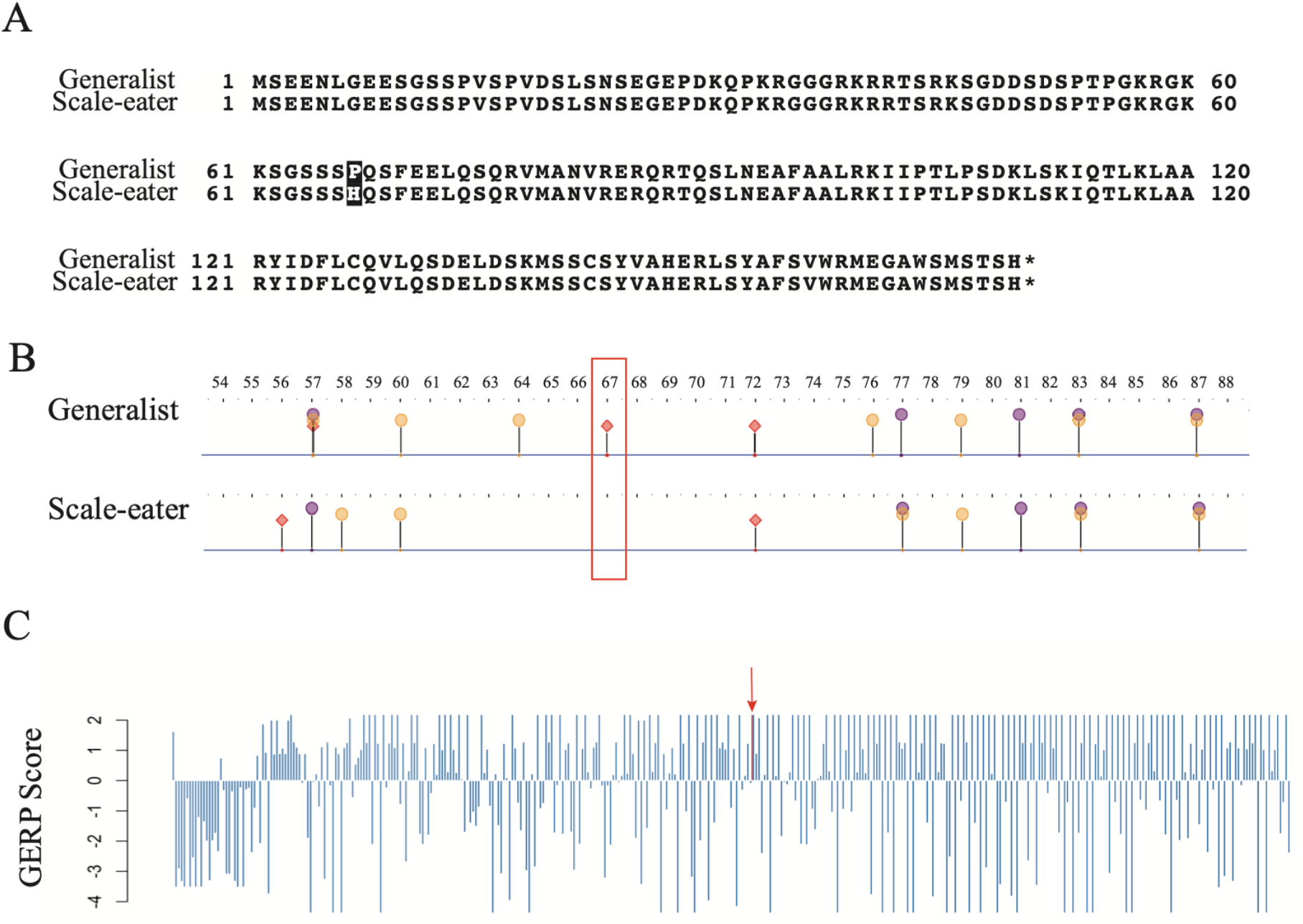
Sequence conservation among fishes around candidate gene *twist1*. A) Amino acid sequence of *twist1* protein for SSI generalists and scale-eaters. The non-synonymous substitution that is nearly fixed between the two species changes the amino acid from a proline to histidine (highlighted in black). B) This amino acid substitution alters a protein binding site (highlighted in red box) predicted and visualized with Predict Protein Open (https://open.predictprotein.org) using the machine-learning prediction method PPsites2 (*50*). C) GERP scores for the 500 base pair region surrounding the non-synonymous coding substitution in *twist1* (red arrow) found only on SSI. Conservation scores were obtained from aligning scale-eater genomes to the 60 fish EPO low coverage genome alignment on Ensembl (release 98) and a conservation score above 2 is considered highly conserved (*51*).

**Fig. S5.**
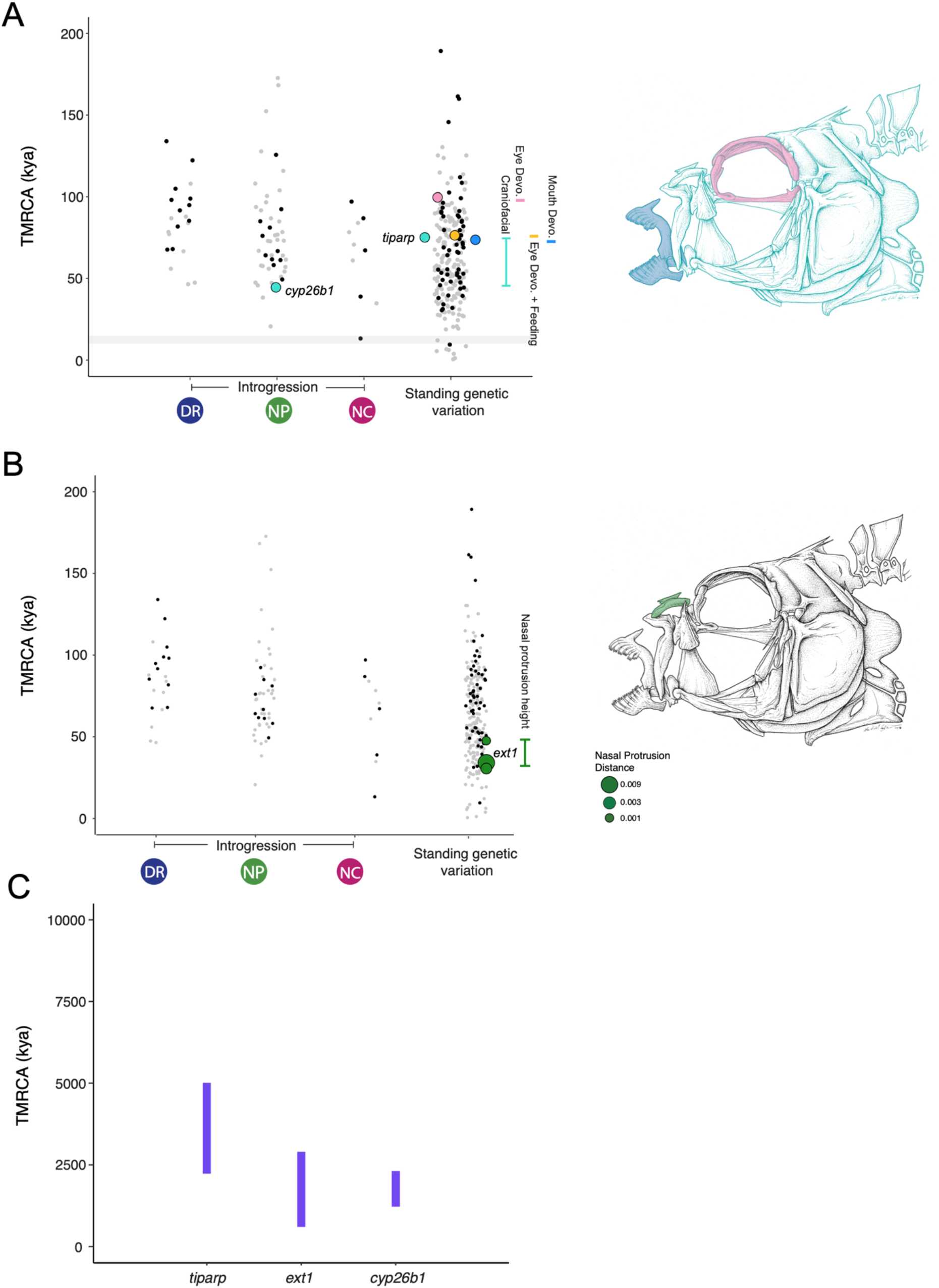
The spatiotemporal landscape of genetic variation for the molluscivore. Time to most recent common ancestor (TMRCA) of the region surrounding candidate adaptive variants (LD-pruned so that each point is independent) based on relative genetic divergence metric *Da* (*52*) which captures only the amount divergence that has accumulated since the two populations diverged. Variants are separated by spatial distribution into: introgressed from a single population and standing genetic variation observed in more than two populations. No variants existed as de novo variation in the molluscivore populations on SSI. No nearly fixed variants introgressed from the Venezuela population. Points are colored by their adaptive relevance to this system (black points: adaptive variants annotated for non-focal GO terms or unannotated; gray points: nearly fixed variants between specialists that are not under hard selective sweeps). A) All variants annotated for GO categories from the Biological Processes database (Ensemble 92) for feeding behavior, muscle, eye, and mouth development, or craniofacial (mouth and/or eye, muscle) are shown. B) All variants in the 99^th^ percentile of PIP scores for association with either smaller oral jaw size, lighter caudal fin pigmentation, or premaxillary nasal protrusion from GWAS are shown. Dot sizes represent PIP score of each GWAS hit within a trait. C) The 95% high posterior density of the posterior distribution of ages of selective sweeps in focal gene regions. Focal regions are colored based on GO and GWAS annotations relevant to the stages of adaptive radiation hypothesis: habitat preference (feeding behavior: red), trophic morphology (craniofacial and muscle: blue-violet), and sexual communication (pigmentation: orange).

**Fig. S6.**
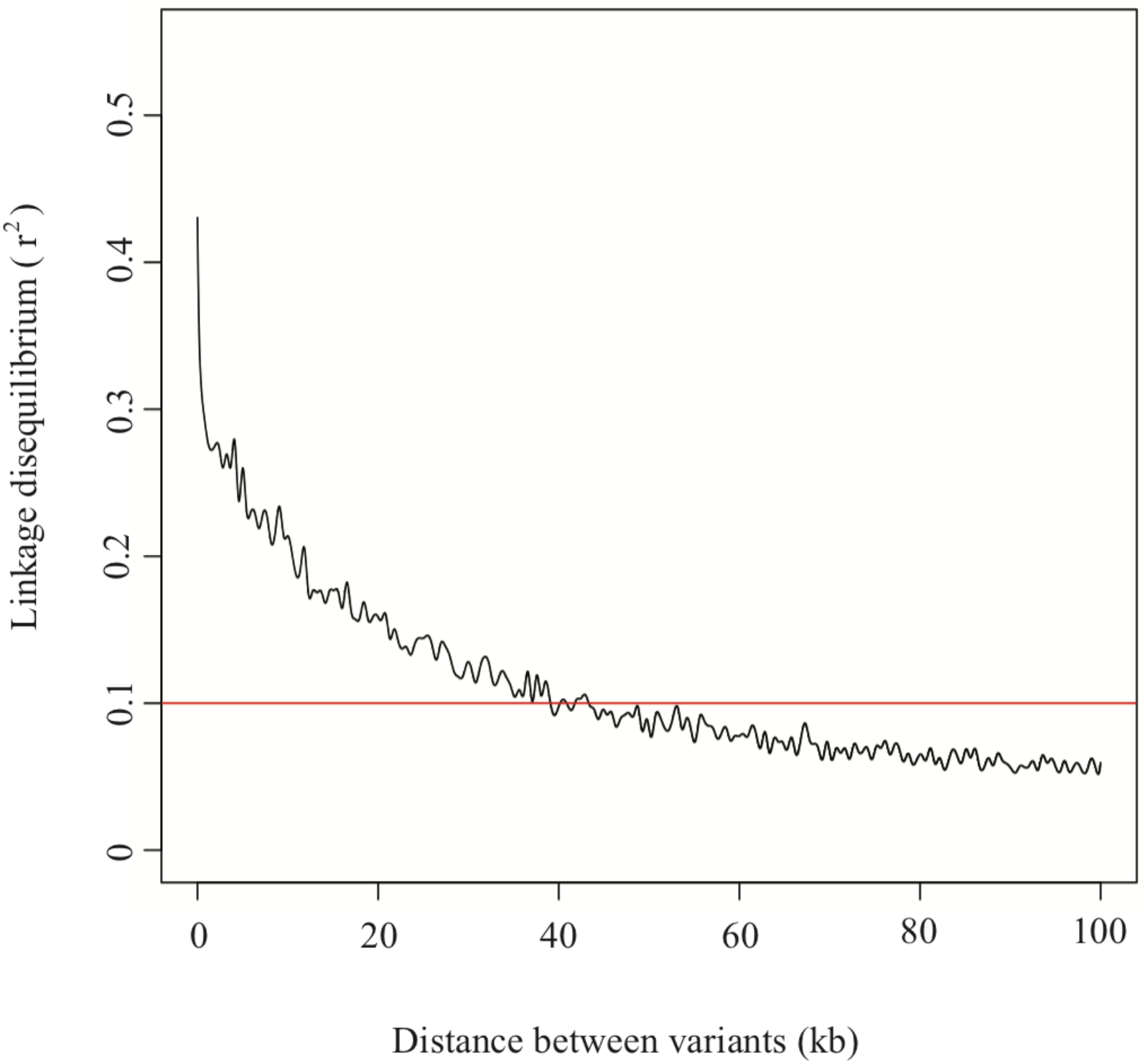
Linkage disequilibrium decay along the genome. LD decay over pairwise combinations of variants within 100 kb of each other on the longest scaffold in the genome (49,059,223 bp), with r^2^=0.1 marked for reference. From this pattern of decay, we chose a window size of 50-kb for sliding windows analyses used in this study.

**Fig. S7.**
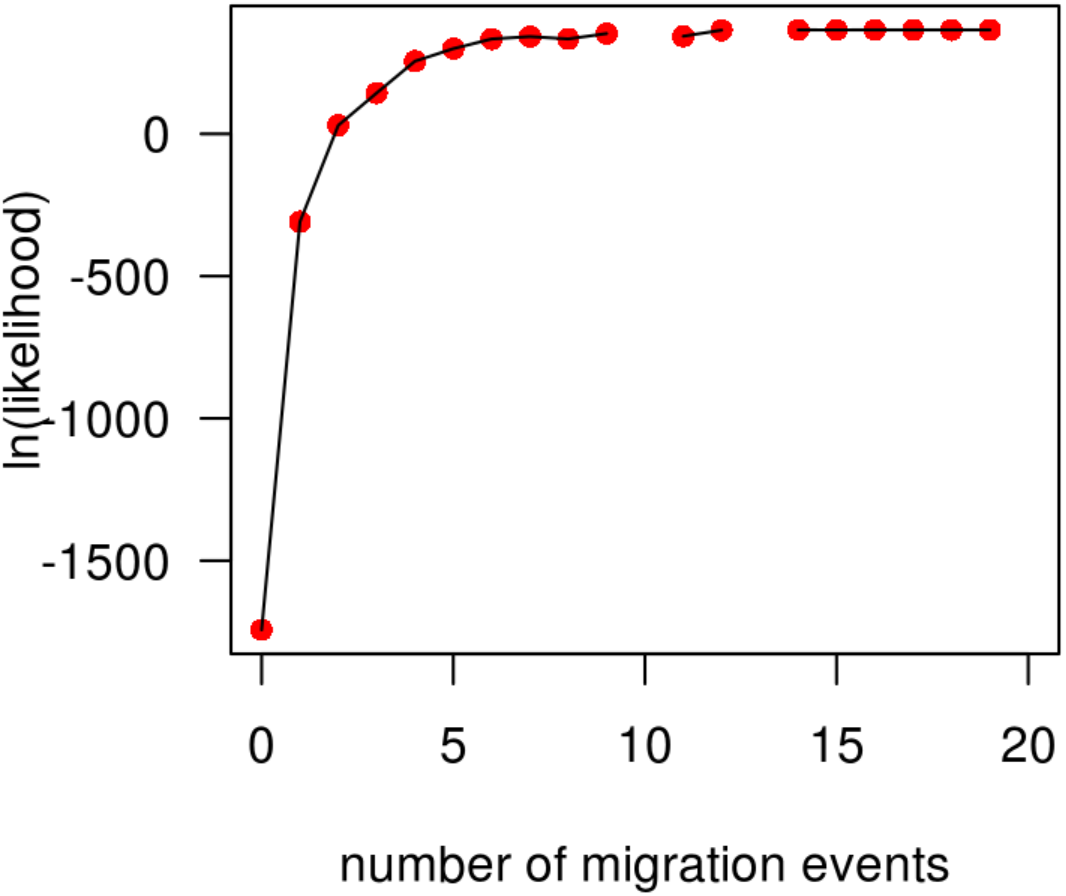
The log likelihood of number of migration events on the population graph inferred using TREEMIX (*28*). The rate of change in the likelihood began to decline after three migration events, so three migration arrows were included in the population graph in Fig. 4B.

**Fig S8.**
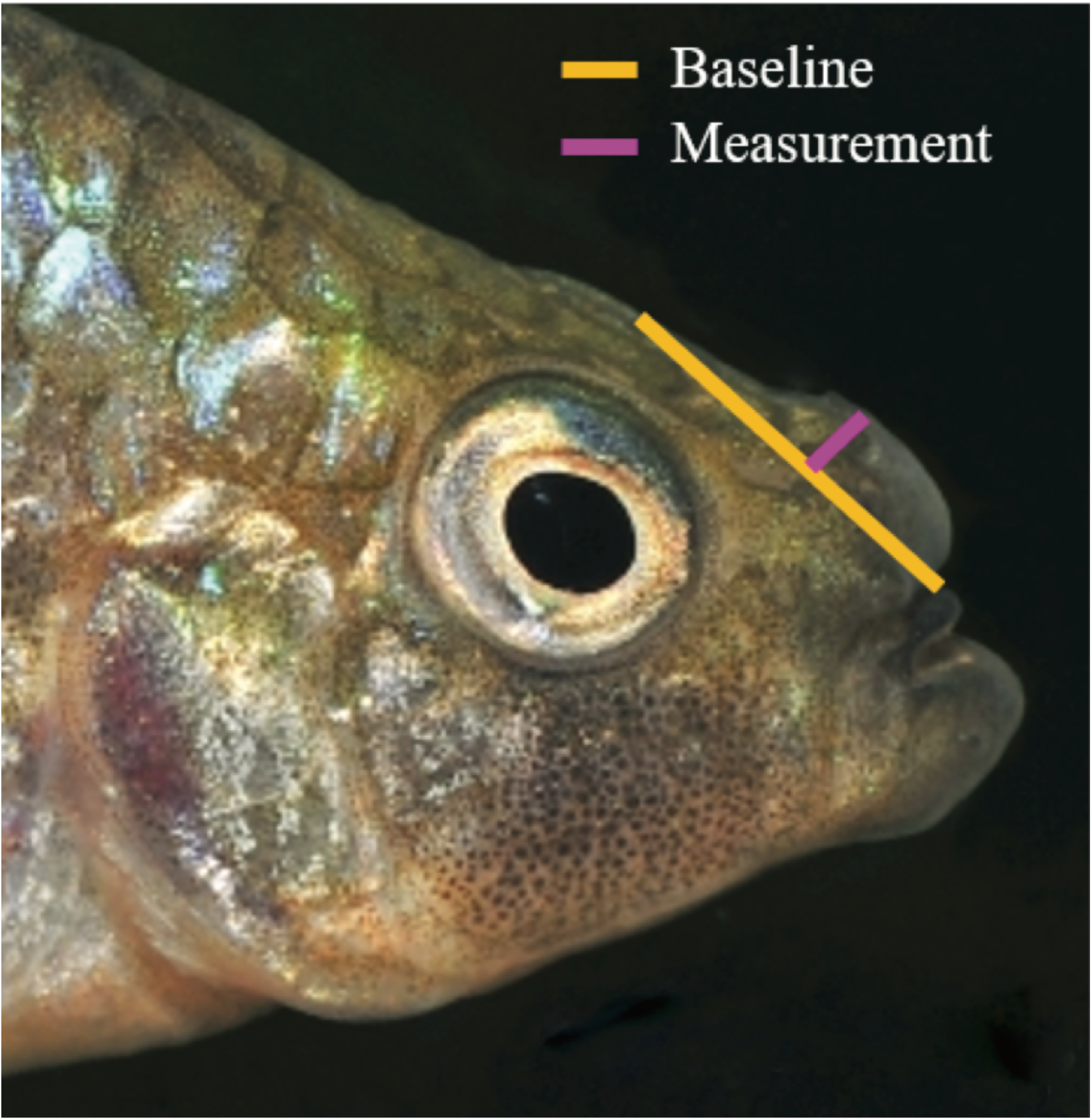
Example image of nasal protrusion distance measurement for GWAS. The Purple line represents the nasal protrusion distance. The yellow line represents a baseline tangent line from the dorsal surface of the neurocranium to the tip of the premaxilla used for reference.

**Fig. S9.**
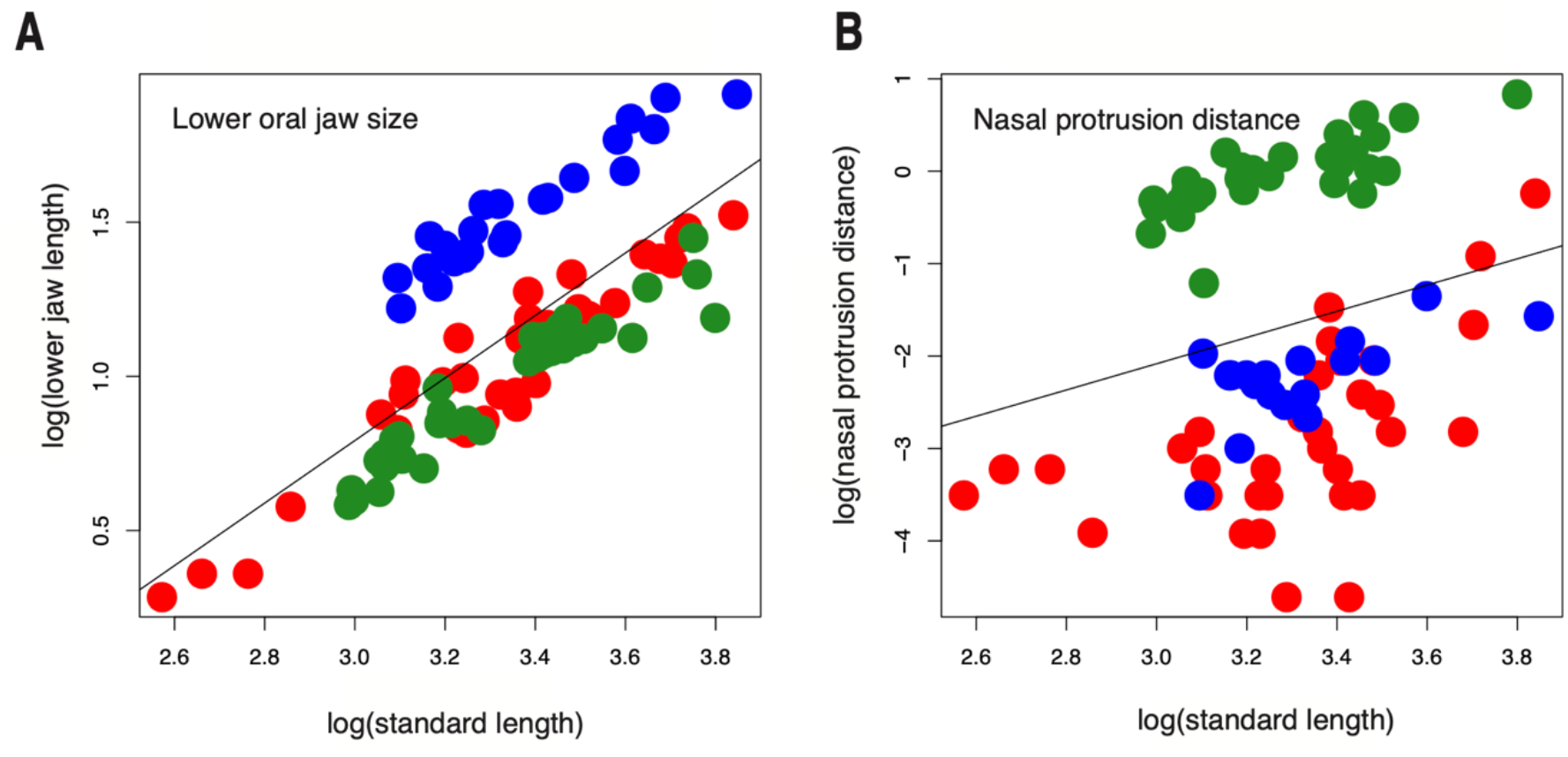
Standardized craniofacial trait measurements in SSI species. Log-transformed A) lower oral jaw length (mm) and B) nasal protrusion distance (mm) standardized by log-transformed standard length (mm) for SSI generalist (red), molluscivore (green), and scale-eater (blue).

**Fig. S10.**
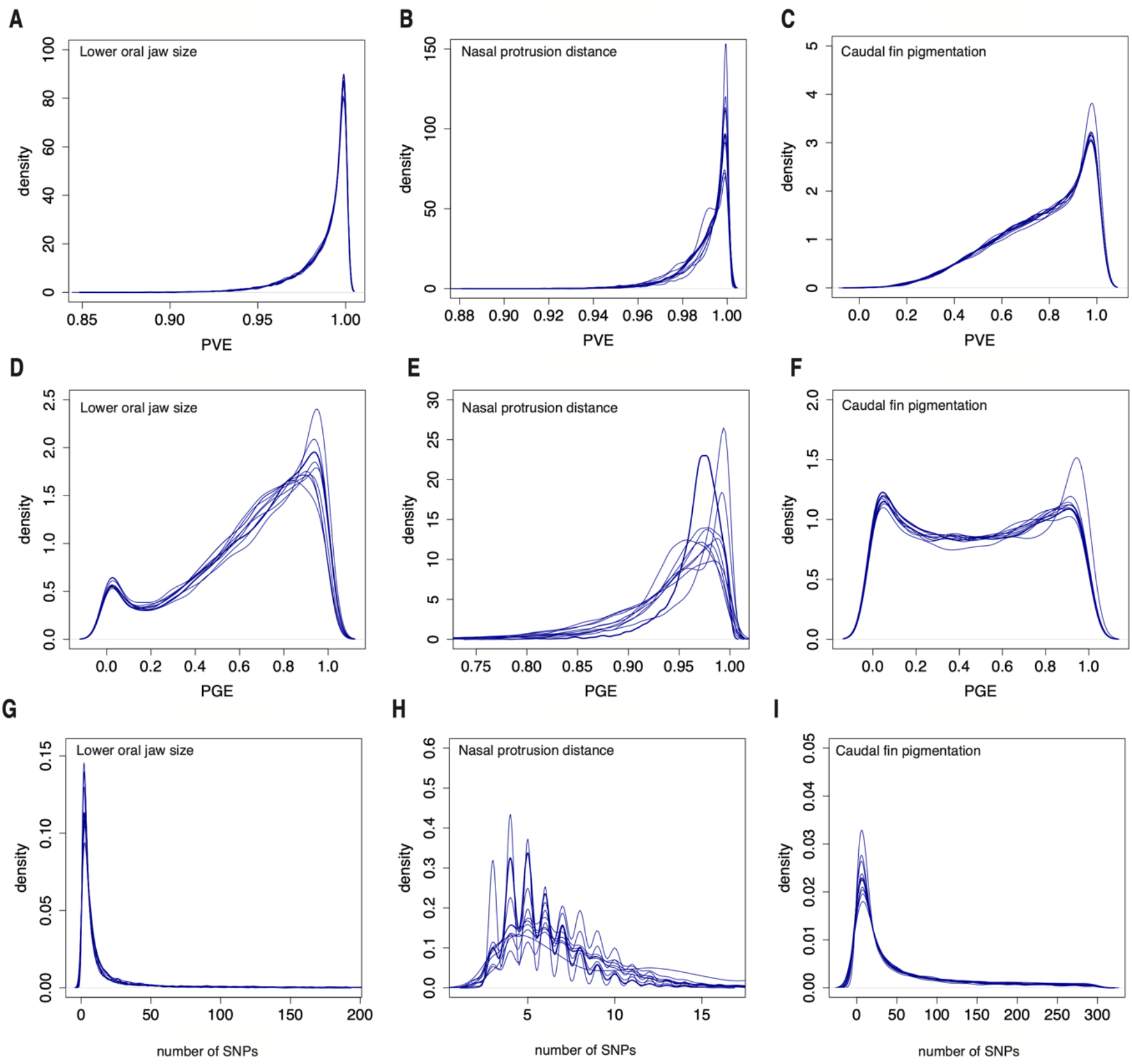
Posterior density distributions for hyper-parameters describing the proportion of variance in phenotypes for the three focal traits. (lower jaw size, nasal protrusion distance, and caudal fin pigmentation) explained by A-C) every SNP (proportion of phenotype variance explained (PVE)), D-F) SNPs of large effect (proportion of genetic variance explained by sparse effects) PGE)), and G-H) the number of large effect SNPS required to explain PGE. Individual lines represent 10 independent MCMC runs of GEMMA’s Bayesian sparse linear mixed model.

**Fig. S11.**
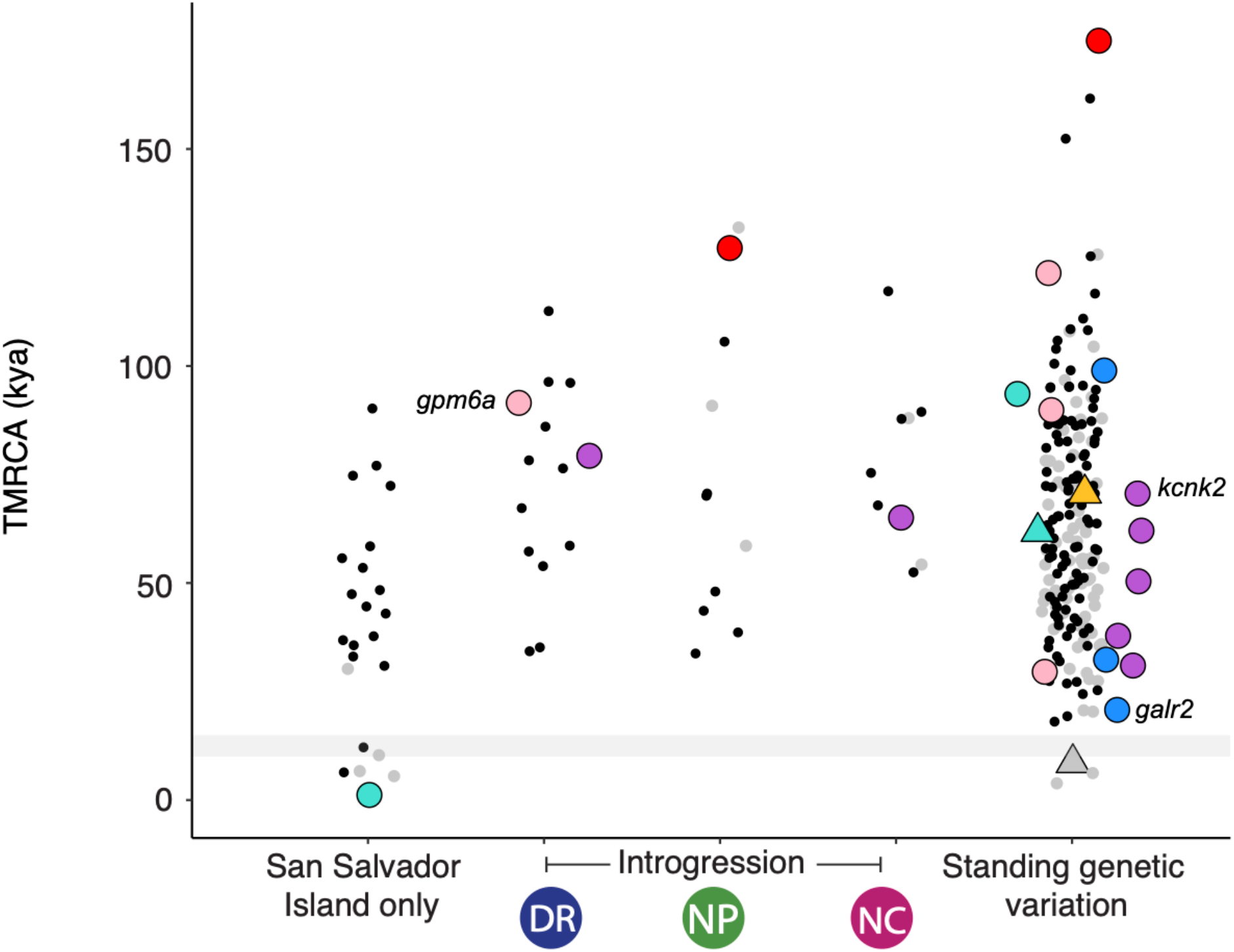
The spatiotemporal landscape of adaptive radiation from the perspective of the alternative allele (larger spatial distribution). Time to most recent common ancestor (TMRCA) of the region surrounding candidate adaptive variants (LD-pruned so that each point is independent) based on relative genetic divergence metric *Dxy* (*52*) which captures only the amount divergence that has accumulated since the two populations diverged. Variants are separated by spatial distribution into: de novo, introgressed from a single population and standing genetic variation observed in more than two populations. No nearly fixed variants introgressed from the Venezuela population. Points are colored by significantly enriched GO terms chosen for their adaptive relevance to this system (black points: adaptive variants annotated for non-focal GO terms or unannotated; gray points: nearly fixed variants between specialists that are not under hard selective sweeps) and triangle points represent those variants also associated with pigmentation. All variants annotated for GO categories from the Biological Processes database (Ensemble 92) for feeding behavior, muscle, eye, and mouth development, or craniofacial (mouth and/or eye, muscle) are shown. Points labeled with gene names indicate the three regions which have alternative spatial distributions shown in Fig. 4.

**Fig. S12.**
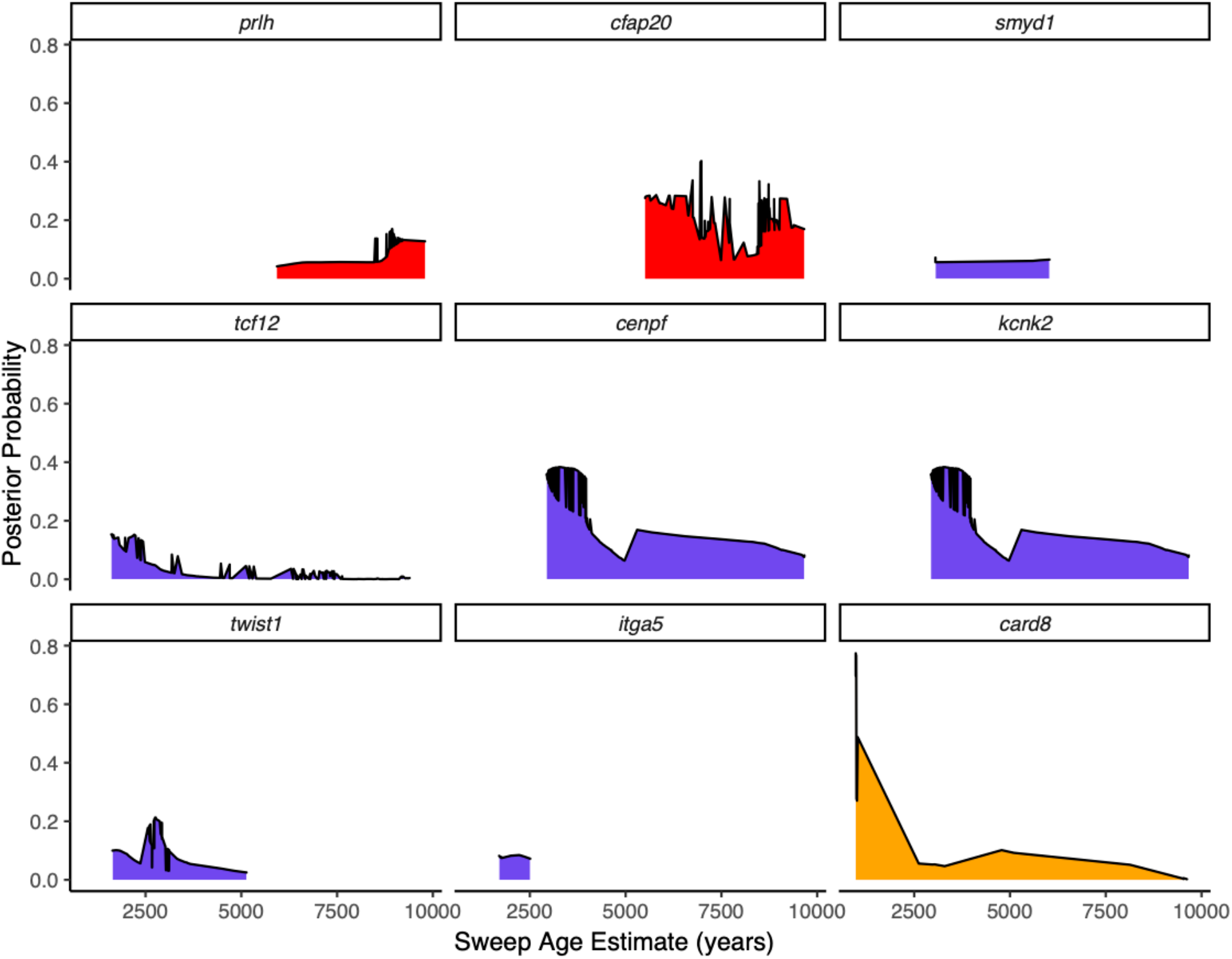
Posterior distributions for scale-eater sweeps. The posterior distributions of sweep ages estimated from focal regions (Table S13). These nine regions were found under a hard selective sweep using both SweeD and McSwan. Posterior distributions of ages were calculated in McSwan. Focal regions are colored based on GO and GWAS annotations relevant to the stages proposed in the stages of adaptive radiation hypothesis: habitat preference (feeding behavior: red), trophic morphology (craniofacial and muscle: blue-violet), and sexual communication (pigmentation: orange).

**Fig. S13.**
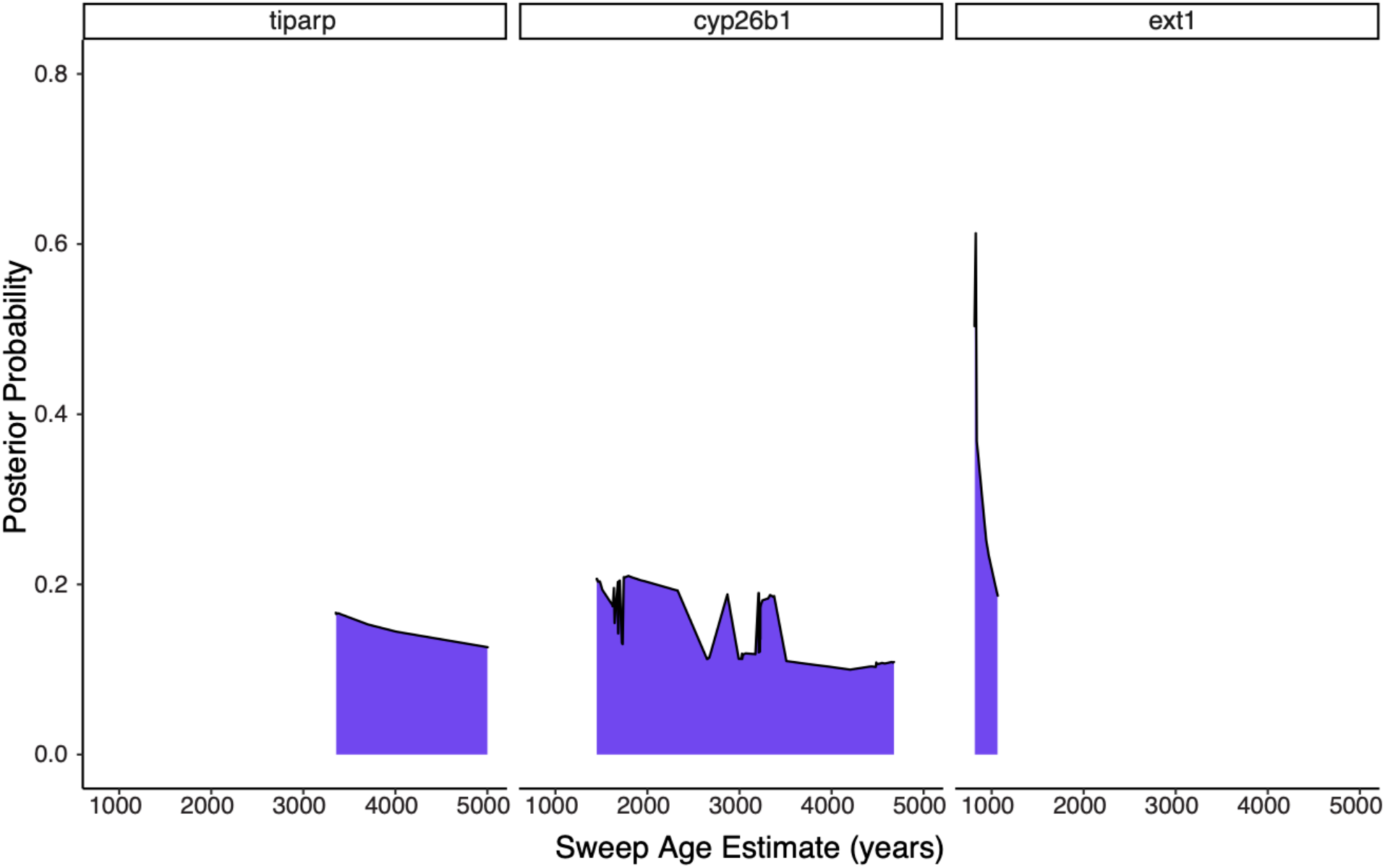
Posterior distributions for molluscivore sweeps. The posterior distributions of sweep ages estimated from focal regions (Table S13). These three regions were found under a hard selective sweep using both SweeD and McSwan. Posteriors distributions of ages were calculated in McSwan. Focal regions are colored based on GO and GWAS annotations relevant to the stages proposed in the stages of adaptive radiation hypothesis: trophic morphology (craniofacial and muscle: blue-violet).

**Table S1.**
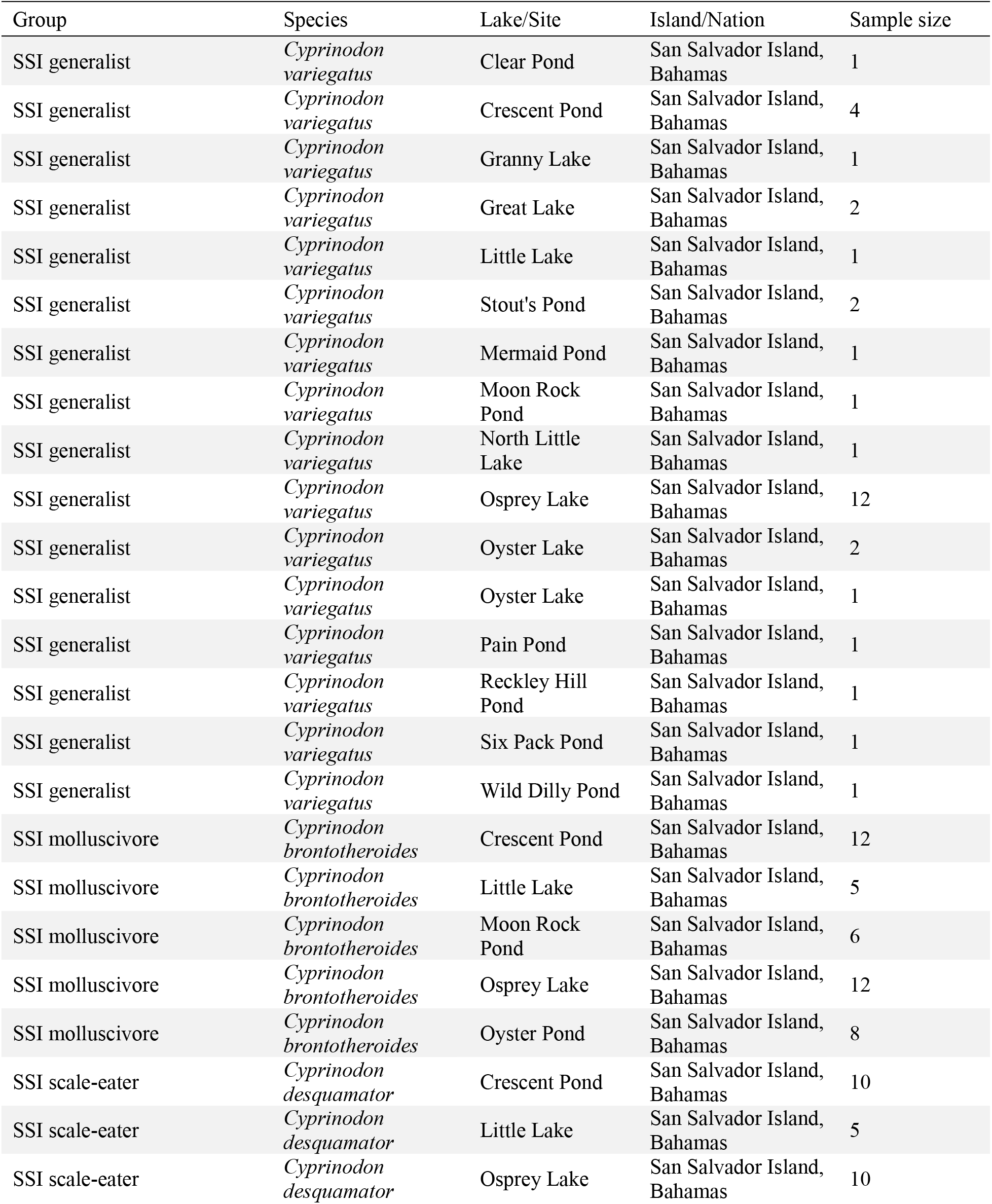
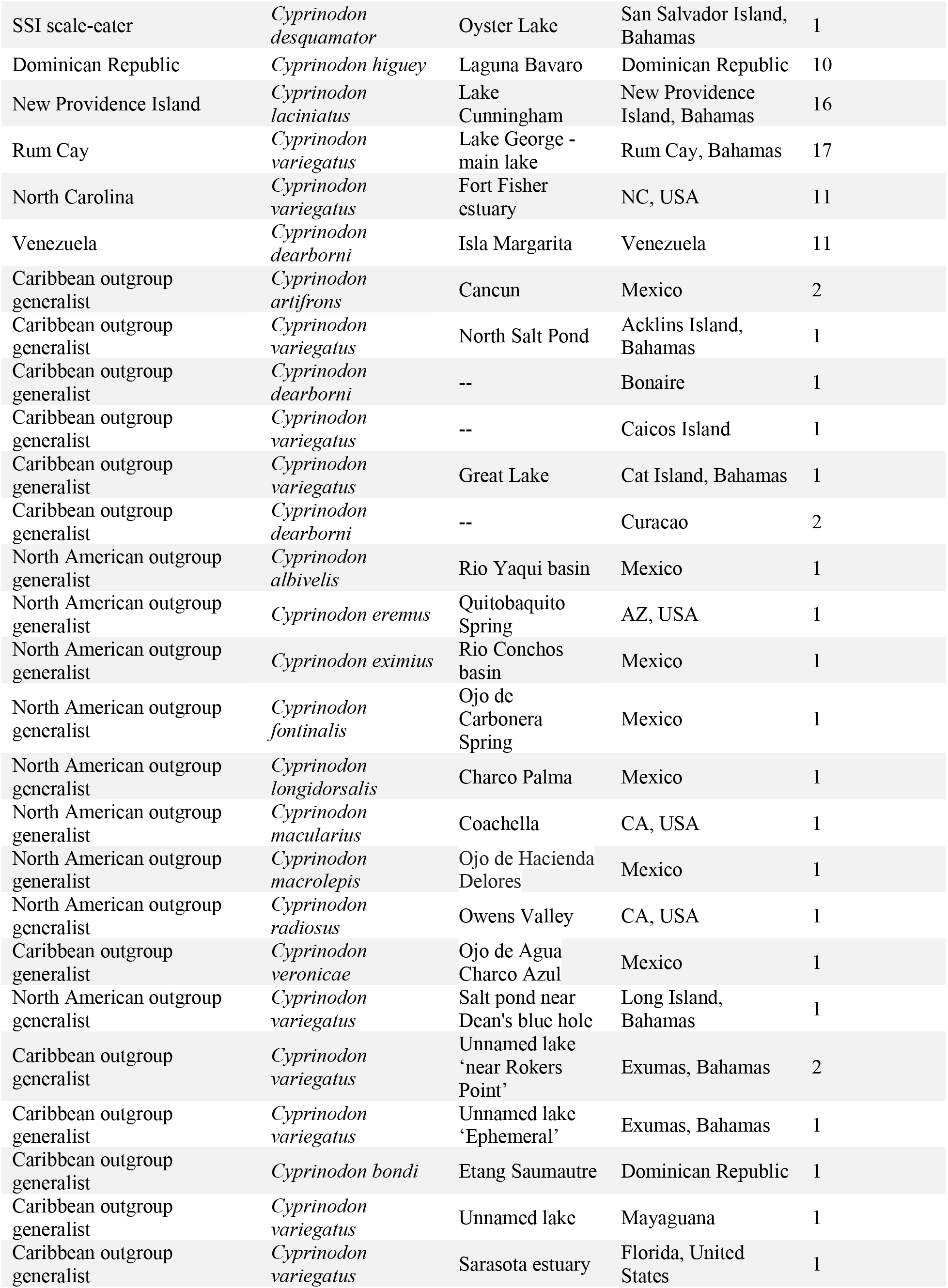
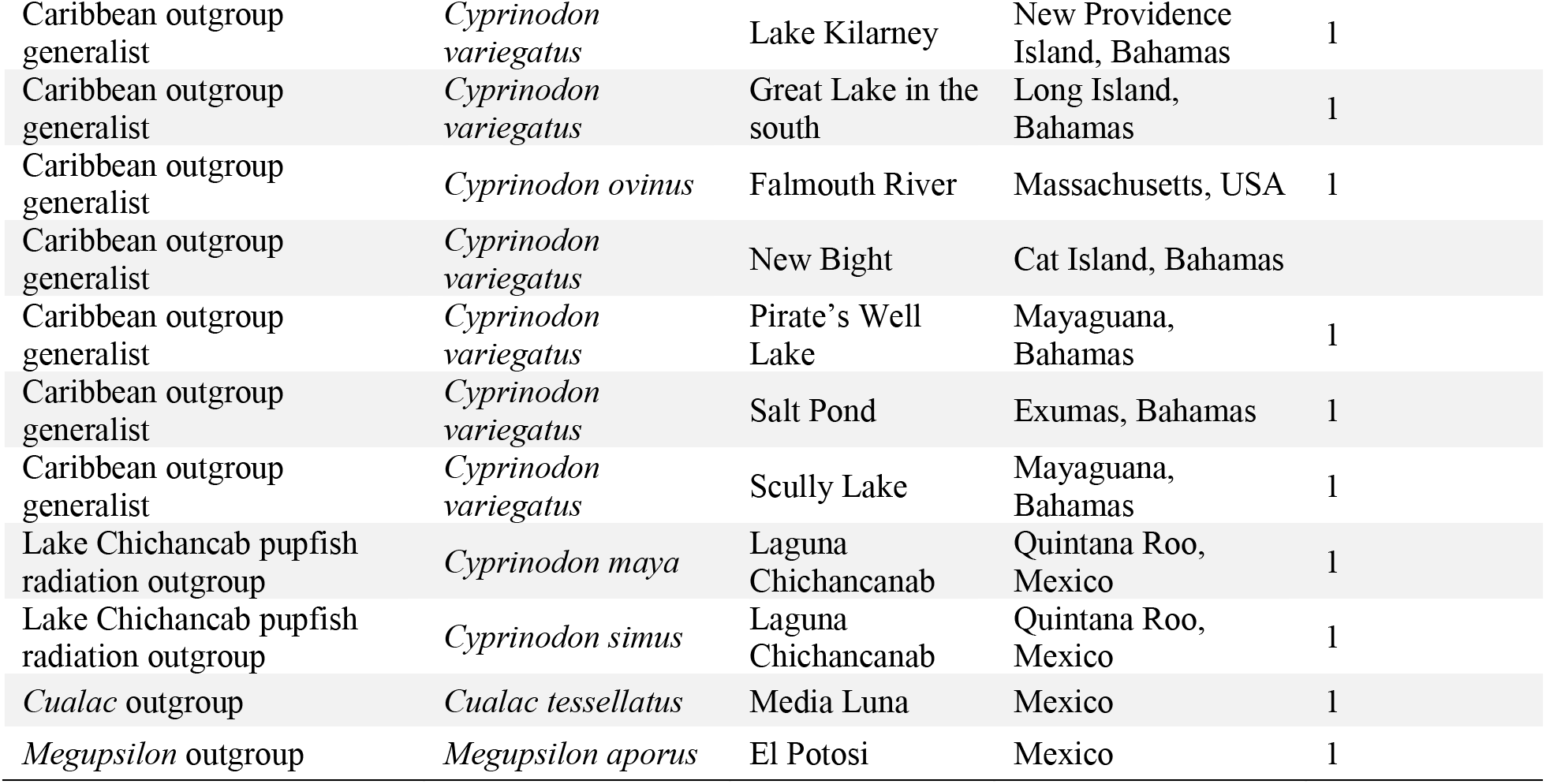
Summary of Caribbean pupfish sampling. The sampling localities of individuals that were whole genome resequenced from San Salvador Island radiation (SSI), other *Cyprinodon* across the Caribbean, Mexico, and United States, and two outgroups. Full details including sample codes, collector identities, GPS coordinates are included in Data S1 table.

**Table S2.**
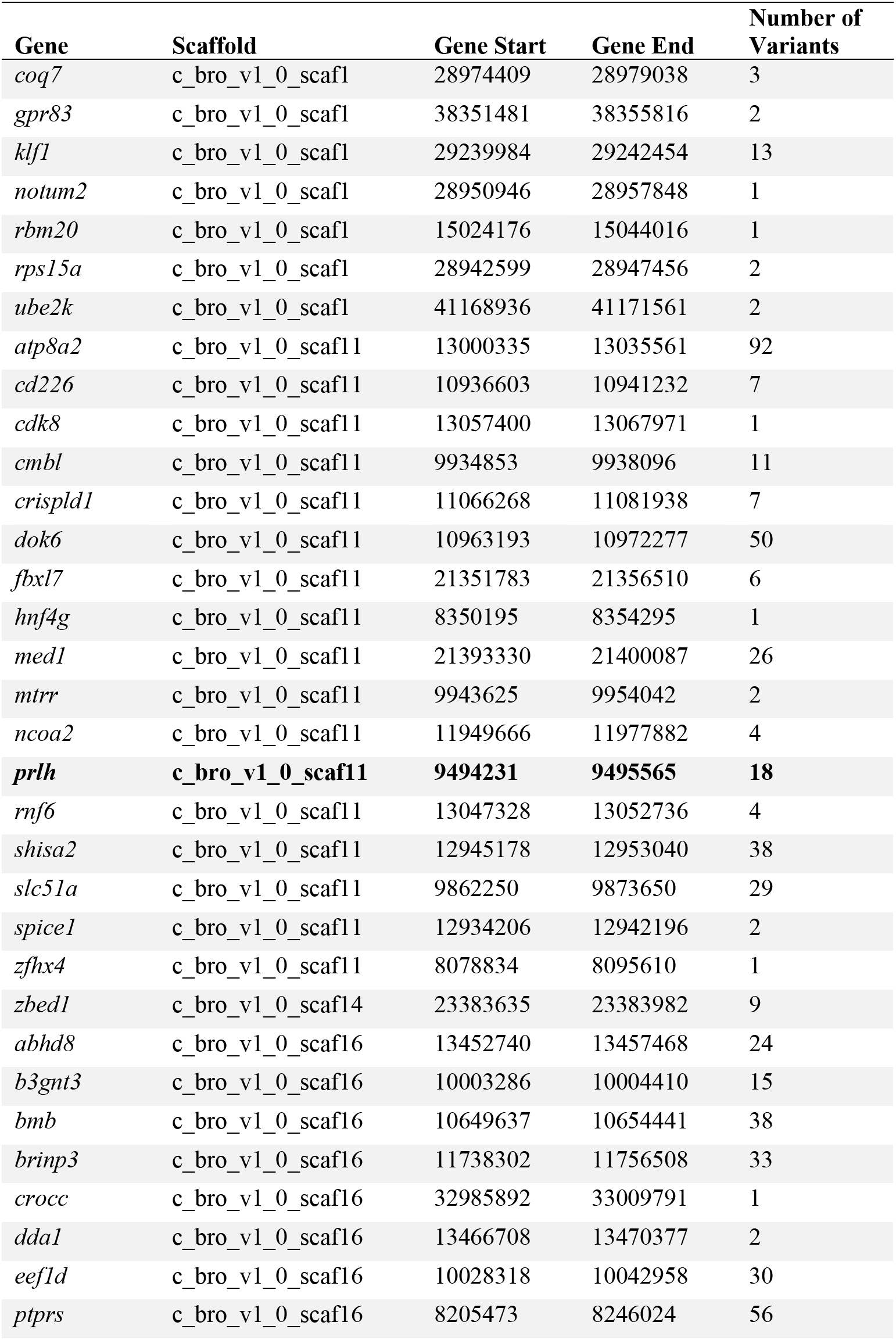
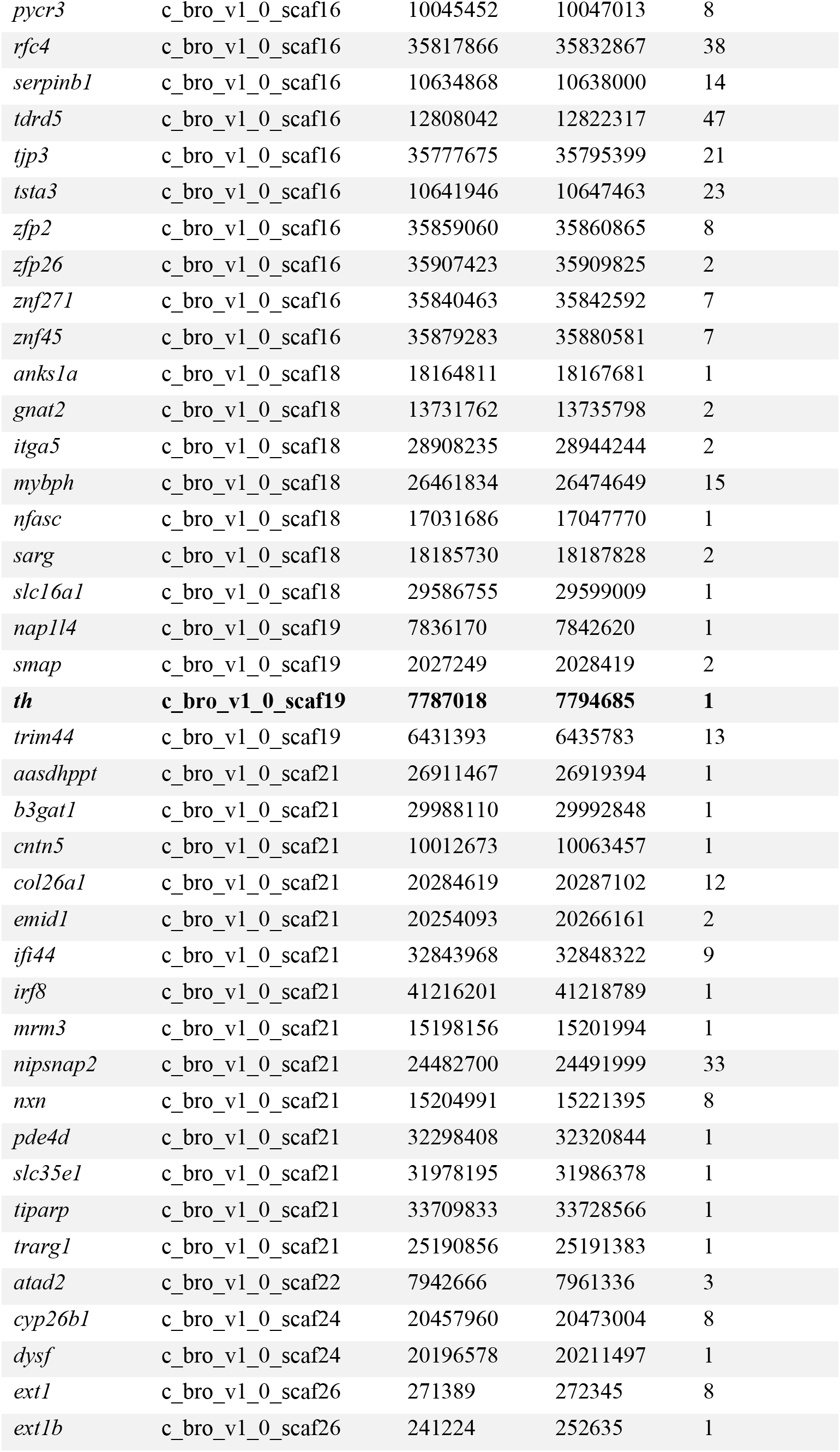
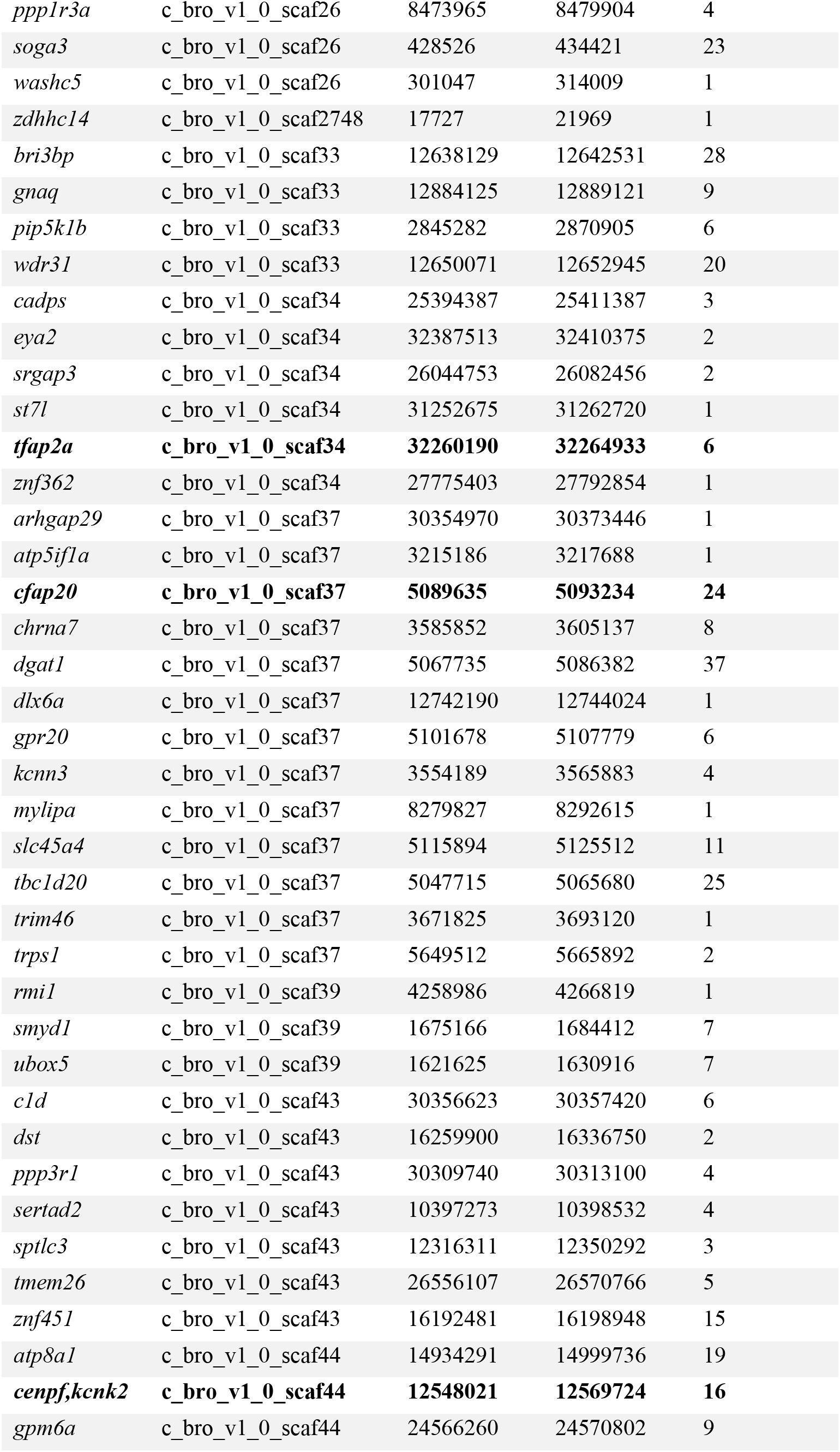
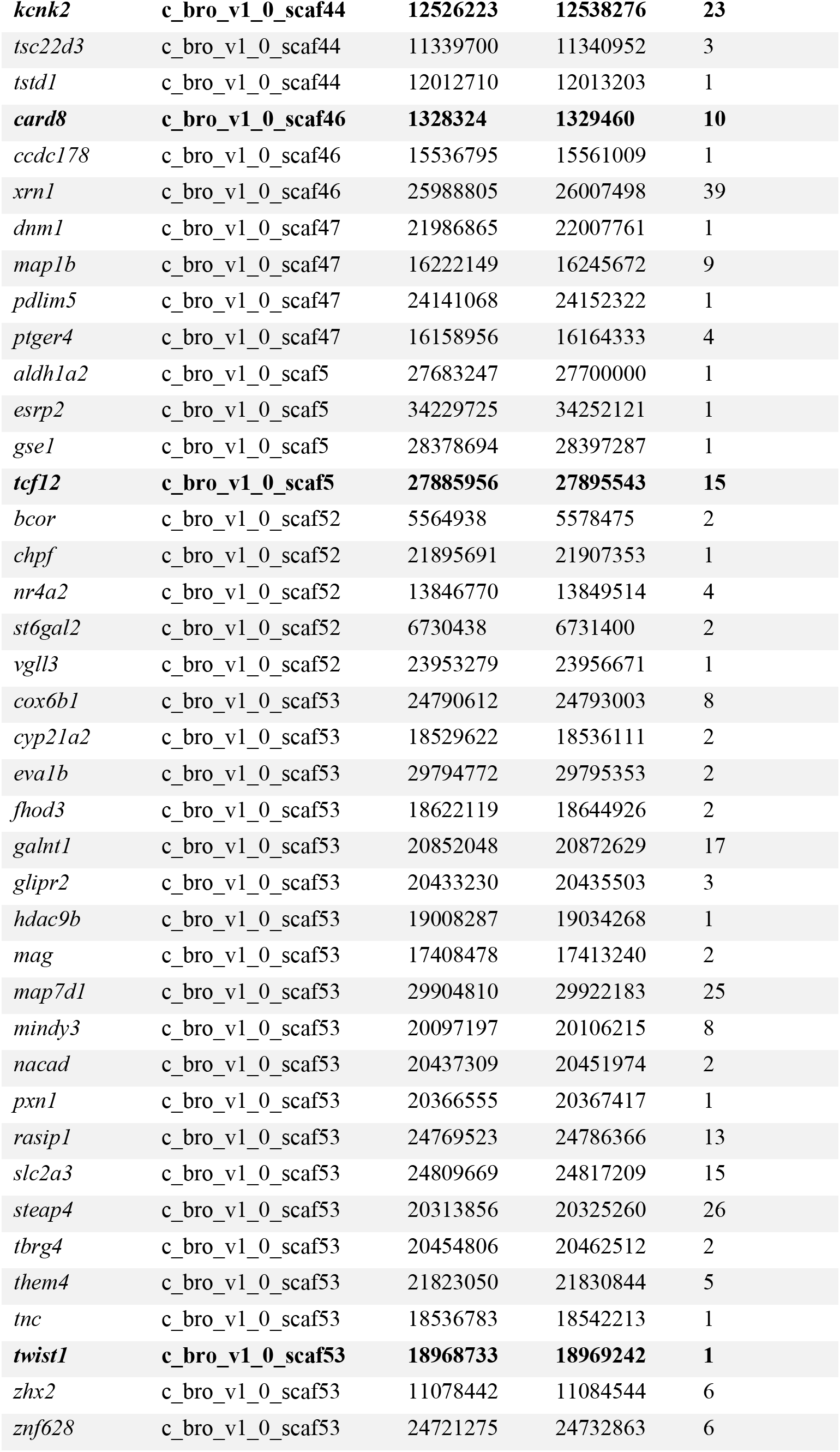
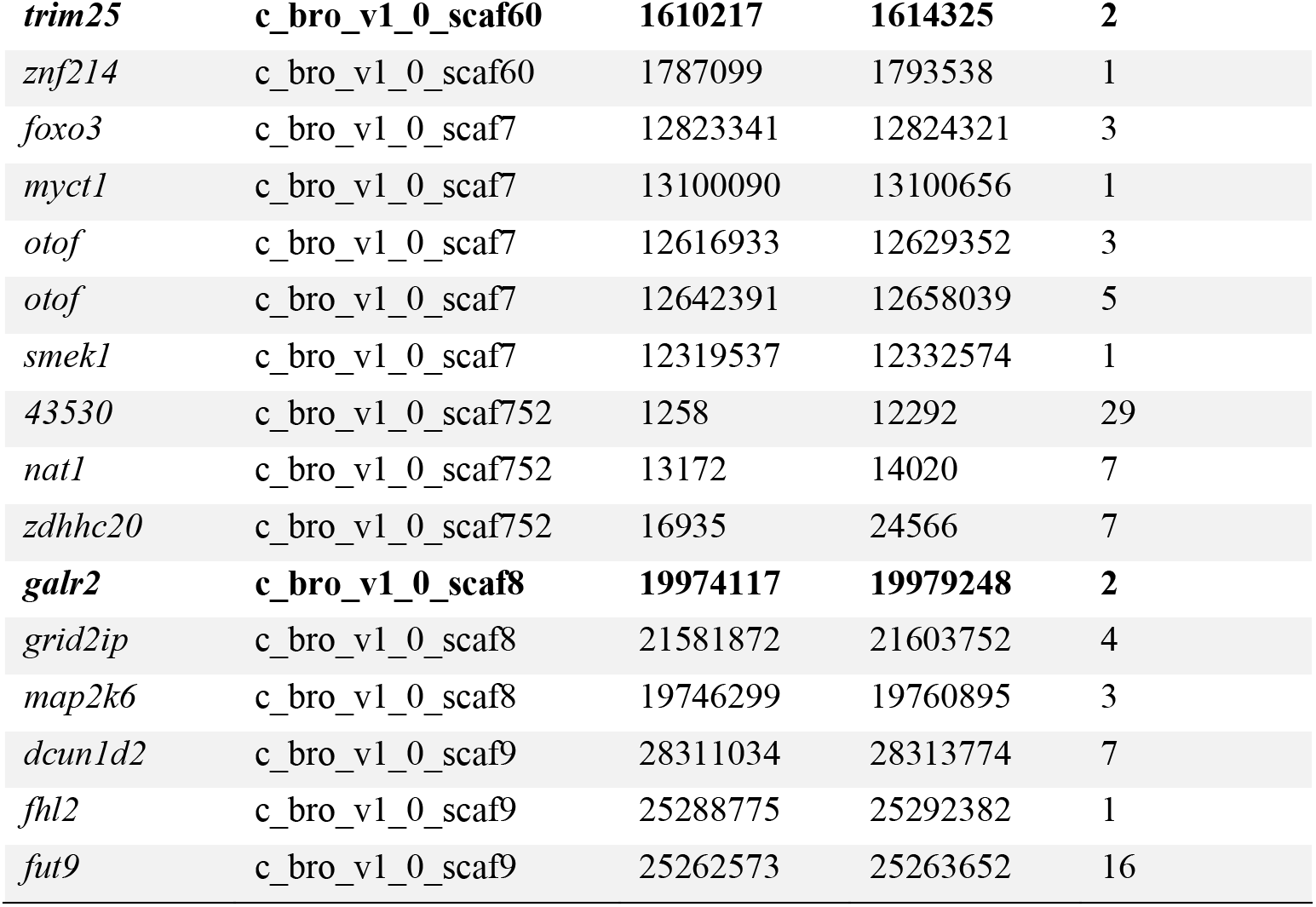
Candidate adaptive regions for the San Salvador Island scale-eater. Location of the genic regions that contained signatures of a strong selective sweep in the scale-eater (SweeD CLR ≥ 5.28) and at least one divergent variant between the specialists (*F_st_* ≥ 0.95). Full list of variants, including unannotated candidate regions provided in Data S2. Regions highlighted in Figure 4 are listed in bold.

**Table S3.**
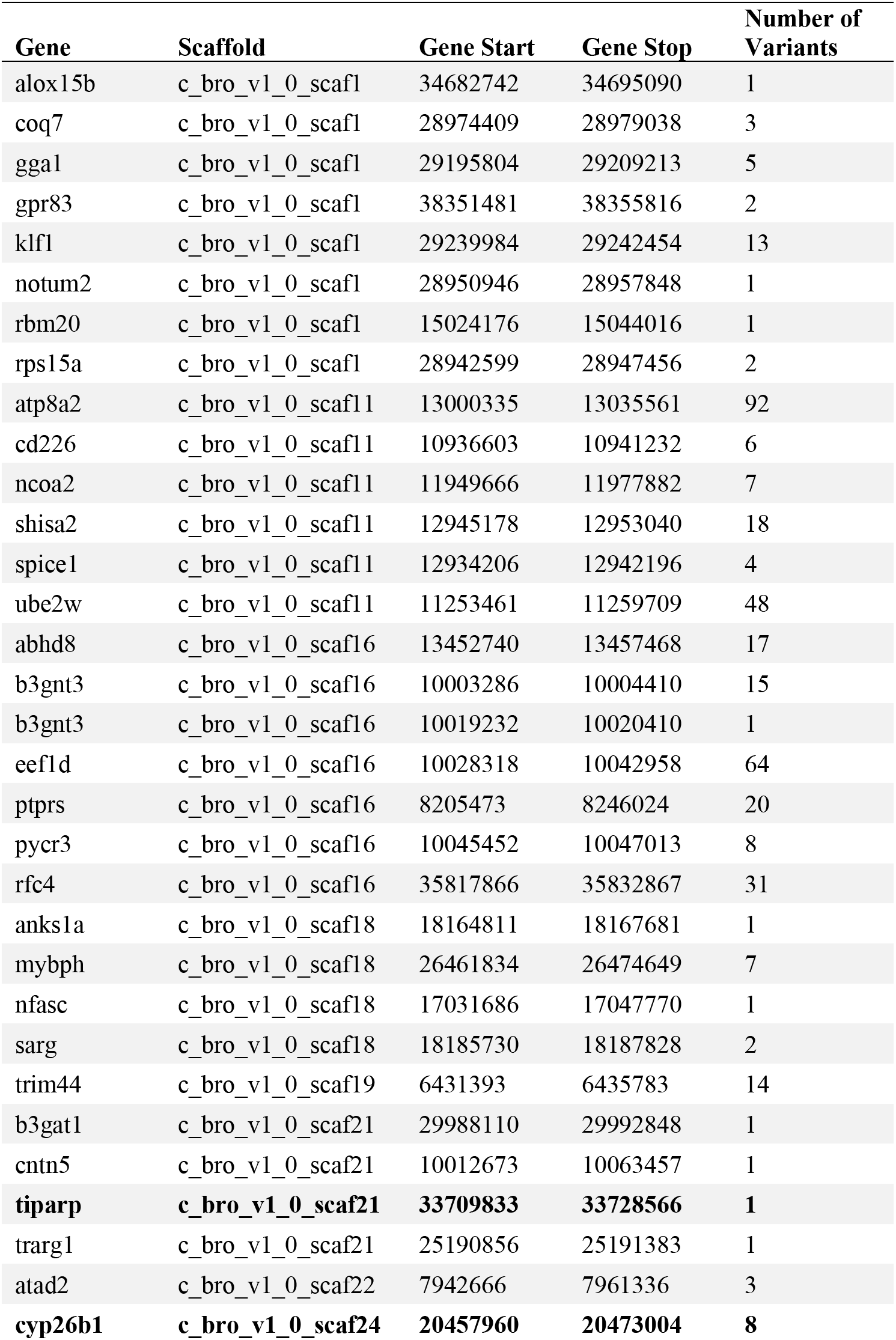
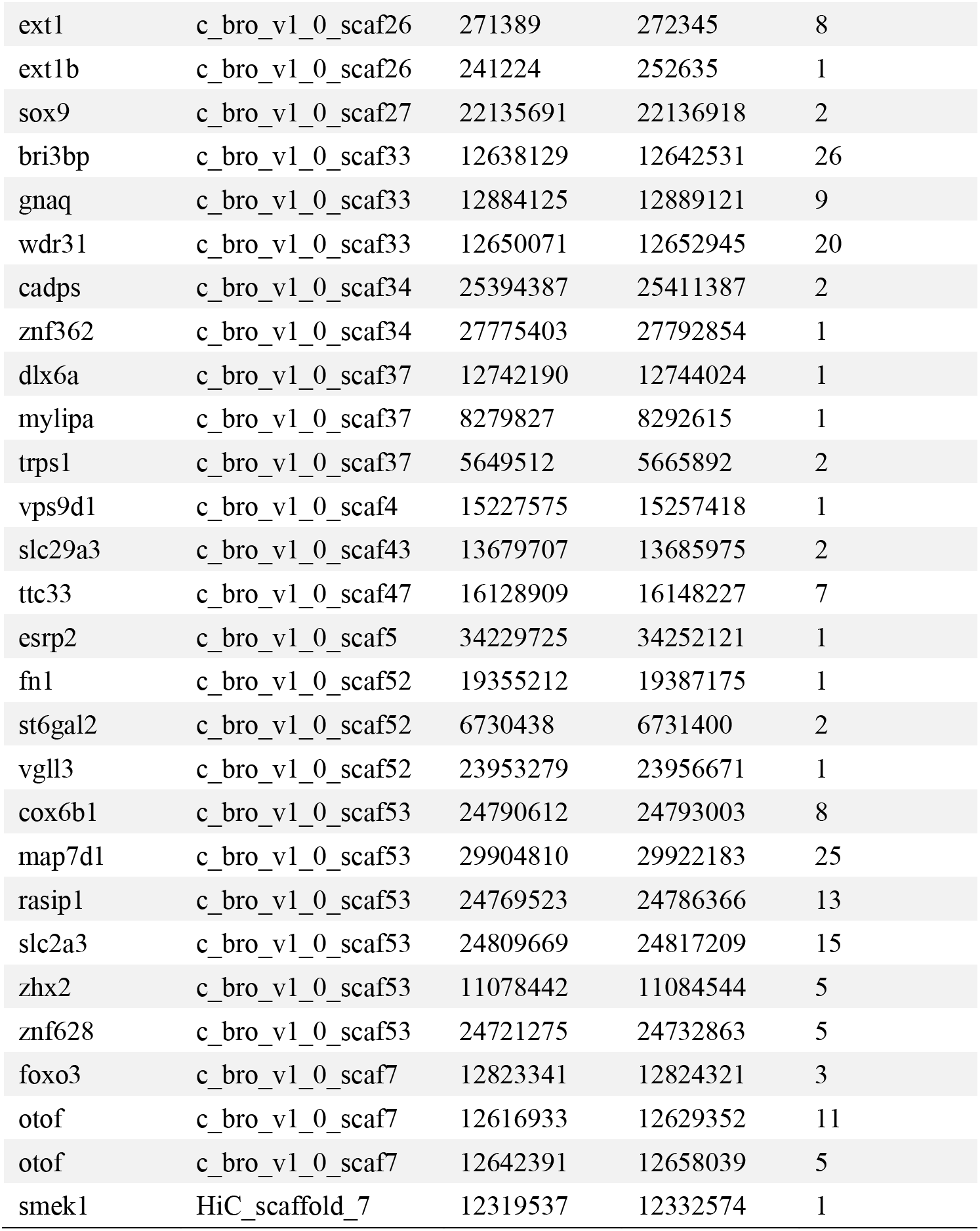
Candidate adaptive regions for the San Salvador Island molluscivore. Location of the genic regions that contained signatures of a strong selective sweep in the molluscivore (SweeD CLR ≥ 4.47) and at least one divergent variant between the specialists (*F_st_* ≥ 0.95). Full list of variants, including one unannotated candidate regions provided in Data S3. Regions highlighted in Figure S5 are listed in bold.

**Table S4.**
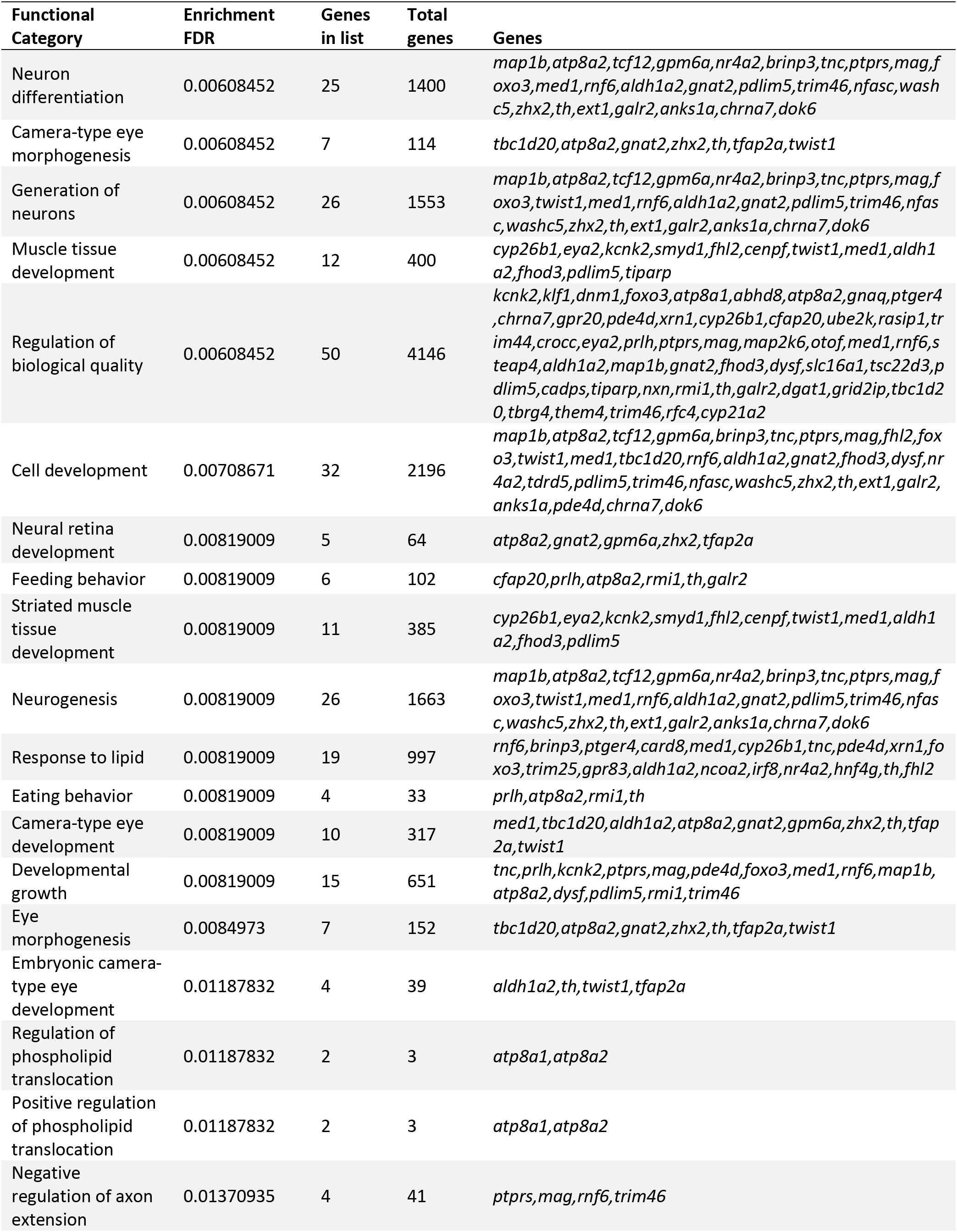
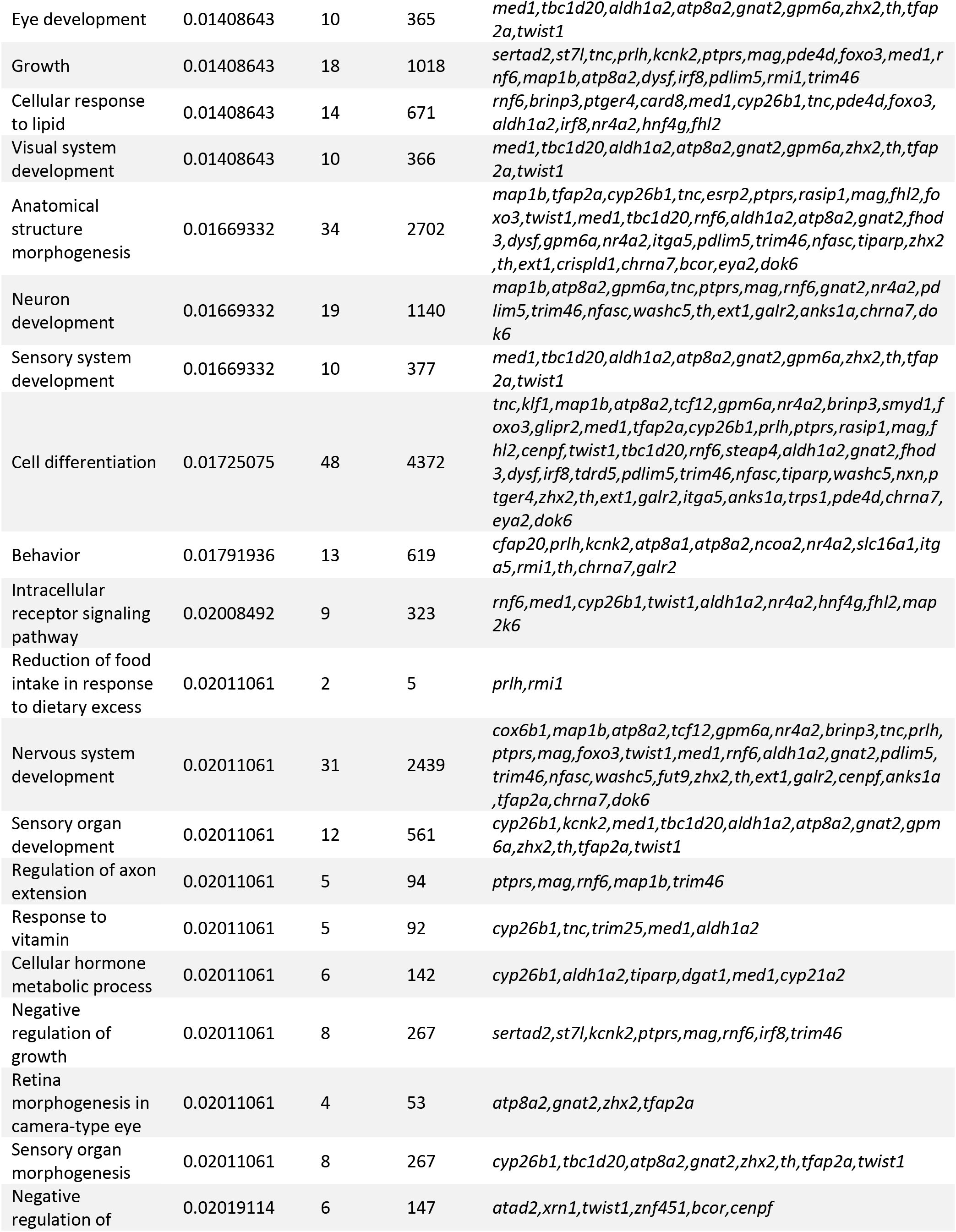
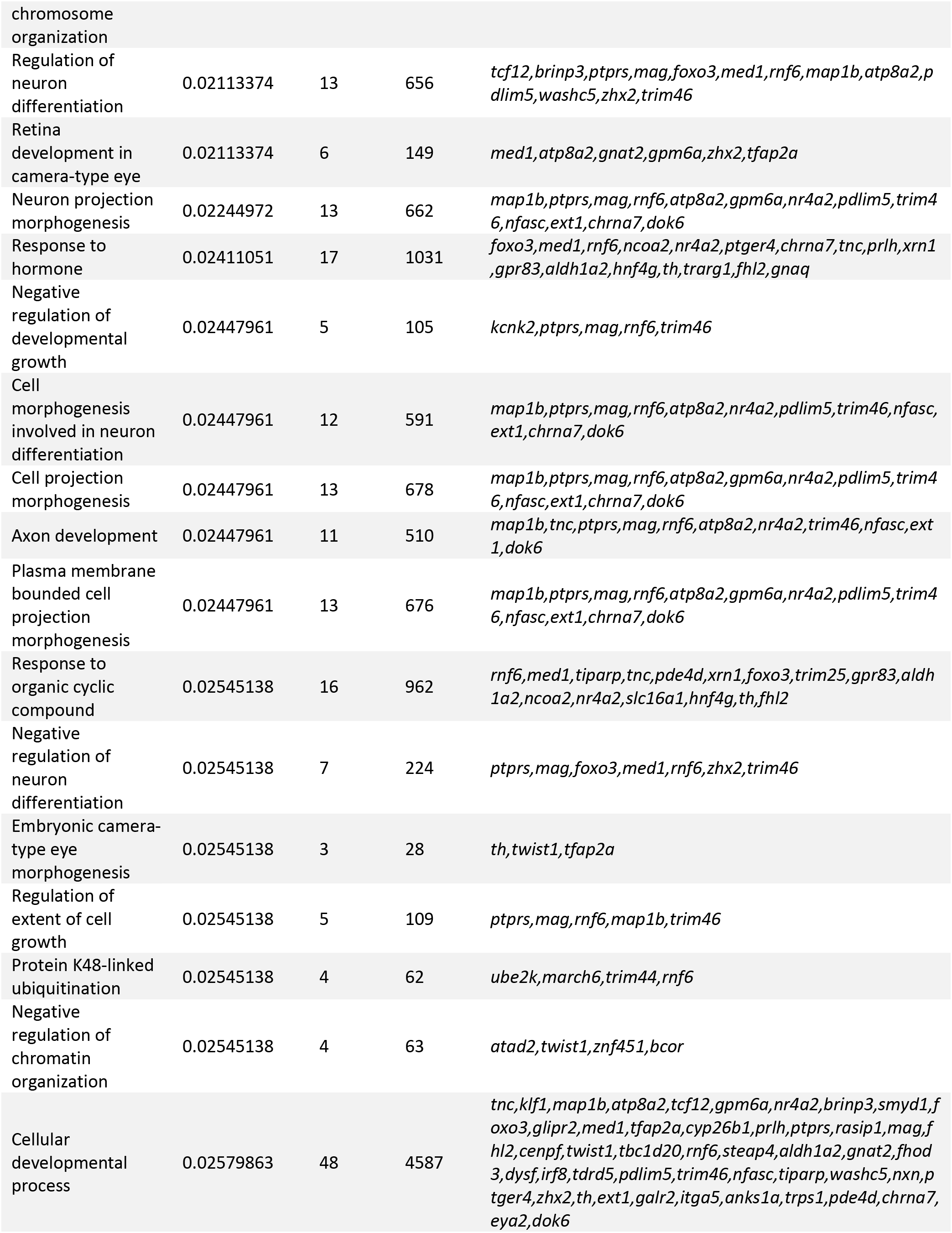
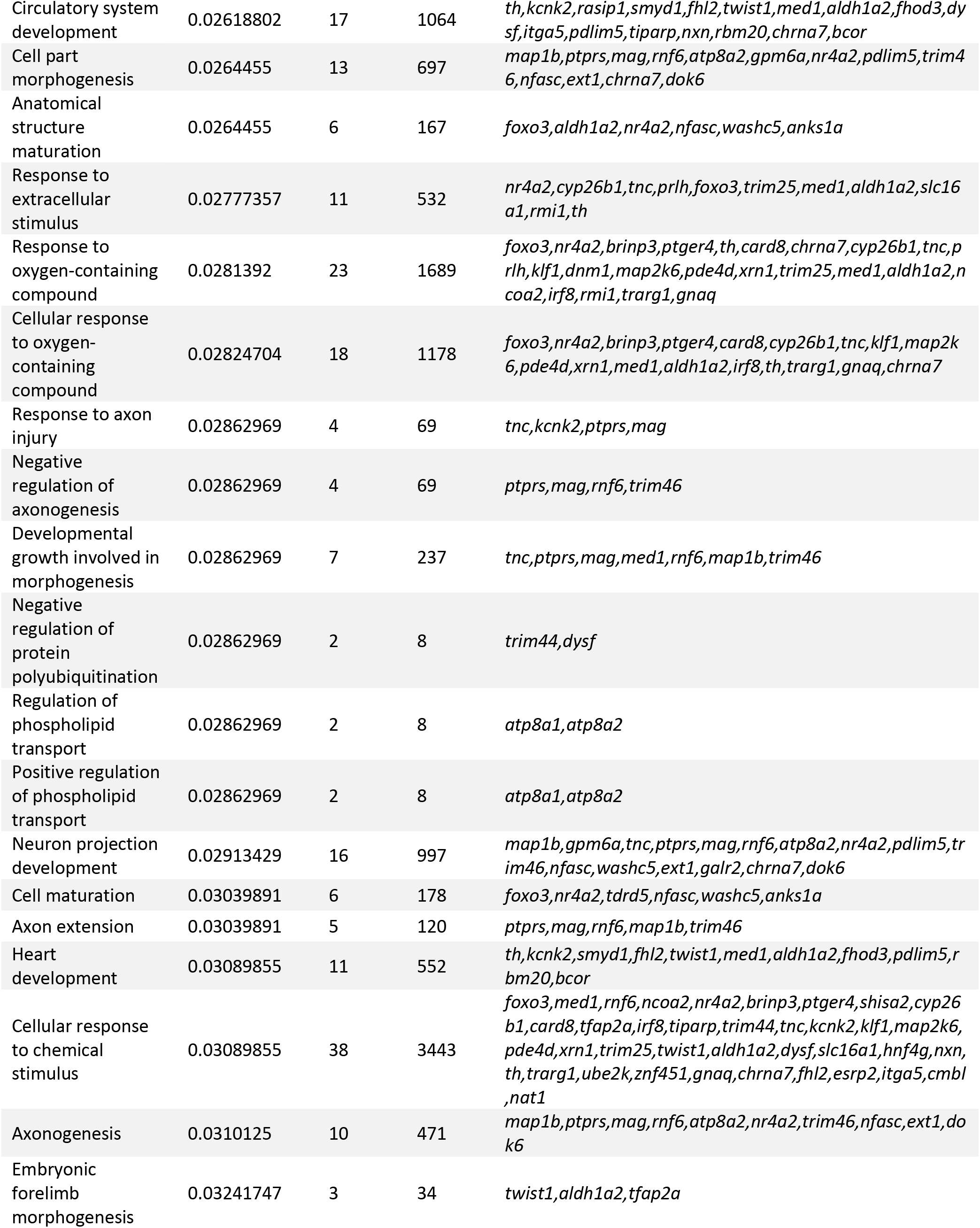
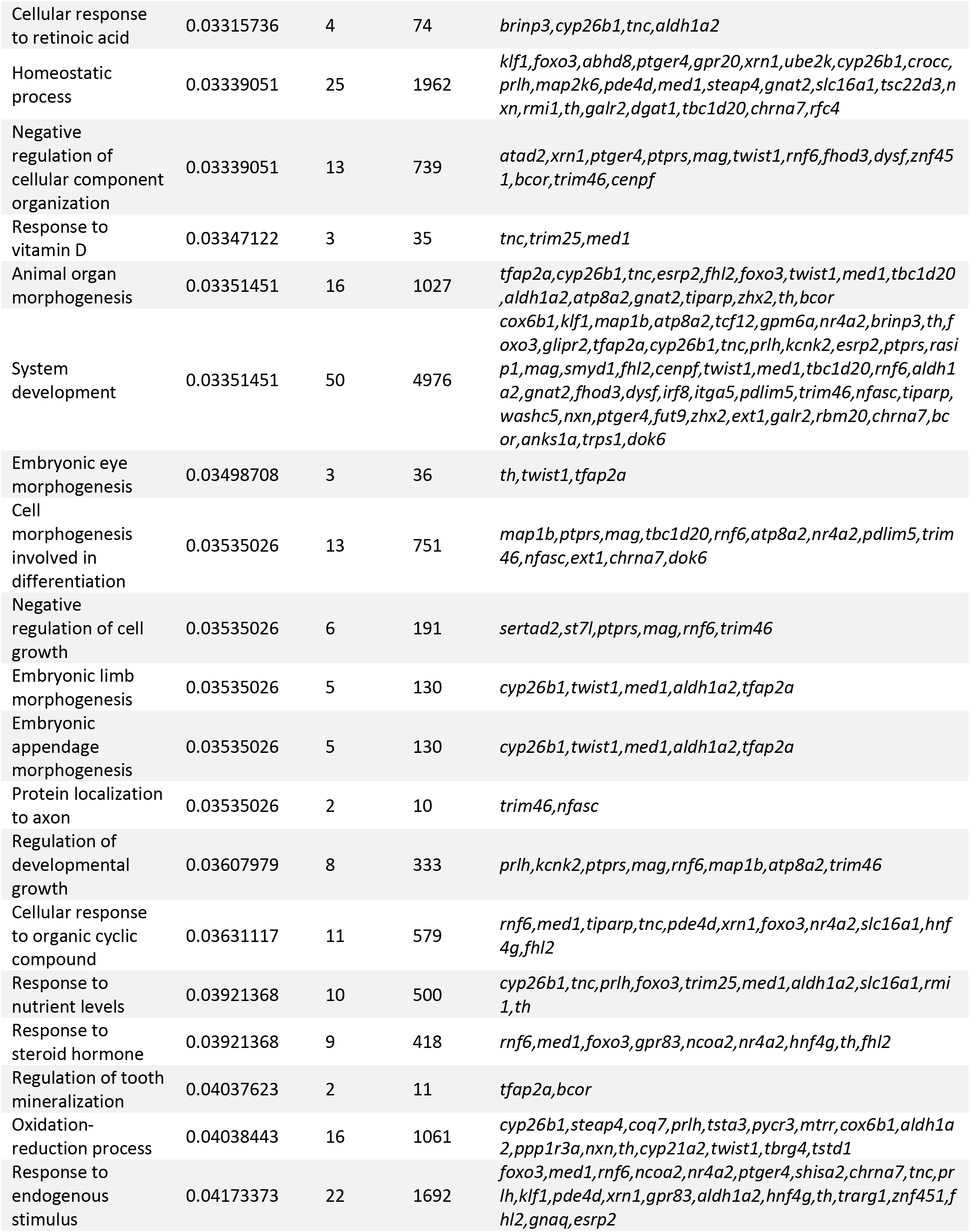
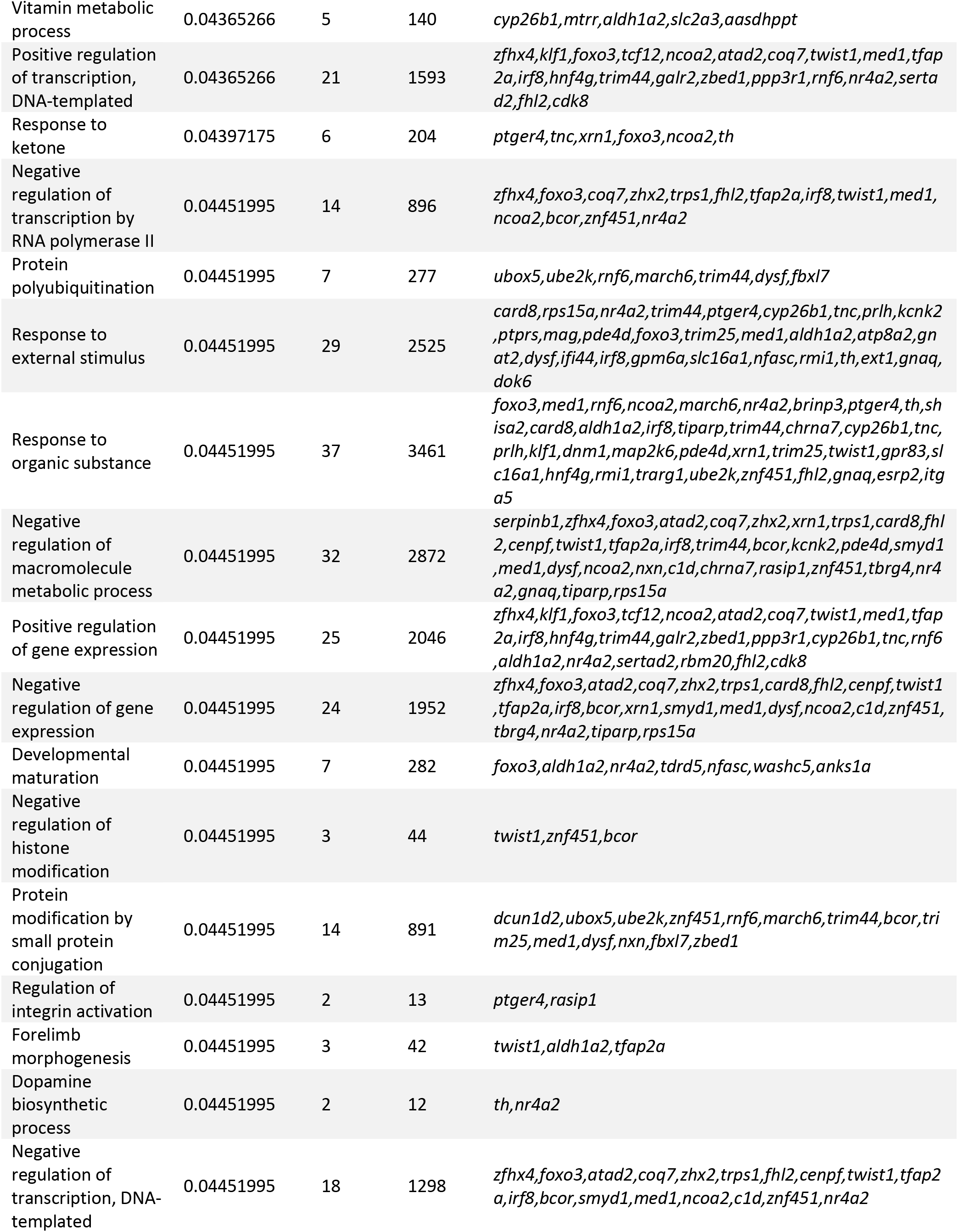
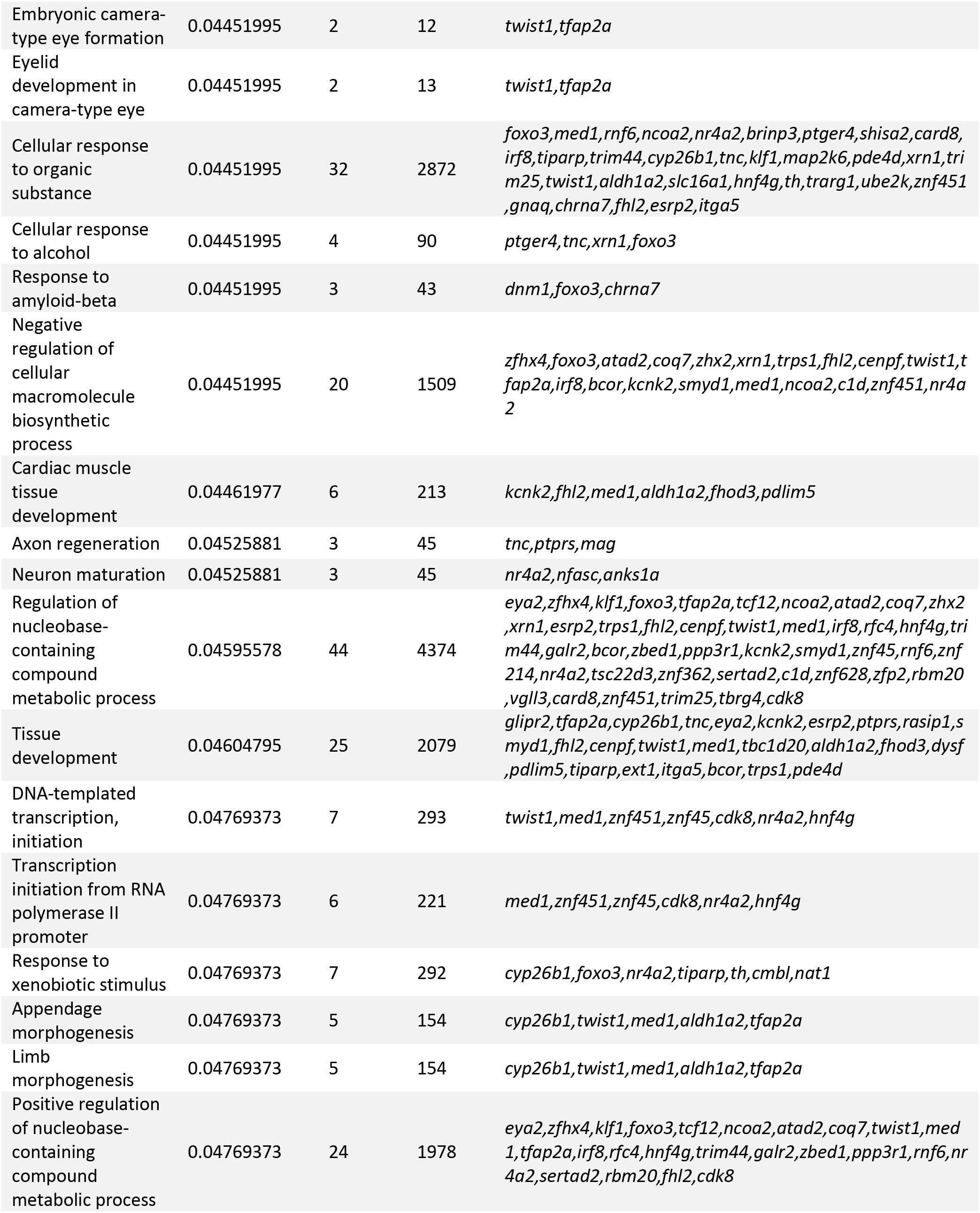
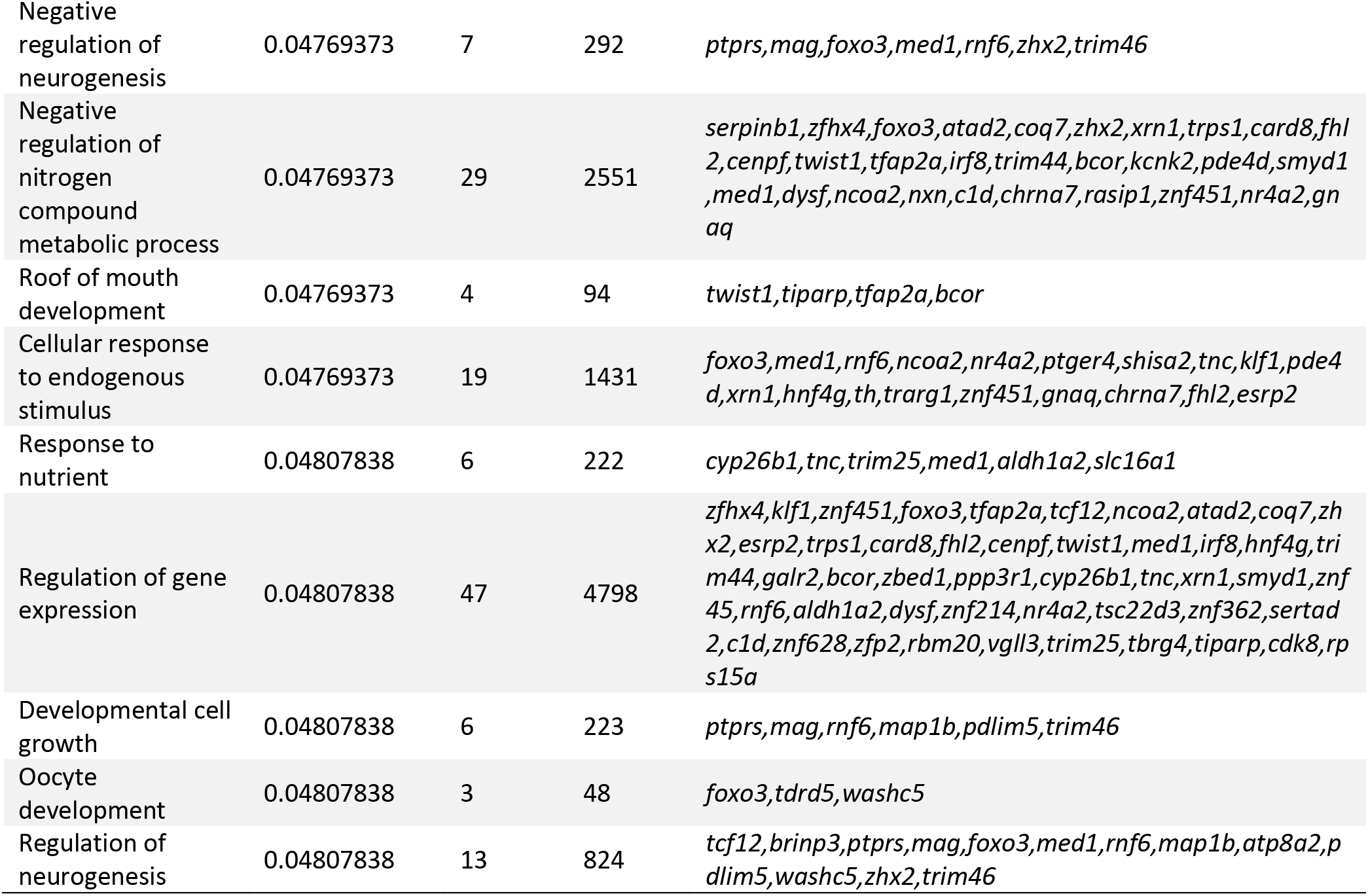
Full list of functional terms associated with genes in candidate regions for the scale-eaters that were significantly enriched (FDR <0.05) in a GO analysis. Focal functional terms related to key axes of diversification in this system: habitat preference (scale-eating/snail-eating niches), trophic morphology, and/or pigmentation.

**Table S5.** Top 5 BLAST hits for LG15 QTL. Bolded values indicate the top hit that was used to determine the region the significant oral jaw size QTL aligned to an 18-Mb region on scaffold c_bro_v1_0_scaf8 (8840660-27314762) in the *C. brontotheroides* reference genome that contained 3 genes (*map2k6*, *galr2*, and *grid2ip*).

| LG15<br>marker | Scaffold | %<br>identity | Length<br>(bp) | Mismatch | Start | End | E-value | Bitscore |
| --- | --- | --- | --- | --- | --- | --- | --- | --- |
| **10999** | **c_bro_v1_0_scaf8** | **97.917** | **96** | **2** | **8840660** | **8840755** | **1.95E-42** | **174** |
| 10999 | c_bro_v1_0_scaf8 | 100 | 17 | 0 | 17544438 | 17544422 | 4.6 | 34.2 |
| 10999 | c_bro_v1_0_scaf36 | 100 | 20 | 0 | 201747 | 201728 | 0.074 | 40.1 |
| 10999 | c_bro_v1_0_scaf7 | 100 | 19 | 0 | 13795738 | 13795756 | 0.29 | 38.2 |
| 10999 | c_bro_v1_0_scaf52 | 100 | 18 | 0 | 23185857 | 23185840 | 1.2 | 36.2 |
| 10999 | c_bro_v1_0_scaf38 | 100 | 18 | 0 | 1880370 | 1880387 | 1.2 | 36.2 |
| **33382** | **c_bro_v1_0_scaf8** | **100** | **93** | **0** | **27314670** | **27314762** | **1.97E-45** | **184** |
| 33382 | c_bro_v1_0_scaf8 | 93.617 | 47 | 3 | 26627380 | 26627426 | 8.05E-11 | 69.9 |
| 33382 | c_bro_v1_0_scaf8 | 93.617 | 47 | 3 | 27916662 | 27916616 | 8.05E-11 | 69.9 |
| 33382 | c_bro_v1_0_scaf8 | 95.238 | 42 | 2 | 1464518 | 1464477 | 3.18E-10 | 67.9 |
| 33382 | c_bro_v1_0_scaf8 | 95.238 | 42 | 2 | 11224060 | 11224019 | 3.18E-10 | 67.9 |

**Table S6.** Per generation mutation rate estimation from high coverage sequencing of parents and F1 from two crosses of San Salvador Island species.

| Cross | *C. variegatus* x<br>*C. brontotheroides* |  | *C. variegatus* x *C.*<br>*desquamator* |
| --- | --- | --- | --- |
| Offspring | F1.A | F1.B | F1.A |
| Avg. coverage | 67.5X | 45.1X | 32.7X |
| Known heterozygous sites genotype quality (GQ) | X>99 | X>99 | X>99 |
| Known heterozygous sites base quality rank sum (BaseQRankSum) | 1.4<x<2.6 | 1.4<x<2.6 | 1.4<x<2.7 |
| Known heterozygous sites mapping quality (MQ) | x>54 | x>54 | x>54 |
| Known heterozygous sites mapping quality rank sum (MQRankSum) | 1.6<x<1.9 | 1.6<x<1.9 | 1.4<x<2 |
| Known heterozygous sites quality by depth (QD) | 24<x<36 | 24<x<36 | 24<x<36 |
| Known heterozygous sites depth (DP) | 27 <x<77 | 15 < x< 54 | 12< x < 39 |
| Known heterozygous sites allele depth (AD) | 10<x<42 | 5<x<30 | 4< x < 21 |
| Known heterozygous sites read position rank sum (ReadPosRankSum) | -1.8<x<2.3 | 1.8<x<2.3 | 1.4<x<2.34 |
| Known heterozygous sites StrandOddsRatio (SOR) | 0.17<x<1.4 | 0.14<x<1.4 | 0.19<x<1.3 |
| Known heterozygous sites FisherStrand (FS) | 4.6<x<7.5 | 4.6<x<7.3 | 45<x<7.5 |
| GATK new mutation sites (bp) | 9114 | 8936 | 331 |
| mpileup new mutation sites (bp) | 14772 | 14182 | 7206 |
| Shared variants (bp) | 20 | 37 | 9 |
| Accessible genome (bp) | 698887016 | 712364816 | 695995433 |
| Mutation rate estimate | 1.43x10<sup>-8</sup> | 2.59x10<sup>-8</sup> | 6.46x10<sup>-9</sup> |

**Table S7.** Parameters for selective sweep analyses. The average coverage, composite likelihood ratio threshold based on neutral simulations, and the population size change parameters and individual used for each species.

| Species | Average Coverage | CLR threshold | SweeD Commands |
| --- | --- | --- | --- |
| SSI generalist | 28.87X | 4.89 | -folded -strictPolymorphic -G 0.4068 -eN 5.45 181.8 -s 64 |
| SSI molluscivore | 17.37X | 4.47 | -folded -strictPolymorphic -G 0.389 -eN 5.88 196 -s 88 |
| SSI scale-eater | 18.21X | 5.28 | -folded -strictPolymorphic -G 0.218 -eN 8.11 270 -s 52 |
| RC | 21.04X | 4.41 | -folded -strictPolymorphic -G 0.23 -eN 11.15 269.1 -s 34 |
| NP | 22.67X | 2.28 | -folded -strictPolymorphic -G 0.198 -eN 13.35 445.07 -s 30 |
| DR | NA | 5.37 | -folded -strictPolymorphic -G 0.236 -eN 10.83 362.8 -s 20 |
| NCC | 27.62X | 5.09 | -folded -strictPolymorphic -G 0.29 -eN 8.01 374.4 -s 24 |
| VEN | 17.21X | 18.05 | -folded -strictPolymorphic -G 8.87 -eN 0.086 0.345 -eN 1.077 38.78 -s 22 |

**Table S8.** The number of introgression regions in the SSI specialists. We determined introgressed regions of the genome as region with a *f_d_* statistic (ranges from 0 to 1) value above the threshold found in neutral simulations with no gene flow. These introgressed regions from each donor population were then overlapped with regions of the genome with strong genetic divergence (variants with Fst >= 0.95) and signatures of a hard selective sweep (CLR > 5.28 and > 4.47 for scale-eaters and molluscivores respectively) to determine the number of adaptive introgression regions. These adaptive introgression regions range in size from 50-kb to 110-kb in length.

| Donor population (P3) | $f_d$ threshold | Number of candidate introgression regions | Number of candidate adaptive introgression regions |
| --- | --- | --- | --- |
| Introgression with Molluscivore |  |  |  |
| Rum Cay | 0.81 | 536 | 5 |
| New Providence | 0.72 | 660 | 7 |
| Dominican Republic | 0.81 | 375 | 8 |
| North Carolina | 0.69 | 138 | 0 |
| Venezuela | 0.69 | 54 | 0 |
| Introgression with Scale-eater |  |  |  |
| Rum Cay | 0.81 | 385 | 5 |
| New Providence | 0.72 | 645 | 9 |
| Dominican Republic | 0.81 | 426 | 11 |
| North Carolina | 0.71 | 163 | 3 |
| Venezuela | 0.69 | 15 | 0 |

**Table S9.**
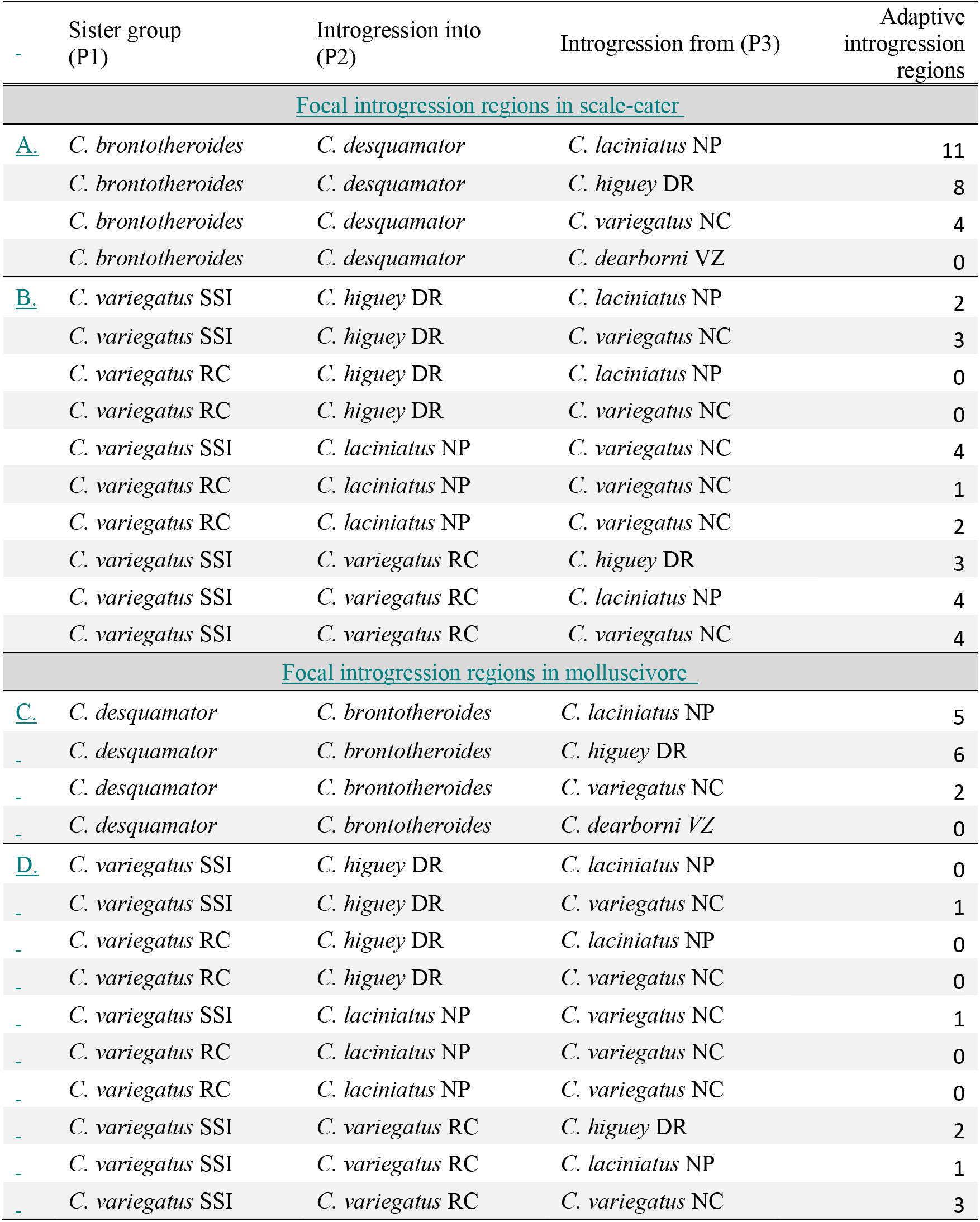
Combinations of Caribbean pupfish populations used to detect signatures of introgression in San Salvador Island specialists and generalist lineages on other islands. The *f_d_* statistic was used to detect introgression between combinations of P2 and P3 populations, given the tree (((P1,P2),P3),O). For this series of tests we used *C. artifrons* as the outgroup in which limited gene flow is expected to have occurred with the others.

**Table S10.** Candidate adaptive introgression regions from Rum Cay generalists (*C. variegatus*) and San Salvador Island specialists.

| Scaffold | Variant<br>Position | Start | End | Gene |
| --- | --- | --- | --- | --- |
| Introgression with Molluscivore |  |  |  |  |
| c_bro_v1_0_scaf11 | 12962909 | 12965001 | 13010000 | *shisa2, atp8a2* |
| c_bro_v1_0_scaf16 | 35813565 | 35765001 | 35875000 | *rfc4* |
| c_bro_v1_0_scaf18 | 18167642 | 18150001 | 18215000 | *anks1a* |
| c_bro_v1_0_scaf18 | 18177499 | 18150001 | 18225000 | *sarg* |
| c_bro_v1_0_scaf52 | 19358574 | 19345001 | 19395000 | *fn1* |
| Introgression with Scale-eater |  |  |  |  |
| c_bro_v1_0_scaf1 | 15017907 | 14995001 | 15065000 | *rbm20* |
| c_bro_v1_0_scaf5 | 28411973 | 28365001 | 28455000 | *gse1* |
| c_bro_v1_0_scaf37 | 3586373 | 3585001 | 3650000 | *chrna7* |
| c_bro_v1_0_scaf43 | 30358142 | 30355001 | 30405000 | *c1d* |
| c_bro_v1_0_scaf53 | 11080970 | 11080001 | 11130000 | *zhx2* |

**Table S11.** Candidate adaptive introgression regions from Dominican Republic generalists (C. higuey) and San Salvador Island specialists.

| Scaffold | Variant Position | Start | End | Gene |
| --- | --- | --- | --- | --- |
| Introgression with Molluscivore |  |  |  |  |
| c_bro_v1_0_scaf1 | 28938769 | 28935001 | 28985000 | *rps15a* |
| c_bro_v1_0_scaf1 | 28962108 | 28935001 | 28995000 | *notum2* |
| c_bro_v1_0_scaf1 | 28969771 | 28935001 | 28995000 | *coq7* |
| c_bro_v1_0_scaf7 | 12326193 | 12305001 | 12375000 | *smek1* |
| c_bro_v1_0_scaf7 | 12606143 | 12605001 | 12685000 | *otof* |
| c_bro_v1_0_scaf11 | 11256440 | 11210001 | 11295000 | *ube2w* |
| c_bro_v1_0_scaf18 | 18167642 | 18135001 | 18225000 | *anks1a,sarg* |
| c_bro_v1_0_scaf19 | 6430544 | 6410001 | 6465000 | *trim44* |
| Introgression with Scale-eater |  |  |  |  |
| c_bro_v1_0_scaf5 | 28411973 | 28385001 | 28450000 | *gse1* |
| c_bro_v1_0_scaf8 | 19759133 | 19735001 | 19790000 | *map2k6* |
| c_bro_v1_0_scaf18 | 28961523 | 28915001 | 29010000 | *itga5* |
| c_bro_v1_0_scaf19 | 7822448 | 7815001 | 7870000 | *nap1l4* |
| c_bro_v1_0_scaf34 | 25414453 | 25400001 | 25460000 | *cadps* |
| c_bro_v1_0_scaf34 | 26069290 | 26020001 | 26115000 | *srgap3* |
| c_bro_v1_0_scaf37 | 3700741 | 3685001 | 3750000 | *trim46* |
| c_bro_v1_0_scaf44 | 12541185 | 12540001 | 12620000 | *kcnk2, cenpf* |
| c_bro_v1_0_scaf44 | 24564920 | 24540001 | 24620000 | *gpm6a* |
| c_bro_v1_0_scaf53 | 18998120 | 18990001 | 19045000 | *hdac9b* |
| c_bro_v1_0_scaf53 | 20294941 | 20245001 | 20330000 | *steap4* |

**Table S12.**
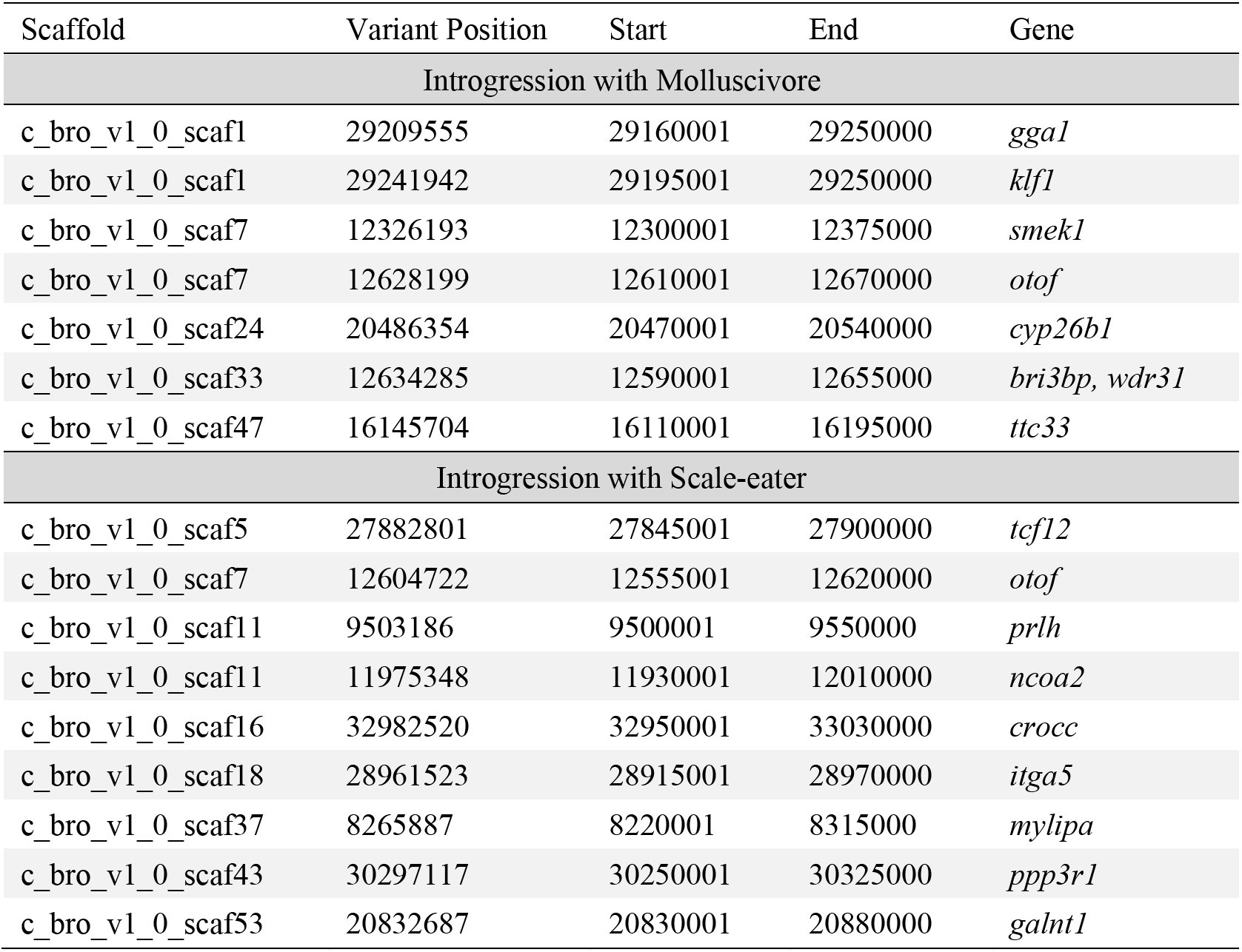
Candidate adaptive introgression regions from New Providence Island generalists (*C. laciniatus*) and San Salvador Island specialists.

| Scaffold | Variant Position | Start | End | Gene |
| --- | --- | --- | --- | --- |
| Introgression with Molluscivore |  |  |  |  |
| c_bro_v1_0_scaf1 | 29209555 | 29160001 | 29250000 | *gga1* |
| c_bro_v1_0_scaf1 | 29241942 | 29195001 | 29250000 | *klf1* |
| c_bro_v1_0_scaf7 | 12326193 | 12300001 | 12375000 | *smek1* |
| c_bro_v1_0_scaf7 | 12628199 | 12610001 | 12670000 | *otof* |
| c_bro_v1_0_scaf24 | 20486354 | 20470001 | 20540000 | *cyp26b1* |
| c_bro_v1_0_scaf33 | 12634285 | 12590001 | 12655000 | *bri3bp, wdr31* |
| c_bro_v1_0_scaf47 | 16145704 | 16110001 | 16195000 | *ttc33* |
| Introgression with Scale-eater |  |  |  |  |
| c_bro_v1_0_scaf5 | 27882801 | 27845001 | 27900000 | *tcf12* |
| c_bro_v1_0_scaf7 | 12604722 | 12555001 | 12620000 | *otof* |
| c_bro_v1_0_scaf11 | 9503186 | 9500001 | 9550000 | *prlh* |
| c_bro_v1_0_scaf11 | 11975348 | 11930001 | 12010000 | *ncoa2* |
| c_bro_v1_0_scaf16 | 32982520 | 32950001 | 33030000 | *crocc* |
| c_bro_v1_0_scaf18 | 28961523 | 28915001 | 28970000 | *itga5* |
| c_bro_v1_0_scaf37 | 8265887 | 8220001 | 8315000 | *mylipa* |
| c_bro_v1_0_scaf43 | 30297117 | 30250001 | 30325000 | *ppp3r1* |
| c_bro_v1_0_scaf53 | 20832687 | 20830001 | 20880000 | *galnt1* |

**Table S13.** Candidate adaptive introgression regions from North Carolina Coast generalists (C. variegatus) and San Salvador Island specialists.

| Scaffold | Variant<br>Position | Start | End | Gene |
| --- | --- | --- | --- | --- |
| Introgression with Scale-eater |  |  |  |  |
| c_bro_v1_0_scaf1 | 28962108 | 28945001 | 28995000 | *notum2,coq7* |
| c_bro_v1_0_scaf1 | 38350857 | 38330001 | 38400000 | *gpr83* |
| c_bro_v1_0_scaf34 | 32388612 | 32380001 | 32440000 | *eya2* |

**Table S14.**
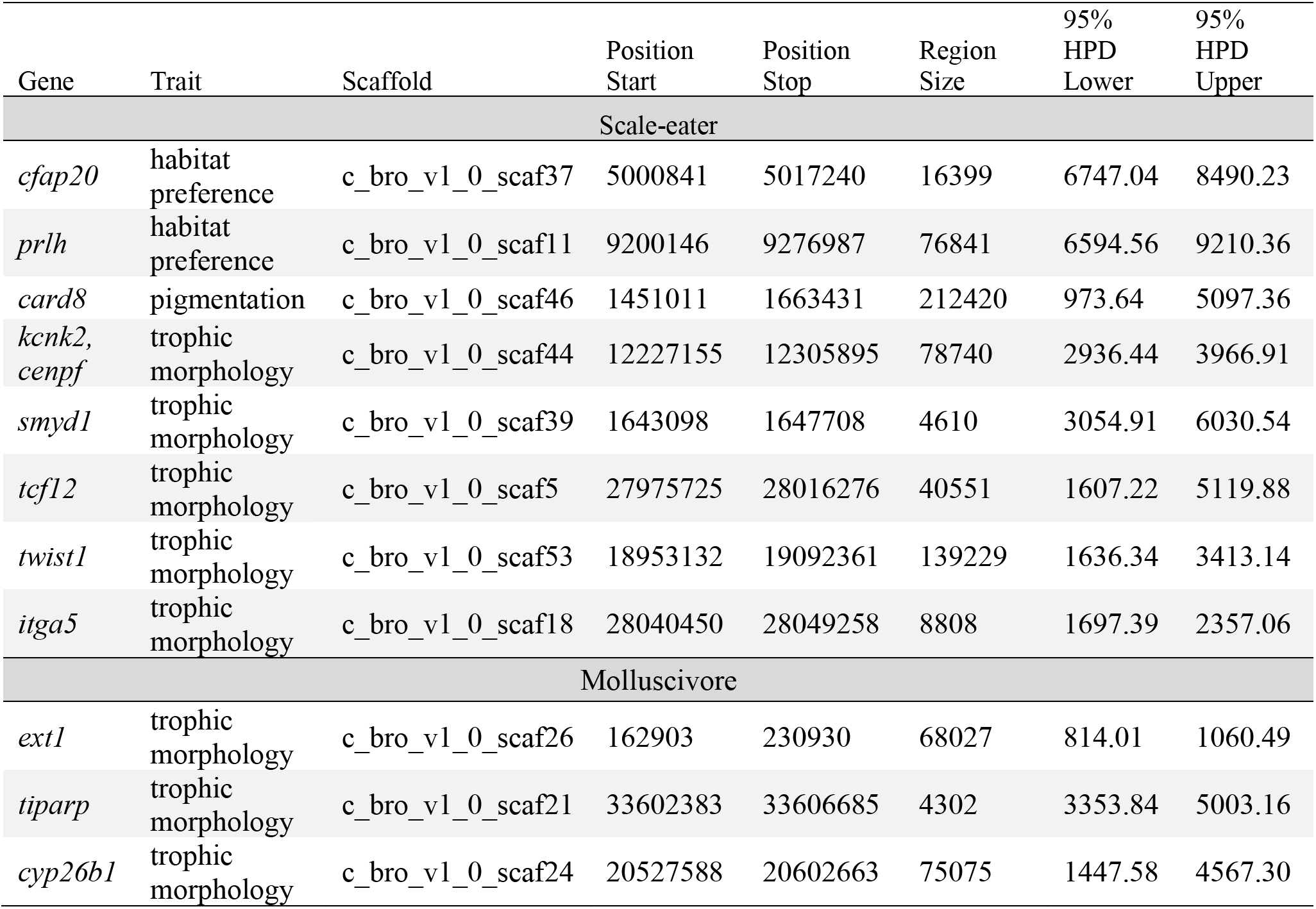
Selective sweep ages on San Salvador Island. 95% high posterior density region of the posterior distribution of sweep ages of focal regions in scale-eater and molluscivore genomes estimated using McSwan (*48*).

**Data S1.**

***Cyprinodon* pupfish sampling information.** The pond/lake names, localities, island, country, and species names, and individual codes of the pupfish individuals used in this study.

**Data S2.**

**San Salvador Island scale-eater candidate variants.** The candidate scale-eater variants that were nearly-fixed (Fst > 0.95) and in a region with a signature of a hard selective sweep and the genes within 20-kb of them.

**Data S3.**

**San Salvador Island molluscivore candidate variants.** The candidate molluscivore variants that were nearly-fixed (Fst > 0.95) and in a region with a signature of a hard selective sweep and the genes within 20-kb of them.

**Data S4.**

**Differentially expressed genes at 2 dpf.** The gene names and *P*-values of genes that were found to be significantly differential expressed (FDR> 0.05) between scale-eaters and mollsucivores at the 2 days post-fertilization larval stage in a previous study (*36*).

**Data S5.**

**Differentially expressed genes at 8 dpf.** The gene names and *P*-values of genes that were found to be significantly differential expressed (FDR> 0.05) between scale-eaters and mollsucivores at the 8 days post-fertilization larval stage in a previous study (*36*).

**Data S6.**

**SSI GWAS trait measurements**. Trait values of standard length, lower oral jaw size, nasal protrusion distance, and caudal fin pigmentation measured across the three species of San Salvador Island radiation to include in a GWAS for candidate variants underlying these traits.

**Data S7**

**Top genomic regions associated with lower oral jaw size in a GWAS of SSI species.** Regions in which all the variants within a 20-kb windows had a summed PIP score that was in the 99^th^ percentile of all summed PIP scores for association with lower oral jaw size across 10 independent runs of Bayesian linear mixed model implemented in GEMMA (*37*).

**Data S8.**

**Top genomic regions associated with lower caudal fin pigmentation in a GWAS of SSI species.** Regions in which all the variants within a 20-kb windows had a summed PIP score that was in the 99^th^ percentile of all summed PIP scores for association with caudal fin pigmentation across 10 independent runs of Bayesian linear mixed model implemented in GEMMA (*37*).

**Data S9.**

Top genomic regions associated with lower maxillary nasal protrusion in a GWAS of SSI species. Regions in which all the variants within a 20-kb windows had a summed PIP score that was in the 99^th^ percentile of all summed PIP scores for association with maxillary nasal protrusion across 10 independent runs of Bayesian linear mixed model implemented in GEMMA.

